# An Optimal Regulation of Fluxes Dictates Microbial Growth In and Out of Steady-State

**DOI:** 10.1101/2022.01.27.477569

**Authors:** Griffin Chure, Jonas Cremer

## Abstract

Effective coordination of cellular processes is critical to ensure the competitive growth of microbial organisms. Pivotal to this coordination is the appropriate partitioning of cellular resources between protein synthesis via translation and the metabolism needed to sustain it. Here, we extend a low-dimensional allocation model to describe the dynamic control of this resource partitioning. At the core of this regulation is the optimal coordination of metabolic and translational fluxes, mechanistically achieved via the perception of charged- and uncharged-tRNA turnover. An extensive comparison with ≈ 60 data sets from *Escherichia coli* establishes this regulatory mechanism’s biological veracity and demonstrates that a remarkably wide range of growth phenomena in and out of steady state can be predicted with quantitative accuracy. This predictive power, achieved with only a few biological parameters, cements the preeminent importance of optimal flux regulation across conditions and establishes low-dimensional allocation models as an ideal physiological framework to interrogate the dynamics of growth, competition, and adaptation in complex and ever-changing environments.

## Introduction

Growth and reproduction is central to life. This is particularly true of microbial organisms where the ability to quickly accumulate biomass is critical for competition in ecologically diverse habitats. Understanding which cellular processes are key in defining growth has thus become a fundamental goal in the field of microbiology. Pioneering physiological and metabolic studies throughout the 20th century laid the groundwork needed to answer this question [1–11], with the extensive characterization of cellular composition across growth conditions at both the elemental [12–14] and molecular [7, 8, 15, 16] levels showing that the dry mass of microbial cells is primarily composed of proteins and RNA. Seminal studies further revealed that the cellular RNA content is strongly correlated with the growth rate [7, 8, 17], an observation which has held for many microbial species [18]. As the majority of RNA is ribosomal, these observations suggested that protein synthesis via ribosomes is a major determinant of biomass accumulation in nutrient replete conditions [19–21]. Given that the cellular processes involved in biosynthesis, particularly those of protein synthesis, are well conserved between species and domains [22–24], these findings have inspired hope that fundamental principles of microbial growth can be found despite the enormous diversity of microbial species and the variety of habitats they occupy.

The past decade has seen a flurry of experimental studies further establishing the importance of protein synthesis in defining growth. Approaches include modern “-omics” techniques with molecular-level resolution [25–38], measurements of many core physiological processes and their coordination [38–46], and the perturbation of major cellular processes like translation [39, 47–49]. Together, these studies advanced a more thorough description of how cells allocate their ribosomes to the synthesis of different proteins depending on their metabolic state and the environmental conditions they encounter, called *ribosomal allocation*. Tied to the experimental studies, different theoretical *ribosomal allocation models* have further been formulated to dissect how ribosomal allocation influences growth [31, 32, 49–56, 56–64]. For example, high-dimensional models have been formulated which simulate hundreds to thousands of biological reactions [51, 59] providing a detailed view of the emergence of distinct internal physiological states and the underlying processes which sustain them. Alternatively, other theoretical considerations follow coarse-grained approaches of moderate dimensionality which group different classes metabolic reactions together and mathematizicing their dynamics [60, 62]. Distinct to those is further an array of extremely low-dimensional models, pioneered by Molenaar *et al*. 2009 [50], which have been developed to describe growth phenomena in varied conditions and physiological limits that rely on only a few parameters [49, 50, 52, 55, 57, 58, 64–66]. A more detailed overview of the different modeling approaches is provided in Appendix 1. Here, we focus particularly on the low-dimensional approaches which have been further linked with phenomenological descriptions [39, 47, 52, 55, 56, 61, 65] many of which took inspiration from a now semnial analysis by Scott *et al* [47] on growth and the adjustment of ribosome content in the presence of translation-inhibiting antibiotics. Several generalizations of the low-dimensional ribosomal allocation have further been proposed [55, 61]. For example, Dourado *et al*. [61] were able to provide solutions of steady-state growth in a generalized manner when several protein classes are included. Though each model is distinct and often tailored to specific environments or growth conditions, their shared minimalism suggests that a single, self-consistent framework can by synthesized to rationalize the fundamental aspects of growth in both static and dynamic environments.

In this work, we build on these low-dimensional allocation models [52, 55, 60, 61, 65] and the results from dozens of experimental studies to synthesize a self-consistent and quantitatively predictive description of resource allocation and growth. At the core of our model is the dynamic reallocation of resources between the translational and metabolic machinery, which is sensitive to the metabolic state of the cell. We demonstrate how “optimal allocation”—meaning, an allocation towards ribosomes which contextually maximizes the steady-state growth rate—emerges when the flux of amino acids through translation to generate new proteins and the flux of uncharged-tRNA through metabolism to provide charged-tRNA required for translation are mutually maximized, given the environmental conditions and corresponding physiological constraints. This regulatory scheme, which we term *flux-parity regulation*, can be mechanistically achieved by a global regulator (e.g., guanosine tetraphosphate, ppGpp, in bacteria) capable of simultaneously measuring the turnover of charged- and uncharged-tRNA pools and routing protein synthesis. The explanatory power of the flux-parity regulation circuit is confirmed by extensive comparison of model predictions with ≈ 60 data sets from *Escherichia coli*, spanning more than half a century of studies using varied methodologies. This comparison demonstrates that a simple argument of flux-sensitive regulation is sufficient to predict bacterial growth phenomena in and out of steady-state and across diverse physiological perturbations. The accuracy of the predictions coupled with the minimalism of the model establishes the optimal regulation and cements the centrality of protein synthesis in defining microbial growth. The mechanistic nature of the theory—predicated on a minimal set of biologically meaningful parameters—provides a low-dimensional framework that can be used to explore complex phenomena at the intersection of physiology, ecology, and evolution without requiring extensive characterization of the myriad biochemical processes which drive them.

### A simple allocation model describes translation-limited growth

We begin by formulating a simplified model of growth which follows the flow of mass from nutrients in the environment to biomass by building upon and extending the general logic of low-dimensional resource allocation models [39, 47, 50, 52, 55]. Specifically, we focus on the accumulation of *protein* biomass, as protein constitutes the majority of microbial dry mass [67, 68] and peptide bond formation commonly accounts for ≈ 80% of the cellular energy budget [33, 69]. Furthermore, low-dimensional allocation models utilize a simplified representation of the proteome where proteins can be categorized into only a few functional classes [48, 50, 52, 54, 61]. In this work, we consider proteins to be either ribosomal (i.e. directly involved in peptide bond formation using charged-tRNAs), metabolic (i.e. enzymes catalyzing synthesis of charged-tRNA molecules from environmental nutrients), or being involved in all other biological processes (e.g. lipid synthesis, DNA replication, and chemotaxis) [47, 48, 50, 52] (Supplementary Figure 1). Simple allocation models further do not distinguish between different cells but only consider the overall turnover of nutrients and biomass. To this end, we explicitly consider a well-mixed batch culture growth as reference scenario where the nutrients are considered to be in abundance. This low-dimensional view of living matter may at first seem like an unfair approximation, ignoring the decades of work interrogating the multitudinous biochemical and biophysical processes of cell-homeostasis and growth [48, 51, 59, 70, 71]. However, at least in nutrient replete conditions, many of these processes appear not to impose a fundamental limit on the rate of growth in the manner that protein synthesis does [33]. In Appendix 2, we discuss this along with other simplifications in more detail.

To understand protein synthesis and biomass growth within the low-dimensional allocation framework, consider the flux diagram (Fig. 1(A), [31, 33, 50, 52, 55]) showing the masses of the three protein classes, precursors which are required for protein synthesis (e.g. charged-tRNA molecules), nutrients which are required for the synthesis of precursors, and the corresponding fluxes through the key biochemical processes (arrows). This diagram emphasizes that growth is autocatalytic in that the synthesis of ribosomes is undertaken by ribosomes which imposes a strict speed limit on growth [33, 72, 73]. While this may imply that the rate of growth monotonically increases with increasing ribosome abundance, it is important to remember that metabolic proteins are needed to supply the ribosomes with the precursors needed to form peptide bonds. Herein lies the crux of ribosomal allocation models: the abundance of ribosomes is constrained by the need to synthesize other proteins and growth is a result of how new protein synthesis is partitioned between ribosomal, metabolic, and other proteins. How is this partitioning determined, and how does it affect growth?

**Figure 1:**
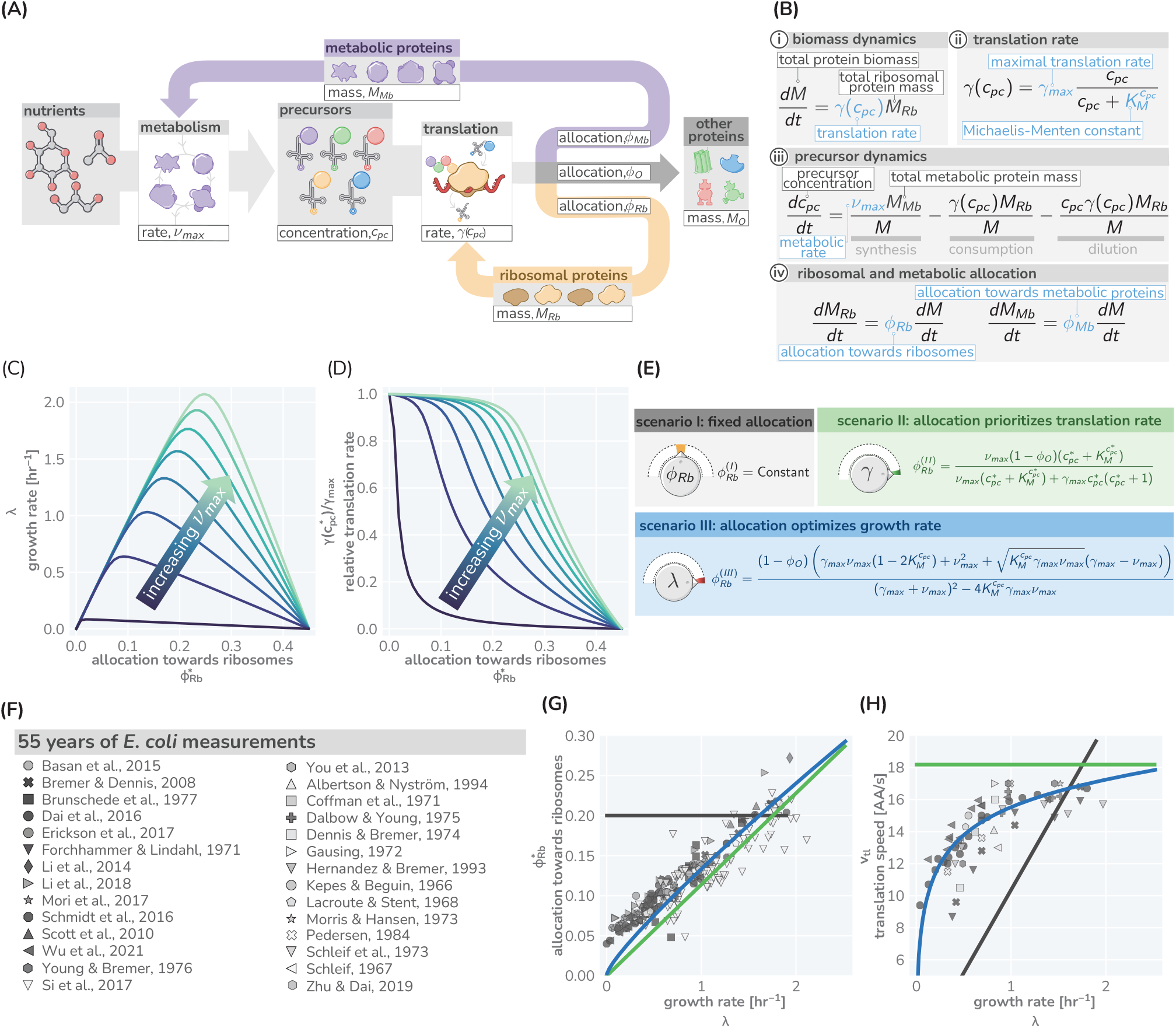
A simple model of ribosomal allocation and hypothetical regulatory strategies. (A) The flow of mass through the self-replicating system. Biomolecules and biosynthetic processes are shown as grey and white boxes, respectively. Nutrients in the environment passed through cellular metabolism to produce “precursor” molecules, here diagrammed as charged-tRNA molecules. These precursors are consumed through the process of translation to produce new protein biomass, either as metabolic proteins (purple arrow), ribosomal proteins (gold arrow), or “other” proteins (gray arrow). The mathematical symbols as used in the simplistic ribosomal allocation model are indicated. (B) Annotated equations of the model with key parameters highlighted in blue. An interactive figure where these equations can be numerically integrated is provided on paper website. The steady-state values of (C) the growth rate *λ* and (D) the relative translation rate 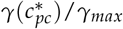, are plotted as functions of the allocation towards ribosomes for different metabolic rates (colored lines). (E) Analytical solutions for candidate scenarios for regulation of ribosomal allocation with fixed allocation, allocation to prioritize translation rate, and allocation to optimal growth rate highlighted in grey, green, and blue respectively. (F) A list of collated data sets of *E. coli* ribosomal allocation and translation speed measurements spanning 55 years of research. Details regarding these sources and method of data collation is provided in Table S1. A comparison of the observations with predicted growth-rate dependence of ribosomal allocation (G) and translation speeds (H) for the three allocation strategies. An interactive version of the panels allowing the free adjustment of parameters is available on the associated paper website (cremerlab.github.io/flux_parity).

To answer these questions, we must understand how these different fluxes interact at a quantitative level and thus must mathematize the biology underlying the boxes and arrows in Fig. 1(A). Taking inspiration from previous models of allocation [47, 50, 52, 55, 61], we enumerate a minimal set of coupled differential equations which captures the flow of mass through metabolism and translation [Fig. 1(C)]. While we present a step-by-step introduction of this model in the Methods, we here focus on a summary of the underlying biological intuition and implications of the approach.

We begin by codifying the assertion that protein synthesis is key in determining growth. The synthesis of new total protein mass *M* depends on the total proteinaceous mass of ribosomes *M*_*Rb*_ present in the system and their corresponding average translation rate *γ* [Fig. 1(B, i)]. As ribosomes rely on precursors to work, it is reasonable to assert that this translation rate must be dependent on the concentration of precursors *c*_*pc*_ such that *γ ≡ γ*(*c*_*pc*_) [52, 55], for which a simple Michaelis-Menten relation is biochemically well motivated [Fig. 1(B, ii)]. The standing precursor concentration *c*_*pc*_ is set by a combination of processes [Fig. 1(B, iii)], namely the production of new precursors through metabolism (synthesis), their degradation through translation (consumption), and their dilution as the total cell volume grows. The synthetic process is driven by the abundance of metabolic proteins *M*_*Mb*_ in the system and the average metabolic rate *ν* at which they convert nutrients into charged-tRNAs. While this rate is in general dependent on the concentration of nutrients in the environment (Supplementary Figure 2 and Methods, we here focus on a growth scenario in which nutrient concentrations are saturating. In such a scenario, metabolism operates at a nutrient-specific maximal metabolic rate *ν ≡ ν*_*max*_. Finally, the relative magnitude of the ribosomal, metabolic, and “other” protein masses is dictated by *φ*_*Rb*_, *φ*_*Mb*_, and *φ*_*O*_, three *allocation parameters* which range between 0 and 1 and follow the constraint *φ*_*Rb*_ + *φ*_*Mb*_ + *φ*_*O*_ = 1 to describe the allocation of total protein synthesis [Fig. 1(B, iv)]. Together, these equations provide a full mathematization of the mass flow diagram shown in Fig. 1 (A).

For constant allocation parameters 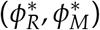 a steady-state regime emerges from this system of differential equation. Particularly, the precursor concentration is stationary in time 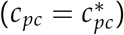, meaning the rate of synthesis is exactly equal to the rate of consumption and dilution. Furthermore, the translation rate 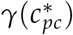 is constant during steady-state growth and the mass-abundances of ribosomes and metabolic proteins are equivalent to the corresponding allocation parameters, e.g. 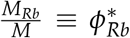. As a consequence, biomass is increasing exponentially 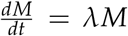, with the growth rate 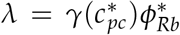. The emergence of a steady state and analytical solutions describing steady growth are further discussed in Supplementary Figures 2 and 3. Notably, dilution is important to obtain a steady state as has been highlighted previously by Giordano *et al*. [55] *and Dourado et al*. [61] *but is often neglected (Appendix 3)*.

Fig. 1 (C) and (D) show how the steady-state growth rate *λ* and translation rate 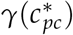 are dependent on the allocation towards ribosomes 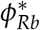. The figures also show the dependence on the metabolic rate *ν*_*max*_ which we here assert to be a proxy for the “quality” of the nutrients in the environment (with increasing *ν*_*max*_, less metabolic proteins are required to obtain the same synthesis of precursors). The non-monotonic dependence of the steady-state growth rate on the ribosome allocation and the metabolic rate poses a critical question: What biological mechanisms determine the allocation towards ribosomes in a particular environment and what criteria must be met for the allocation to ensure efficient growth?

### Different Strategies for Regulation of Allocation Predicts Different Phenomenological Behavior

While cells might employ many different ways to regulate allocation, we here consider three specific allocation scenarios to illustrate the importance of allocation on growth. These candidate scenarios either strictly maintain the total ribosomal content (scenario I), maintain a high rate of translation (scenario II), or optimize the steady-state growth rate (scenario III). We derive analytical solutions for these scenarios (as has been previously performed for scenario III [55, 61]) [Fig. 1(E) and Methods], and ultimately compare these predictions to observations with *Escherichia coli* to show this organisms’ optimal allocation of resources.

The simplest and perhaps most näive regulatory scenario is one in which the allocation towards ribosomes is completely fixed and independent of the environmental conditions. This strategy [scenario I in Fig. 1(E, grey)] represents a locked-in physiological state where a specific constant fraction of all proteins is ribosomal. This imposes a strict speed-limit for growth when all ribosomes are translating close to their maximal rate, 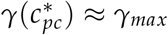. If the fixed allocation is low (for example, 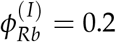, then this speed-limit could be reached at moderate metabolic rates. A more complex regulatory scenario is one in which the allocation towards ribosomes is adjusted to prioritize the translation rate. This strategy [scenario II in Fig. 1(E, green)] requires that the ribosomal allocation is adjusted such that a constant internal concentration of precursors 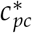 is maintained across environmental conditions, irrespective of the metabolic rate. In the case where this standing precursor concentration is large 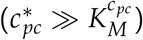, all ribosomes will be translating close to their maximal rate.

The third and final regulatory scenario is one in which the allocation towards ribosomes is adjusted such that the steady-state growth rate is maximized. The analytical solution which describes this scenario [scenario III in Fig. 1(E)] resembles previous analytical solutions by Giordano et al and Dourado et al [55, 61]. More illustratively, the strategy can be thought of as one in which the allocation towards ribosomes is tuned across conditions such that the observed growth rate rests at the peak of the curves in Fig. 1(E). Notably, this does not imply that the translation rate is constantly high across conditions (as in scenario II). Rather, the translation rate is also adjusted and approaches its maximal value *γ*_*max*_ only in very rich conditions (high metabolic rates). All allocation scenarios and their consequence on growth are discussed in further detail in Supplementary Figure 4 and the corresponding interactive figure on the paper website.

### *E. coli* Regulates Its Ribosome Content to Optimize Growth

Thus far, our modeling of microbial growth has remained “organism agnostic” without pinning parameters to the specifics of any one microbe’s physiology. To probe the predictive power of this simple allocation model and test the plausibility of the three different strategies for regulation of ribosomal allocation, we performed a systematic and comprehensive survey of data from a vast array quantitative studies of the well characterized bacterium *E. coli*. This analysis, consisting of 26 studies spanning 55 years of research (listed in Table S2) using varied experimental methods, goes well beyond previous attempts to compare allocation models to data [47, 48, 53–55, 57, 60, 60–63, 65].

These data, shown in Fig. 1(F - H, markers), present a highly-consistent view of *E. coli* physiology where the allocation towards ribosomes (equivalent to ribosomal mass fraction in steady-state balanced growth) and the translation rate demonstrate a strong dependence on the steady-state growth rate in different carbon sources. The pronounced correlation between the allocation towards ribosomes and the steady-state growth rate immediately rules out scenario I, where allocation is constant, as a plausible regulatory strategy used by *E. coli*, regardless of its precise value. Similarly, the presence of a dependence of the translation speed on the growth rate rules out scenario II, where the translation rate is prioritized across growth rates and maintained at a constant value. The observed phenomenology for both the ribosomal allocation *and* the translation speed is only consistent with the logic of regulatory scenario III where the allocation towards ribosomes is tuned to optimize growth rate.

This logic is quantitatively confirmed when we compute the predicted dependencies of these quantities on the steady-state growth rate for the three scenarios diagrammed in Fig. 1(E) based on literature values for key parameters (outlined in Table S1). Deviations from the prediction for scenario III are only evident for the ribosomal content at very slow *steady* growth (*λ ≤* 0.5 hr^−1^) which are hardly observed in any ecologically relevant conditions and can be attributed to additional biological and experimental factors, including protein degradation [74], ribosome inactivation [39, 42], and cultures which have not yet reached steady state, factors we discuss in Appendix 4.

Importantly, the agreement between theory and observations works with a minimal number of parameters and does not require the inclusion of fitting parameters. All fixed model parameters such as the maximum translation rate *γ*_*max*_ and the Michaelis-Menten constant for translation 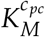, have distinct biological meaning and can be either directly measured or inferred from data (a list is provided in Table S1). Furthermore, we discuss the necessity of other parameters such as the “other protein sector” *φ*_*O*_ (Appendix 5), its degeneracy with the maximum metabolic rate *ν*_*max*_, and inclusion of ribosome inactivation and minimal ribosome content (Appendix 4, 6). We furthermore provide an interactive figure on the paper website where the parametric sensitivity of these regulatory scenarios and the agreement/disagreement with data can be directly explored. Notably there is no combination of parameter values that would allow scenario I or II to adequately describe both the ribosomal allocation and translation speed as a function of growth rate. Notably, our findings are in line with a recent higher-dimensional modeling study [60] which, based on the optimization of a reaction network with > 200 components, rationalized the variation in translation speed with growth as a manifestation of efficient protein synthesis. Together, these results confirm that scenario III can accurately describe observations over a very broad range of conditions, in strong support of the popular but often questioned presumption that *E. coli* optimally tunes its ribosomal content to promote fast growth [49, 55, 65].

In Appendix 7, we present a similar analysis for *Saccharomyces cerevisiae* which, in line with previous studies [34–37], suggests that this eukaryote likely follows a similar optimal allocation strategy, although data for ribosomal content and the translation reate is scarce. The strong correlation between ribosome content and growth rate has further been reported for other microbial organisms in line with an optimal allocation [18, 38, 45, 75], though the absence of translation rate measurements precludes confirmation. An interesting exception is the methanogenic archaeon *Methanococcus maripaludis* which appears to maintain constant allocation, in agreement with scenario I [76]. The presented analysis thus suggests that *E. coli* and possibly many other microbes closely follow an optimal ribosome allocation behavior to support efficient growth. Moreover, the good agreement between experiments and data establishes that a simple low-dimensional allocation model can describe growth with notable quantitative accuracy. However, this begs the question: how do cells coordinate their complex machinery to ensure optimal allocation?

### Optimal Allocation Results From a Mutual Maximization of Translational and Metabolic Flux

To optimize the steady-state growth rate, cells must have some means of coordinating the flow of mass through metabolism and protein synthesis. In the ribosomal allocation model, this reduces to a regulatory mechanism in which the allocation parameters (*φ*_*Rb*_ and *φ*_*Mb*_) are dynamically adjusted such that the metabolic flux to provide new precursors (*νφ*_*Mb*_) and translational flux to make new proteins (*γφ*_*Rb*_, equivalent to the steady-state growth rate *λ*) are not only equal, but are mutually maximized. Such regulation therefore requires a mechanism by which both the metabolic and translational flux can be simultaneously sensed.

Thus far, we have referred to the end-product of metabolism as ambiguous “precursors” which are used by ribosomes to create new proteins. In reality, these precursors are tRNAs charged with their cognate amino acids. One can think of metabolism as a two-step process where (i) an amino acid is synthesized from environmental nutrients and (ii) an amino acid is attached to the appropriate uncharged-tRNA. As we assume that nutrients are in excess in the environment, we make the approximation that the metabolic rate *ν* is dependent solely on the uncharged-tRNA concentration *ν*(tRNA^*u*^). This enforces some level of regulation of metabolism; if the uncharged tRNA concentration is too low, the rate of metabolism slows and does not add to the already large pool of charged tRNA. But when charged-tRNA is available, translation occurs at a rate *γ*(tRNA^*c*^), forming new protein biomass and converting a charged-tRNA back to an uncharged state. This process is shown by grey arrows in Fig. 2(A).

**Figure 2:**
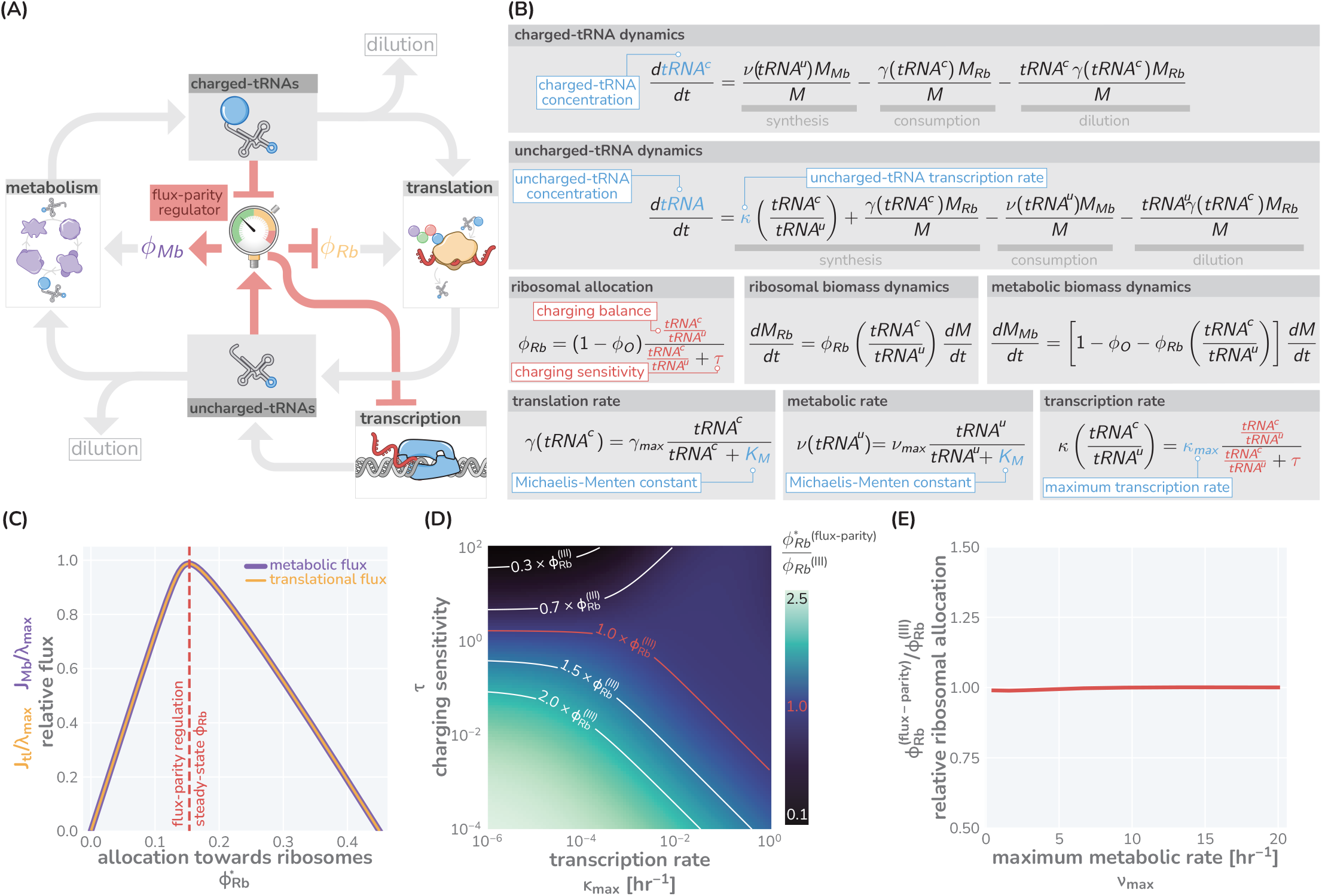
The regulation of ribosome allocation via a flux-sensing mechanism. (A) A circuit diagram of interactions between metabolic and translational fluxes with flux-parity regulatory connections highlighted in red. The fluxes are connected via a positive feedback loop through the generation of mutual starting materials (uncharged- or charged-tRNAs, respectively). The rates of each flux exhibit semi-autoregulatory behavior in that flux through each process reduces the standing pool of tRNAs. (B) The governing dynamics of the flux-parity regulatory circuit with key parameters highlighted in blue and flux-parity regulatory components highlighted in red. (C) The steady-state meabolic (purple) and translational (gold) fluxes plotted as a function of the ribosomal allocation under the simple allocation model. Vertical red line indicates the steady-state solution of the flux-parity model under physiological parameter regimes. (D) The steady-state allocation towards ribosomes emerging from flux-parity regulation 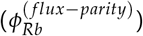 relative to the optimal allocation of the simplistic model 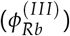 is shown across different parameter regimes for the charging sensitivity *τ* and the uncharged-tRNA transcription rate *κ*. Red contour demonstrate the plane of parameter space where the flux-parity regulatory circuit exactly matches the result of optimal allocation. (E) The relative allocation towards ribosomes 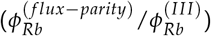 under physiological parameter regimes plotted as a function of the maximal metabolic rate, *ν*_*max*_.

To describe the state-dependent adjustment of the allocation parameters (*φ*_*Rb*_ and *φ*_*Mb*_), we further include in this feedback loop a regulatory system we term a “flux-parity regulator” [Fig. 2(A), red], which controls the allocation parameters in response to relative changes in the concentrations of the two tRNA species. Together, the arrows in Fig. 2 represents a more fine-grained view of a proteinaceous self replicating system, yet maintains much of the structural minimalism of the simple ribosomal allocation model without requiring explicit consideration of different types of amino acids [65], inclusion of their myriad synthesis pathways [60], or reliance on observed phenomenology [42].

The boxes and arrows of Fig. 2(A) can be mathematized to arrive at a handful of ordinary differential equations [Fig. 2(B)] structurally similar to those in Fig. 1(B). At the center of this model is the ansatz that the ribosomal allocation *φ*_*Rb*_ is dependent on the ratio of charged- and uncharged-tRNA pools and has the form

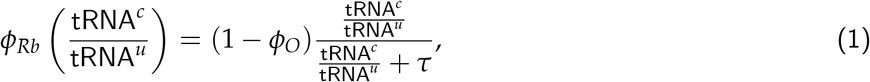

where the ratio 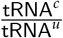 represents the “charging balance” of the tRNA and *τ* is a dimensionless “sensitivity parameter” which defines the charging balance at which the allocation towards ribosomes is half-maximal. Additionally, we make the assertion that the synthesis rate of new uncharged-tRNA via transcription *κ* is coregulated with ribosomal proteins [77, 78] and has a similar form of

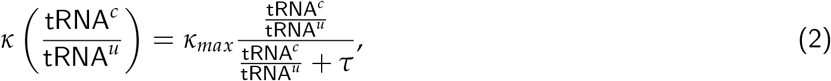

where *κ*_*max*_ represents the maximal rate of tRNA transcription relative to the total biomass.

Numerical integration of this system of equations reveals that the flux-parity regulation is capable of optimizing the allocation towards ribosomes, *φ*_*Rb*_, such that the metabolic and translation fluxes are mutually maximized [Fig. 2(C)], thus achieving optimal allocation. Importantly, the optimal behavior inherent to this regulatory mechanism can be attained across a wide range of parameter values for the charging sensitivity *τ* and the transcription rate *κ*_*max*_, the two key parameters of flux-parity regulation [Fig. 2(D)]. Moreover, the emergent optimal behavior of this regulatory scheme occurs across conditions without the need for any fine-tuning between the flux-parity parameters and other parameters. For example, the control of allocation via the flux parity regulation matches the optimal allocation (scenario III above) when varying the metabolic rate *ν*_*max*_ [Fig. 2(E) and Appendix 8].

The theoretical analysis presented in Fig. 2 suggests that a flux-parity regulatory mechanism may be a simple way to ensure optimal ribosomal allocation that is robust to variation in the key model parameters. To test if such a scheme may be implemented in *E. coli*, we compared the behavior of the steady-state flux-parity regulatory circuit within physiological parameter regimes to steady-state measurements of ribosomal allocation and the translation rate as a function of the growth rate [Fig. 3(A & B)]. Remarkably, the predicted steady-state behavior of the flux-parity regulatory circuit describes the observed data with the same quantitative accuracy as the optimal behavior defined by scenario III, as indicated by the overlapping red and blue lines, respectively.

**Figure 3:**
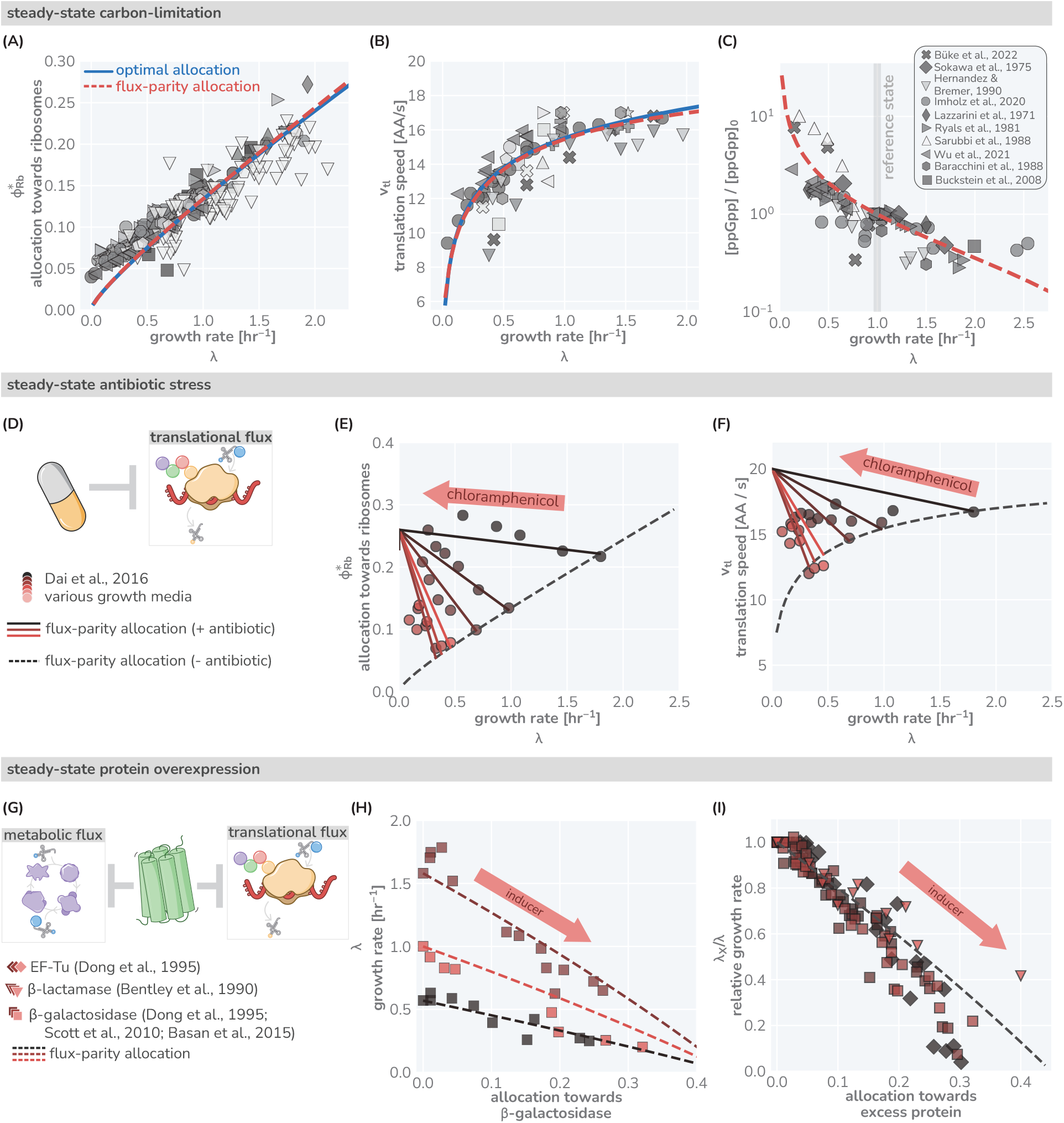
The predictive power of flux-parity regulation in steady-state. Measurements of the (A) ribosomal allocation and the (B) translation rate are plotted alongside the steady-state behavior of the flux-parity regulatory circuit (red dashed line) and the optimal behavior of scenario III (solid blue line). Points and markers are the same as those shown in Fig. 1(F). (C) Measurements of intracellular ppGpp concentrations relative to a reference condition (*λ*_0_ ≈ 1 hr^−1^) are plotted as a function of growth rate alongside the prediction emergent from the flux-parity regulatory circuit (red dashed line). (D-F) Inhibition of ribosome activity via antibiotic modeled repression of translational flux. Plots show comparison with data for different media (red shades) with the flux-parity model predictions (dashed lines). (G-I) Inhibition of metabolic and translational fluxes through excess gene expression. (H) shows data where *β*-galactosidase is expressed at different levels. Different shades of red correspond to different growth media. Right-hand panel shows collapse of the growth rates of overexpression of *β*-galactosidase (squares), *β*-lactamase (inverted triangles), and EF-Tu (diamonds) relative to the wild-type growth rate in different media conditions. The same set of model parameters listed in Table S2 has been used to generate the predictions.

While the flux-parity regulation scheme appears to accurately describe the behavior of *E. coli*, how are metabolic and translational fluxes sensed at a mechanistic level? Many bacteria, including *E. coli*, utilize the small molecule guanosine tetraphosphate (ppGpp) as a molecular indicator of amino acid limitation and has been experimentally shown to regulate ribosomal, metabolic, and tRNA genes [79–82]. Mechanistically, ppGpp levels are enzymatically controlled depending on the metabolic state of the cell, with synthesis being triggered upon binding of an uncharged-tRNA into an actively translating ribosome. While many molecular details of this regulation remain unclear [42, 79–81], the behavior of ppGpp meets all of the criteria of a flux-parity regulator. Rather than explicitly mathematicizing the biochemical dynamics of ppGpp synthesis and degradation, as has been undertaken previously [42, 55, 65], we model the concentration of ppGpp being inversely proportional to the charging balance,

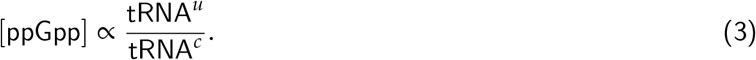

This ratio, mathematically equivalent to the odds of a ribosome binding an uncharged-tRNA relative to binding a charged-tRNA, is one example of a biochemically-motivated ansatz that can be considered (Methods) and provides a relative measure of the metabolic and translational fluxes.

With this approach, the amount of ppGpp present at low growth rates, and therefore low ribosomal allocation, should be significantly larger than at fast growth rates where ribosomal allocation is larger and charged-tRNA are in abundant supply. While our model cannot make predictions of the *absolute* ppGpp concentration, we can compute the *relative* ppGpp concentration to a reference state [ppGpp]_0_ as

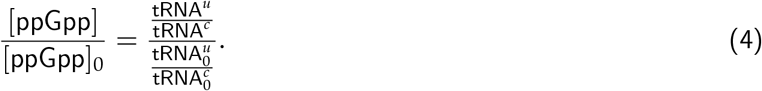

To test this, we compiled and rescaled ppGpp measurements of *E. coli* across a range of growth rates from various literature sources [Fig. 3(C)]. The quantitative agreement between the scaling predicted by Eq. 4 and the experimental measurements strongly suggests that ppGpp assumes the role of a flux-sensor and enforces optimal allocation through the discussed flux-parity mechanism.

### The Flux-Parity Allocation Model Predicts E. coli Growth Behavior In and Out Of Steady-State

We find that the flux-parity allocation model is extremely versatile and allows us to quantitatively describe aspects of microbial growth in and out of steady-state and under various physiological stresses and external perturbations with the same core set of parameter. Here, we demonstrate this versatility by comparing predictions to data for four particular examples *using the same self-consistent set of parameters we have used thus far* (Table S1). First, we examine the influence of translation-targeting antibiotics like chloramphenicol [Fig. 3(D)] on steady-state growth in different growth media [39, 47]. By incorporating a mathematical description of ribosome inactivation via binding to chloramphenicol (described in Methods), we find that the flux-parity allocation model quantitatively predicts the change in steady-state growth and ribosomal content with increasing chloramphenicol concentration [Fig. 3(E, red shades)]. Furthermore, the effect on the translation speed is qualitatively captured [Fig. 3(F, red shades)]. The ability of the flux-parity allocation model to describe these effects without readjustment of the model and its core parameters is notable and provides a mechanistic rationale for previously established phenomenological relations [39, 47].

As a second perturbation, we consider the burden of excess protein synthesis by examining the expression of synthetic genes [Fig. 3(G)]. A decrease in growth rate results when cells are forced to synthesize different amounts of the lactose cleaving enzyme *β*-galactosidase in different media lacking lactose [Fig. 3(H, red shades)]. The flux-parity allocation model (dashed lines) quantitatively predicts the change in growth rate with the measured fraction of *β*-galactosidase without further fitting (Methods). The trends for different media (red shades) quantitatively collapse onto a single line [Fig. 3(I)] when comparing changes in relative growth rates, a relation which is also captured by the model (dashed black line) and is independent of the overexpressed protein (symbols). This collapse, whose functional form is derived in the Methods, demonstrates that the flux-parity allocation model is able to describe excess protein synthesis in general, rather than at molecule- or media-specific level.

As the flux-parity regulatory circuit responds to changes in the metabolic and translational fluxes, it can be used to explore behavior in changing conditions. Consider a configuration where the starting conditions of a culture are tuned such that the ribosomal allocation *φ*_*Rb*_, the tRNA charging balance tRNA^*c*^/tRNA^*u*^, and the ribosome content *M*_*Rb*_/*M* are set to be above or below the appropriate level for steady-state growth in the environment [Fig. 4(A)]. As the culture grows, the observed ribosomal content *M*_*Rb*_/*M* is steadily adjusted until the steady-state level is met where it directly matches the optimal allocation [Fig. 4(B)]. This adaptation of the ribosomal content is controlled by dynamic adjustment of the allocation parameters via the flux-parity regulatory circuit [Fig. 4(C)]. To further test the flux-parity allocation model, we examine how accurately this system can predict growth behavior under nutritional shifts [Fig. 4(D-F)]and the entry to starvation [Fig. 4(G-I)].

**Figure 4:**
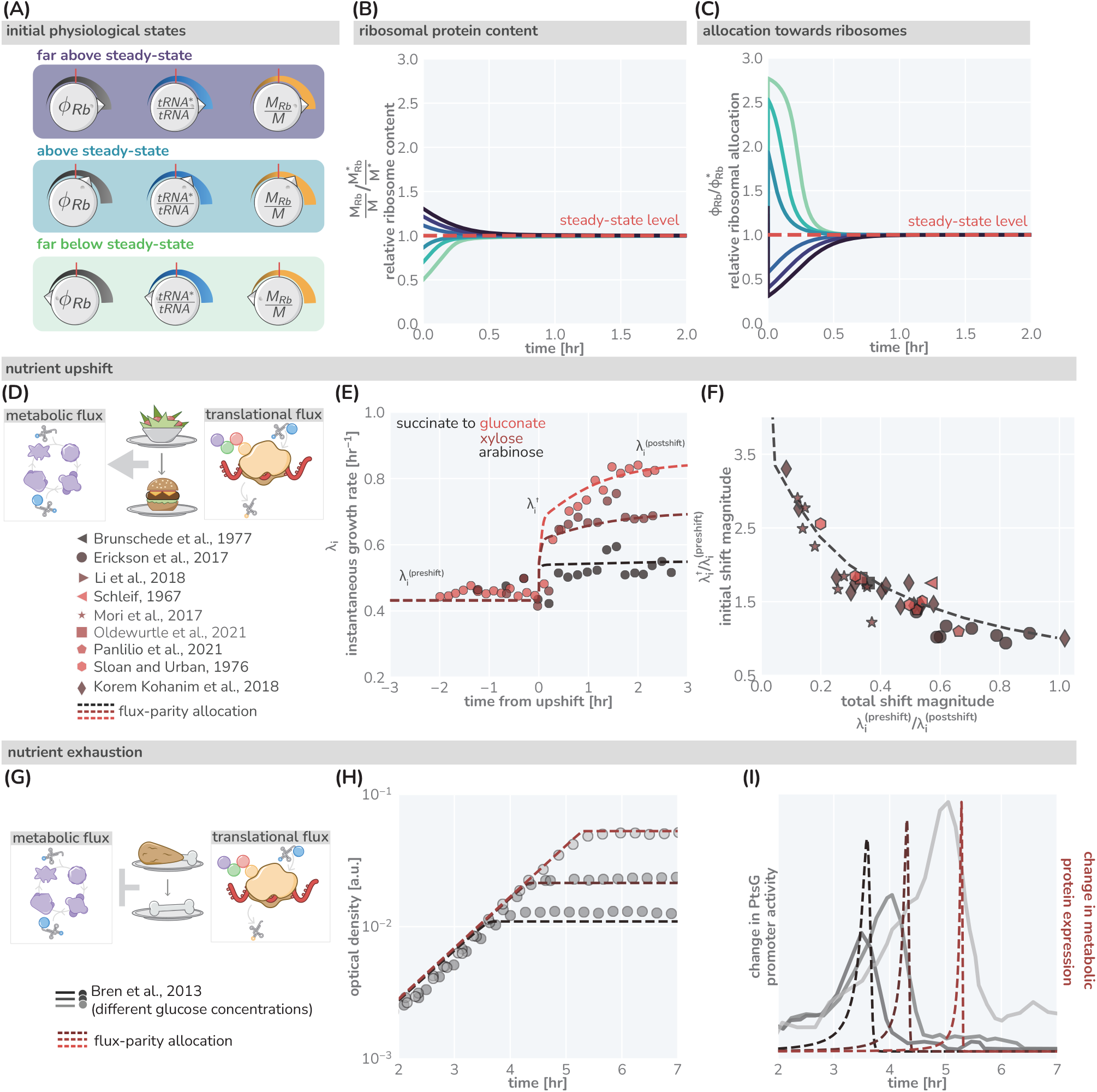
The predictive power of flux-parity regulation out of steady-state. (A) Hypothetical initial configurations of model parameters and variables before begining numerical integration. (B) The equilibration of the ribosomal protein content *(M*_*Rb*_ /*M*). (C) Dynamic adjustment of the ribosomal allocation parameter in response to the new environment. Green and purple colored lines correspond to the initial conditions of the culture from well above to well below the steady-state values, respectively. Dashed red line indicates the steady-state solution. (D-E) Nutrient upshifts with increased metabolic flux. (E) The instantaneous growth rate *λ*_*i*_ for shifts from succinate to gluconate (bright red), xylose (dark red), or arabinose (black) [57]. Collapse of instantaneous growth rate measurements immediately after the shift (relative to the preshift-growth rate) as a function of the total shift magnitude. (G-I) Exhaustion of nutrients in the environment yields a decrease in the metabolic flux, promoting expression of more metabolic proteins. (H) Growth curve measurements in media with different starting concentrations of glucose (0.22 mM, 0.44 mM, and 1.1 mM glucose from light to dark, respectively) overlaid with flux-parity predictions. (I) The change in total metabolic protein synthesis in the flux-parity model (dashed lines) overlaid with the change in expression of a fluorescent reporter from a PtsG promoter (solid lines).

We first consider a nutrient shift where externally-supplied low-quality nutrients are instantaneously exchanged with rich nutrients. Fig. 4(E) shows three examples of such nutritional upshifts (markers), all of which are well described by the flux-parity allocation theory (dashed lines). The precise values of the growth rates before, during, and after the shift will depend on the specific carbon sources involved. However, by relating the growth rates before and immediately after the shift to the total shift magnitude (as shown in Ref. [58]), one can collapse a large collection of data onto a single curve [Fig. 4(F, markers)]. The collapse emerges naturally from the model (dashed-line) when decomposing the metabolic sector into needed and non-needed components (Methods), demonstrating that the flux-parity allocation model is able to quantitatively describe nutritional upshifts at a fundamental level.

Finally, we consider the growth dynamics during the onset of starvation, another non steady-state phenomenon [Fig. 4(G-I)]. Fig. 4(H) shows the growth of batch cultures where glucose is provided as the sole carbon source in different limiting concentrations [83] (markers). The cessation of growth coincides with a rapid, ppGpp-mediated increase in expression of metabolic proteins [79, 84]. Bren *et al*. [83] demonstrated that expression from a glucose-specific metabolic promoter (PtsG) rapidly, yet temporarily, increases with the peak occurring at the moment where growth abruptly stops [Fig. 4(I, solid grey lines)]. The flux-parity allocation model again predicts this behavior [Fig. 4(I, red lines)] without additional fitting (Methods), cementing the ability of the model to describe growth far from steady-state.

## Discussion

Microbial growth results from the orchestration of an astoundingly diverse set of biochemical reactions mediated by thousands of protein species. Despite this enormous complexity, experimental and theoretical studies alike have shown that many growth phenotypes can be captured by relatively simple correlations and models which incorporate only a handful of parameters [7, 39, 47, 50, 52, 55, 57, 58, 65]. Through re-examination of these works, we relax commonly invoked approximations and assumptions, include a generalized description of global regulation, and integrate an extensive comparison with data to establish a self-consistent, low-dimensional model of protein synthesis that is capable of quantitatively describing complex growth behaviors in and out of steady-state.

Growth emerges as in previous allocation models [47, 50, 55] as a consequence of protein synthesis and the allocation of ribosome activity towards (i) making new ribosomes, (ii) making the metabolic proteins which sustain the precursors ribosomes require to translate, and (iii) making other proteins cells require to operate. An *optimal allocation* which yields the fastest growth in a given condition is reached when the synthesis of precursors (metabolic flux) and the consumption of precursors (translational flux) are mutually maximized, a process we term *flux-parity regulation*. We analyze how such regulation can be mechanistically achieved by the relative sensing of charged- and uncharged-tRNA via the abundance of a global regulator (such as ppGpp) which diametrically affects the expression of ribosomal and metabolic genes. Through extensive comparison with 61 data sets from 46 studies, we show that the flux-parity model predicts the fundamental growth behavior of *E. coli* with quantitative accuracy. Beyond the impeccable description of the growth-rate dependent ribosomal content and translation speed across various carbon sources, the flux-parity model quantitatively captures phenomena out of steady-state (including nutrient upshifts and response to starvation) and under externally applied physiological perturbations (such as antibiotic stress or expression of synthetic genes). Notably, the broad agreement across data sets is obtained using a single core parameter set which does not require any adjustment from one scenario to the next. As such, the flux-parity model predicts the microbial “growth laws”, providing a mechanistic explanation for previous phenomenological models formulated to understand them [47, 50, 52]. The finding that these predictions hold so well despite the overwhelmingly complex nature of the cell further highlights that biological systems are not irreducibly complex but can be distilled to a small number of fundamental components sufficient to capture the core behavior of the system.

As proteins commonly account for the majority of biomass in microbial organisms and the core processes of protein synthesis are universally conserved among them, it is likely that protein synthesis is a fundamental growth constraint across many organisms. Accordingly, flux-parity regulation may be a very general scheme which ensures the efficient coordination of metabolic and translational fluxes across many microbial organisms. And as our modeling approach is organism agnostic, it should be transferable to a variety of microbes growing in nutrient-replete conditions. Indeed, other organisms including *S. cerevisiae* (Appendix 7) exhibit a strict interdependence between growth rate and ribosome content [18], as is predicted by the flux-parity model. However, more quantitative data on ribosomal content, translation speeds, upshift dynamics, and more need to be acquired to fully examine the commonality of flux-parity regulation in the microbial world.

A common interpretation of previous allocation models is that cells maximize their growth rate in whatever conditions they encounter [49, 65]. Rather, we believe flux-parity regulation only ensures optimal coordination between metabolic and translational fluxes. It does not imply that the growth rate itself is maximized or directly sensed. In particular, the flux-parity model does not assume that the pool of metabolic proteins is tailored to maximize the metabolic flux and thus growth in the encountered conditions. This is in agreement with an expanding body of evidence which shows that microbes frequently synthesize metabolic and other proteins which are not directly needed in the encountered condition and thus impede growth. *E. coli*, for example, synthesizes a plethora of different transport proteins when exposed to poor growth conditions even if the corresponding substrates are not available, collectively occupying a significant portion of the proteome [27, 33, 48, 64]. Accordingly, it has been observed that cells stop synthesizing these proteins when evolving over many generations in the absence of those sugars [85, 86].

But why, then, do we observe an optimal allocation between metabolic and ribosomal proteins when the pool of metabolic proteins itself shows this apparent non-optimal behavior? We posit here that both behaviors emerge from the adaptation to fluctuating conditions: in contrast to the well-defined static conditions of laboratory experiments, the continuous ebb and flow of nutrients in natural environments precludes any sense of stability. Accordingly, the machinery of the cell should be predominantly adapted to best cope with the fluctuating conditions microbial organisms encounter in their natural habitats. A complex regulation of metabolic proteins is thus expected, including for example, the diverse expression of nutrient transporters which promote growth in anticipated conditions, rather than synthesizing only those specific to nutrients that are present in the moment [64].

However, in those fluctuating conditions, flux-parity regulation promotes rapid growth. To illustrate this point, we consider again a nutrient upshift in which there is an instantaneous improvement in the nutrient conditions. We compare the predicted response via flux-parity [Fig. 5 (B, red box)] with that predicted by a simpler step-wise regulation where the allocation solely depends on the environmental condition (and not the internal fluxes) and immediately adjusts to the new steady value at the moment of the shift [Fig. 5 (B, blue box)]. The dynamic reallocation by flux-parity facilitates a sharp increase in the allocation towards ribosomes [Fig. 5(C)], resulting in a rapid increase in instantaneous growth rate compared to the step-wise reallocation mechanism [Fig. 5(D)], suggesting that flux-parity is advantageous in fluctuating environments. As its regulation solely depends on the internal state of the cell (particularly, the relative abundance of charged-to uncharged-tRNA) it holds independently of the encountered conditions. This stands in contrast to the regulation of metabolic proteins, where both the external and internal states dictate what genes are expressed. As a result, optimal coordination between metabolic and translational fluxes occurs ubiquitously across conditions and not only in those that occur in natural habitats and drive adaptation. These broader conditions include steady-state growth within the laboratory, with the “growth laws” observed under those conditions emerging as a serendipitous consequence.

**Figure 5:**
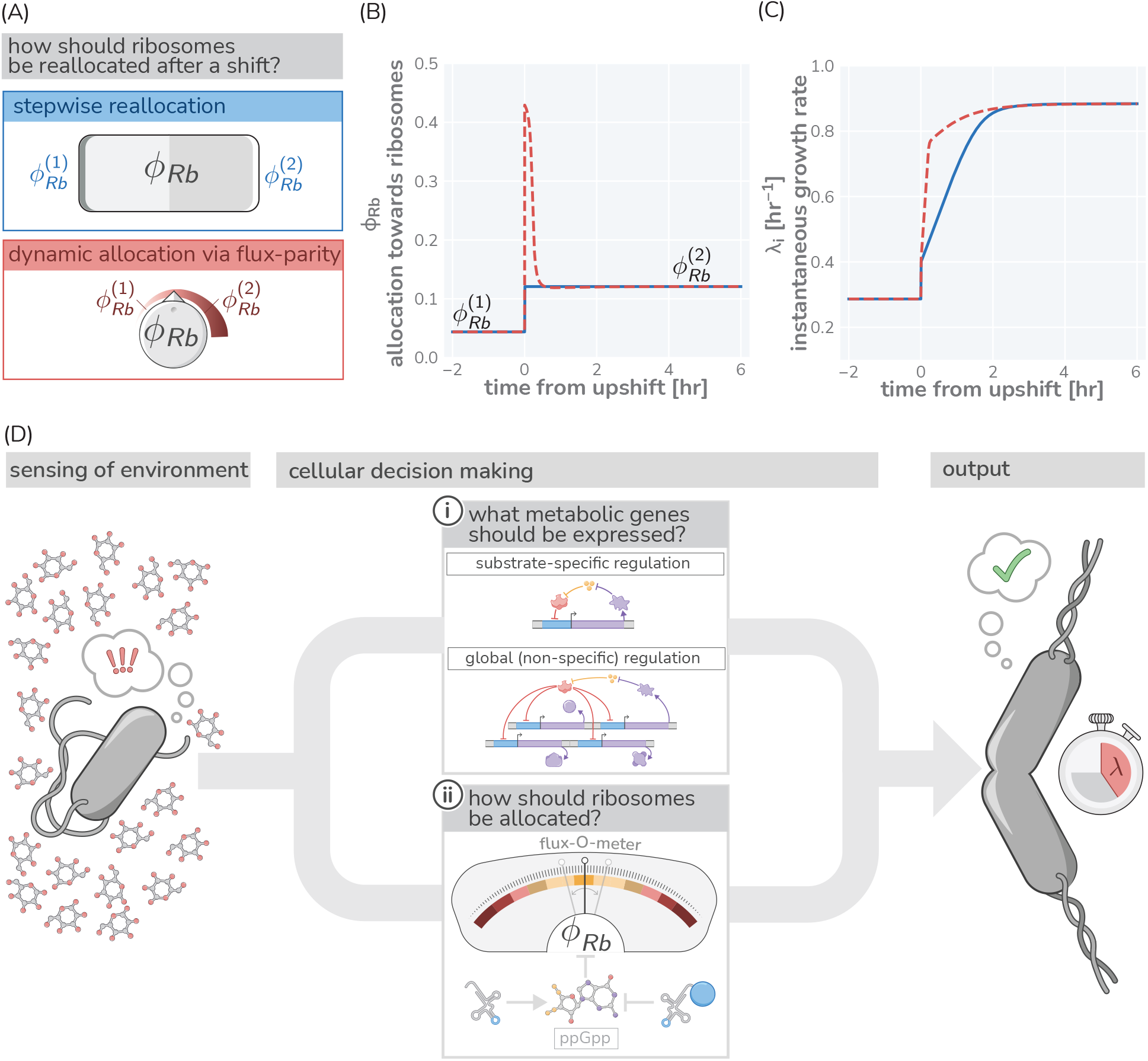
Flux parity allocation as a strategy to adapt to fluctuating conditions. (A) Ribosome reallocation strategies upon a nutrient upshift. After a nutrient upshift, cells either dynamically reallocate their ribosomes given flux-parity regulation (top, red) or they undergo stepwise reallocation from one steady-state value to the next (bottom, blue). (B) The allocation dynamics for both strategies in response to a nutrient upshift. (C) The instantaneous growth rate for both strategies over the course of the shift. Dashed red and solid blue lines correspond to model predictions for optimal allocation and flux-parity regulation, respectively. (D) Cellular decision making in fluctuating environments. Upon sensing features of the environment, cells undergo a two-component decision making protocol defining what metabolic genes should be expressed (top) and how the allocation towards ribosomes should be adjusted to maintain flux-parity. The combination of these processes yield an increase of biomass at a given characteristic growth rate.

In summary, we view the process of cellular decision making as having two major components [Fig. 5(D)]: (i) determining what metabolic genes should be expressed given the environmental and physiological state and (ii) determining how ribosomes should be allocated given the metabolic and translational fluxes. Flux-parity regulation can explain the latter but many details of the former remain enigmatic. Additional studies are thus required to understand how the regulation of metabolic genes depends on encountered conditions and how it is shaped by adaptation to specific habitats. However, the ability of this theory to predict complex phenotypes across scales suggests that it can also act as a basis to answer these questions, and thereby galvanize an integrative understanding of microbial life connecting physiology, ecology, and evolution.

## Methods

### Formulating the allocation model

Here we present a step-by-step derivation of the the low-dimensional allocation model we use to describe bacterial growth. We provide additional biological motivation for its construction and highlight the different assumptions and simplifications invoked. To maintain consistency with the literature, we largely follow the notational scheme introduced by Scott *et al* [52] and define each symbol as it is introduced.

### Synthesis of Proteins

The rate of protein synthesis is determined by two quantities – the total number of ribosomes *N*_*Rb*_ and the speed *v*_*tl*_ at which they are translating. The latter depends on the concentration of precursors needed for peptide bond formation, such as tRNAs, free amino acids, and energy sources like ATP and GTP. Taking the speed *v*_*tl*_ as a function of the concentration of the collective precursor pool *c*_*pc*_, the increase in protein biomass *M* follows as

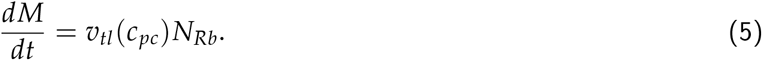

There exists a maximal speed at which ribosomes can operate, 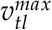, that is reached under optimal conditions when precursors are highly abundant, in *E. coli* approximately 20 amino acids (AA) / second (s) [87]. Conversely, the translation speed falls when precursor concentrations *c*_*pc*_ get sufficiently small. Simple biochemical considerations support a Michaelis-Menten relation [33, 88, 89] as good approximation of this behavior with the specific form

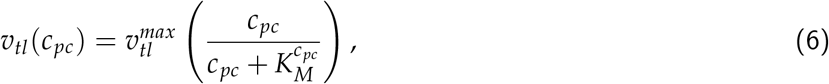

where 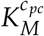 is a Michaelis-Menten constant with the maximum speed 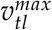 only observed for 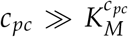. The number of ribosomes *N*_*Rb*_ can be approximated given knowledge of the total mass of ribosomal proteins *M*_*Rb*_ and the proteinaceous mass of a single ribosome *m*_*Rb*_ via *N*_*Rb*_ *≈ M*_*Rb*_/*m*_*Rb*_(more details in Appendix 9). The increase in protein biomass (Eq. 5) is thus

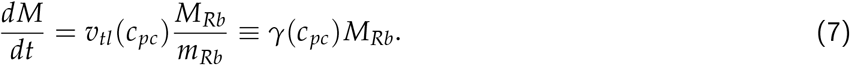

The *translation rate γ*(*c*_*pc*_) *≡ v*_*tl*_(*c*_*pc*_)/*m*_*Rb*_ describes the rate at which ribosomes generate new protein.

The maximal translation rate 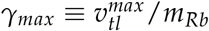 imposes a firm upper limit [33, 72, 73] of how rapidly biomass can accumulate, unrealistically assuming the system would consist of only ribosomes translating at maximum rate. Notably, however, this upper limit is not much faster than the fastest growth observed, highlighting the importance of protein synthesis in defining the timescale of growth. For example, the maximal translation rate for *E. coli* is 10 ≈ hr^−1^ and thus only ≈4 times higher than the growth rates in rich LB media (*λ* ≈ 2.5 hr^−1^). Including the synthesis of rRNA, another major component of the cellular dry mass, lowers this theoretical limit only marginally, further supporting our sole consideration of protein synthesis in defining growth (Appendix 10). The difference between measured growth rates and the theoretical limits can be mostly attributed to the synthesis of metabolic proteins which generate the precursors required for protein synthesis, which we consider next.

### Synthesis of Precursors

Microbial cells are generally capable of synthesizing precursors from nutrients available in the environment, such as sugars or organic acids. This synthesis is undertaken by a diverse array of metabolic proteins ranging from those which transport nutrients across the cell membrane, to the enzymes involved in energy generation (such as those of fermentation or respiration), and the enzymes providing the building blocks for protein synthesis (such as those involved in the synthesis of amino acids). While these enzymes vary in their abundance and kinetics, we group them all into single set of metabolic proteins with a mass *M*_*Mb*_ which cooperate to synthesize the collective pool of precursors from nutrients required for protein synthesis. We make the approximation that these metabolic proteins generate precursors at an effective *metabolic rate ν*. In general, this rate depends on the concentration of nutrients *c*_*nt*_ in the environment. This relation is canonically described by a Monod (Michaelis-Menten) relation

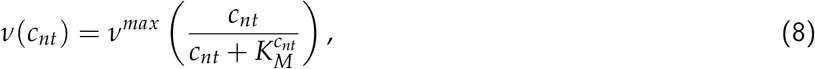

where *ν*_*max*_ is the maximum metabolic rate describing how fast the metabolic proteins can synthesize precursors, and 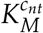 is the Monod constant describing the concentration below which nutrient utilization slows [90]. Novel precursors are thus supplied with a total rate of *ν*(*c*_*nt*_)*M*_*Mb*_ and consumed via protein synthesis at a rate *γ*(*c*_*pc*_)*M*_*Rb*_. Translation relies on precursors and, as introduced above, the translation rate *γ*(*c*_*pc*_)*M*_*Rb*_ thus depends on the concentration of precursors in the cell, *c*_*pc*_. As we do not explicitly model cell division, we here approximate this cellular concentration as the relative mass abundance of precursors to total protein biomass. This approximation is justified by the observation that cellular mass density and total protein content is approximately constant across a wide range of conditions [33, 91, 92]. The dynamics of precursor concentration follows from the balance of synthesis, consumption, and dilution as the total biomass grows:

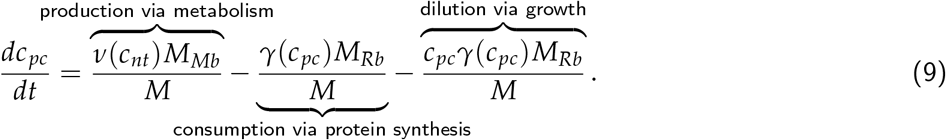

While the dilution term is often assumed to be negligible, this term is critical to describe growth and derive analytical expressions (detailed discussion in Appendix 3). Furthermore, we note that the precursor concentration is defined such that the consumption of one precursor yields the addition of one amino acid to the biomass *M*. As we measure proteins in units of amino acids there is thus no conversion factor needed when describing the consumption of precursors by protein synthesis.

### Simplification of saturating nutrients

The introduced dynamics simplifies when the nutrient concentration in the environment *c*_*nt*_ well exceeds the Monod constant 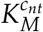 as *ν*(*c*_*nt*_) simplifies to *ν*_*max*_. Steady growth for which biomass increases exponentially readily emerges. This is the scenario we focus on in in the first half of this work. It should be noted however, that biologically such a scenario can only be realized temporarily as the nutrient supply required by the exponentially growing biomass can only be sustained by the environment for a limited amount of time. In general, the nutrient levels vary.

### Consumption of Nutrients in batch culture growth

The synthesis of novel precursors relies on the availability of nutrients which changes depending on the environment. In Supplementary Figure 2 we consider specifically a “batch culture” scenario in which nutrients are provided only at the beginning of growth and are never replenished. Therefore, growth of the culture continues until all of the nutrients have been consumed. The concentration of nutrients in the environment is thus given as

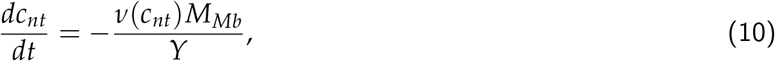

where *Y* is the yield coefficient which describes how many nutrient molecules are needed to produce one unit of precursors.

### Ribosomal Allocation of Protein Synthesis

As final step of the model definition, we must describe how cells direct their protein synthesis towards making ribosomes, metabolic proteins, or all other proteins that make up the cell [colored arrows in Fig. 1(A)]. We do so by introducing three *allocation parameters φ*_*Rb*_, *φ*_*Mb*_, and *φ*_*O*_ (such that *φ*_*Rb*_ + *φ*_*Mb*_ + *φ*_*O*_ = 1) which define how novel protein synthesis is partitioned among these categories:

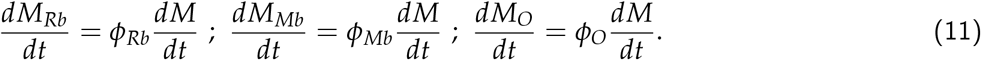

These equations are summarized in Fig.1(B) and Fig. S3 and define the accumulation of biomass, from nutrient uptake to protein synthesis.

### Approximating concentration via relative abundance

In addition to maintaining the *total* macromolecular densities, cells also maintain an approximately constant protein density [93]. This observation allows for a major simplification when formulating the allocation model, namely the approximation of concentrations as relative mass abundances. The rate *γ* at which ribosomes can synthesize protein is dependent on the abundance of precursors, *c*_*pc*_, in the cell. To compute the concentration and/or density in typical units (e.g. *µ*M, or mass / volume), we would require some measure of the total cellular volume, *V*_*cell*_, such that the concentration follows

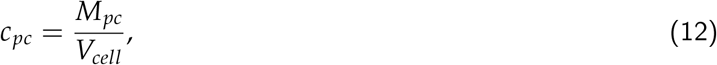

with *M*_*pc*_ denoting the total mass of the precursor pool. By making the experimentally-supported assertion that the protein density *ρ* is constant, we can say that

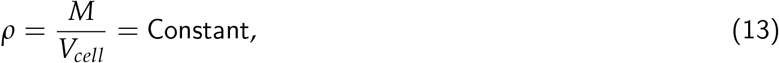

where *M* is the total protein biomass. Thus, the total cellular volume *V*_*cell*_ can be computed as

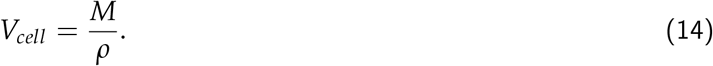

Plugging this result into Eq. 12, we arrive at the approximation

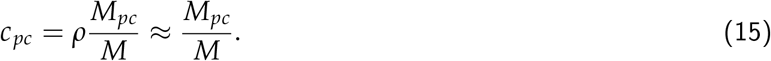

In this work, we neglect *ρ* as a multiplicative constant, and treat *c*_*pc*_ as being dimensionless. We direct the reader to Refs. [47] and [94] for a further discussion of the conversion between concentration and relative abundance.

### Derivation of Analytical Expressions

In the first section of this work, we present several analytical relations pertinent to steady-state growth. These relations follow from the simple allocation model and describe (i) how the growth rate depends on model parameters (Fig. 1C) and (ii) how ribosome content depends on other model parameters for the three different regulation scenarios we discuss (Fig. 1D and E). Here, we introduce a step-by-step derivation of these expressions.

### Deriving the Steady-State Growth Rate

We begin with deriving an expression for the steady-state growth rate *λ* which is similar to previous approaches taken by Giordano *et al* [55] *and Dourado et al* [61]. *As discussed in Supplementary Figure 2, steady-state conditions are satisfied when two conditions are met. First, the dynamics of the precursor concentration is constant (i*.*e*. 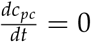) and the composition of the proteome matches the allocation parameters (i.e. 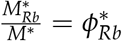 and 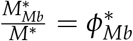). Furthermore, we assume that in steady-state growth, the concentration of nutrients in the environment is saturating 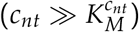, meaning that *ν*(*c*_*nt*_) *≈ ν*_*max*_. With these conditions satisfied, we can rewrite Eq. 9 as

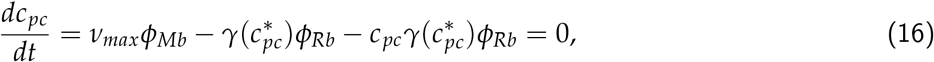

where 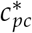 is the steady-state precursor concentration.

Noting that in steady-state conditions, the total biomass increases exponentially at a rate 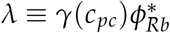, Eq. 16 can be simplified to

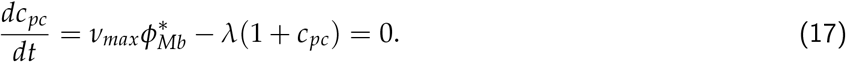

We can therefore solve for the steady-state precursor concentration 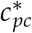 to yield

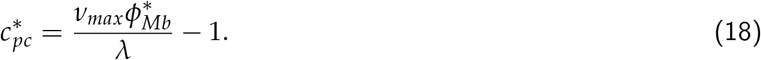

Assuming a Michaelis-Menten form for the translation rate 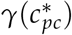, we can now define it as a function of the growth rate *λ* as

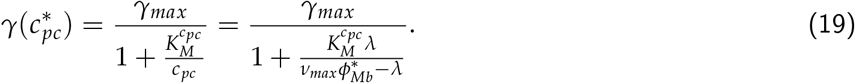

Knowing that the growth rate 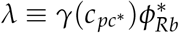, and 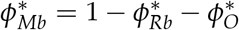 we say that

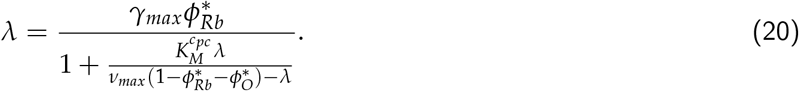

This can be algebraically manipulated to yield a quadratic equation of the form

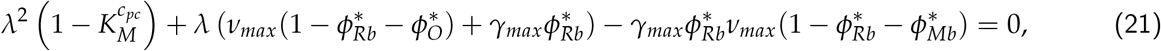

which has one positive root of

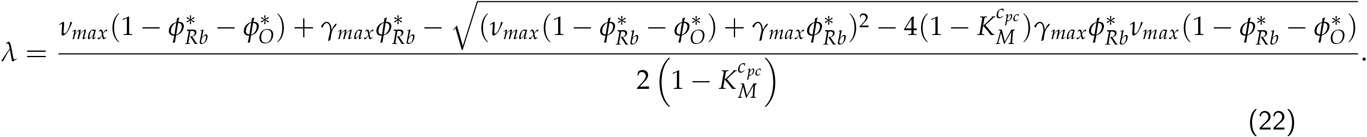

For notational simplicity, we can define the maximum metabolic output and the maximum translational output as N = *ν*_*max*_(1 *− φ*_*Rb*_ *− φ*_*O*_) and Γ = *γ*_*max*_*φ*_*Rb*_, respectively, and substitute them into Eq. 22 to generate

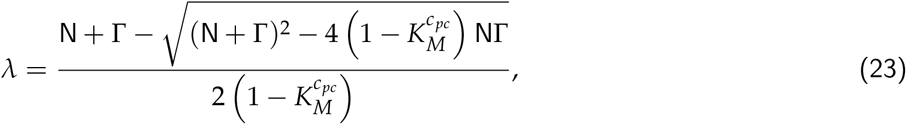

## Defining *φ*_*Rb*_ For Scenarios II and III

In Fig. 1(D), we provide a description of three plausible regulatory scenarios microbes may employ to regulate their ribosomal content. Scenario I assumes just a constant, arbitrary allocation parameter *φ*_*Rb*_ *∈* [0, 1 − *φ*_*O*_]. Here, we provide a short derivation for the more complicated relations describing ribosomal content under scenarios II and III.

### Scenario II: Constant Translation Rate

The second regulatory scenario assumes that the ribosomal content is adjusted to maintain a specific standing concentration of precursors, which we denote as 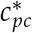. Noting that the growth rate 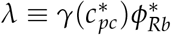, we can restate Eq. 18 in the form

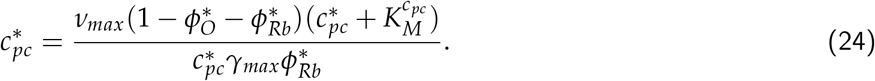

Some algebraic rearrangement allows us to solve for 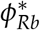, yielding

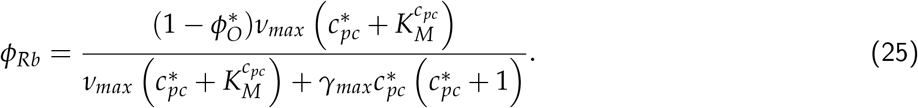

This expression is equivalent to that shown for scenario II in Fig. 3 of the main text. In evaluating this scenario, we considered the regime in which precursors were in abundance, meaning 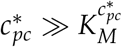. Under this regime, Eq. 25 simplifies further to

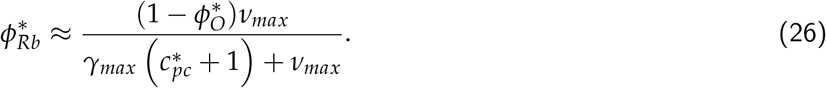

This represents a strategy where the cell adjusts 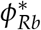 to maintain a translation rate very close to *γ*_*max*_.

### Scenario III: Optimal Allocation

In this work, we define the optimal allocation of ribosomes 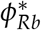 to be that which maximizes the growth rate in a given environment and at a given metabolic state. To determine the optimal 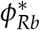, we can differentiate Eq. 22 with respect to 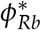 to yield the cumbersome expression

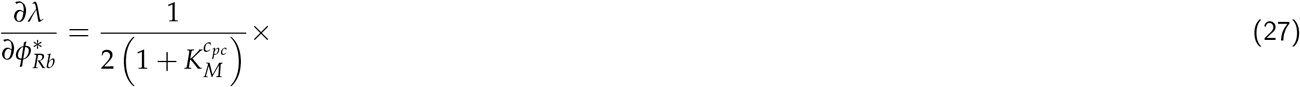

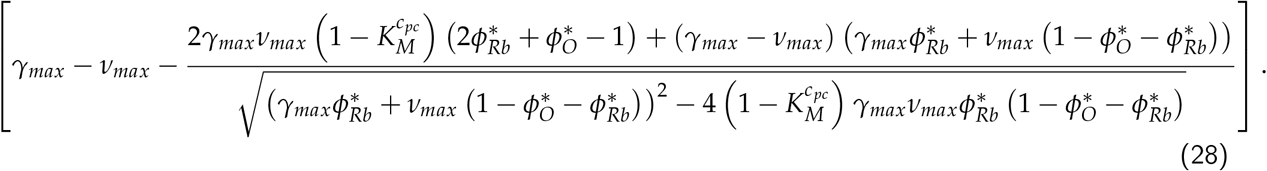

Setting this expression equal to zero and solving for *φ*_*Rb*_ results in

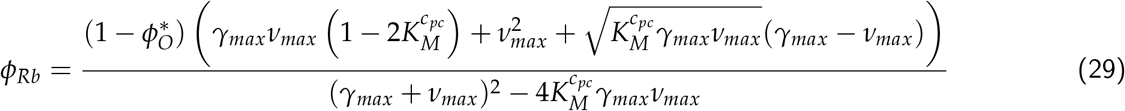

which is the optimal allocation towards ribosomes as presented in Fig. 1D of the main text.

### Implementing Flux-Parity Regulation via ppGpp

Here we expand upon and derive the equations defining the flux-parity allocation model shown schematically in Fig. 2(B) and explore its dependence on parameter values.

### Formulation of model

To include ppGpp signaling into the ribosomal allocation model, we must perform two tasks. First, we must explicitly model the dynamics of both charged- and uncharged-tRNAs. Secondly, we must tie the relative abundances of these tRNAs to the allocation parameters such that when charged-tRNAs are limiting and uncharged-tRNAs in abundance, the system reacts by adjusting the allocation parameters towards ribosomal proteins and away from metabolic proteins (*φ*_*Rb*_ and *φ*_*Mb*_).

We consider there to be two pools of tRNAs – those charged with an amino acid (denoted as *tRNA*^*c*^) and those that are uncharged (*tRNA*^*u*^). Rather than keeping track of the copy numbers of these tRNAs, we instead model their concentration as relative mass abundances (relative to the total protein biomass *M*), treating each tRNA to have an effective mass of one amino acid as each tRNA can in principle be charged. Much as for consideration of precursors in the simpler model we can model the concentration dynamics of these pools of tRNAs by considering three processes – the generation of the tRNAs, the consumption of the tRNAs, and the effect of dilution as the biomass grows.

We begin first with modeling the dynamics of the charged-tRNA pool, *tRNA*^*c*^. Here, we consider that charged-tRNAs are synthesized from one free amino acid and one uncharged-tRNA and further assume that the pool of free amino acids is abundant enough such that the tRNA pool is the rate limiting component. Making this assumption allows us to state that the conversion of one uncharged-tRNA to one charged-tRNA via the metabolic machinery proceeds at a rate *ν*(*tRNA*^*u*^), itself dependent on the *uncharged-tRNA*^*u*^ concentration. Likewise, we consider that the conversion of one charged-tRNA to an uncharged-tRNA is only possible via protein synthesis, which proceeds at a rate *γ*(*tRNA*^*c*^) that is dependent on the *charged-tRNA* concentration. Finally, as is discussed in Sec. **??** of this supplement, we must also consider how the mere fact of growing biomass effectively dilutes the charged-tRNA concentration. Together, these processes can be combined to enumerate the dynamics of the charged-tRNA pool as

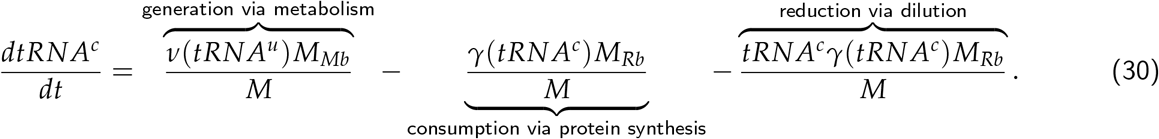

The dynamics for the pool of uncharged-tRNAs can be constructed in a similar manner, with the caveat that the generation of new uncharged-tRNAs occurs from both protein synthesis (converting one charged-tRNA into one uncharged-*tRNA*^*u*^) and from transcription of the individual tRNA genes. We consider the latter to occur at a rate *κ*, which has dimensions of concentration per unit time. Using the same logic of mapping the productive and consumptive processes, we can enumerate the dynamics of the uncharged-tRNA pool as

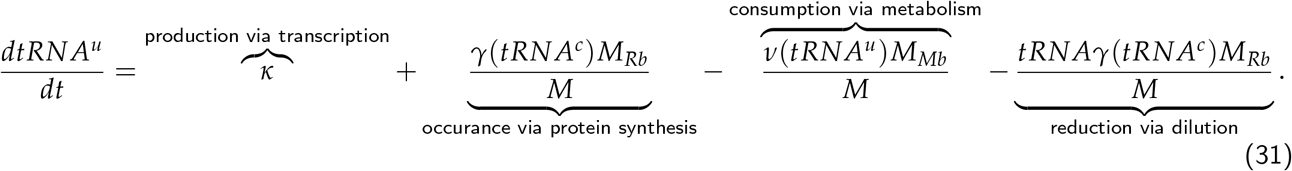

These expressions comprehensively define the dynamics of the tRNA pool, from generation via transcription to their recycling between charged and uncharged states through metabolic and translational fluxes, respectively. As in the main text, we posit that the dynamics of the ribosomal *M*_*Rb*_, metabolic *M*_*Mb*_, and “other” *M*_*O*_ protein masses follow via the allocation parameters *φ*_*Rb*_, *φ*_*Mb*_, and *φ*_*O*_ respectively. However, in this treatment of the model, we consider these parameters, with the exception of *φ*_*O*_, to be dynamic and depending on the intracellular concentration of ppGpp. Mathematically, we state this as

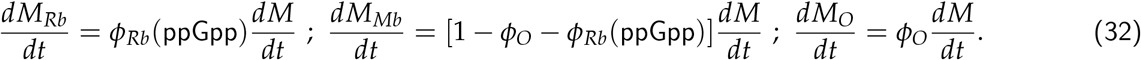

We are now tasked with (i) enumerating the dynamics of ppGpp and (ii) assigning a specific functional form to *φ*_*Rb*_(ppGpp). The biochemistry of ppGpp synthesis, degradation, and binding to the transcription machinery has been studied in *E. coli* among other prokaryotes, revealing the enzyme(s) important for this process. In *E. coli* RelA and SpoT. Many molecular details revealing how those enzymes control ppGpp levels in response to the abundance of tRNA levels are known but important details also remain puzzling [79, 80]. Thus, while previous works have consider the dynamics of these specific proteins in more detail [55, 65], we here take a more coarse-grained view. Specifically, we first make the ansatz that the dynamics of ppGpp synthesis and degradation are sufficiently fast compared to the timescale of protein synthesis such that it can be treated as being in steady-state instantaneously. Secondly, we take the concentration of ppGpp to be inversely proportional to the relative abundance of charged-to uncharged-tRNAs,

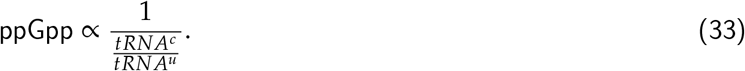

This is a well-motivated starting point as in *E. coli*, ppGpp is primarily synthesized via RelA when an uncharged-tRNA enters the A-site of a translating ribosome, forming a stalled complex. As binding of a charged-tRNA or an uncharged-tRNA is a competitive process, the probability of one or the other being bound is dependent on their relative concentrations, rather than the absolute concentrations of either species. However other processes which affect ppGpp levels, including the synthesis and degradation by SpoT in relation to ribosome activity, are less well understood [95]. Accordingly, we consider our approach to describe ppGpp as inversely proportional to the relative abundance of charged-to uncharged-tRNAs as a motivated ansatz rather than a fully established biochemical relation. And we furthermore show below that this ansatz works much better for describing the experimental observations as a few different ones we probed.

Given the relation between ppGpp and tRNA charging ratio, Eq. (33), we can now define the allocation towards ribosomes to be a function of the tRNA charging ratio, 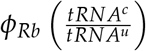. To assign a specific functional form to this relation, we assume that the expression of ribosomal genes is in first order described by a simple binding kinetics of ppGpp to the transcriptional machinery and the allocation towards ribosomes follows a form similar to that of a Michaelis-Menten relation,

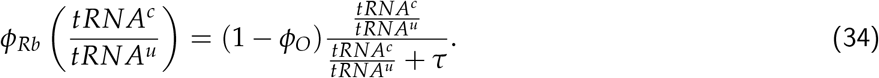

Here, the parameter *τ* represents the value of the charged-to uncharged-tRNA ratio where *φ*_*Rb*_ is at its half-maximal value. The maximal value itself depends on the magnitude of *φ*_*O*_, the allocation towards other proteins, which we are considering to be independent of ppGpp; 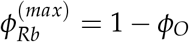.

The transcription of tRNA genes towards novel tRNA synthesis has also been shown to be regulated with ppGpp, appearing to closely match the regulatory behavior of ribosomal proteins [96]. We therefore model that the tRNA synthesis rate *κ* [introduced in Eq. (31)] is similarly modulated by the charged-to uncharged-tRNA ratio,

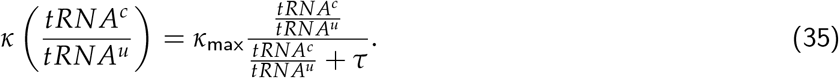

Here, *κ*_*max*_ is the rate of tRNA transcription when all tRNA genes are fully saturated with RNA polymerase in rich growth conditions where gene dosage is high. Finally, we must establish functional forms for the tRNA dependencies on the metabolic and translation rate. Simple biochemical assumptions permit a formulation of a Michaelis-Menten function for each rate. Noting that the translation rate *γ* is defined as 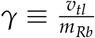, where *v*_*tl*_ is the translation speed and *m*_*Rb*_ is the proteinaceous mass of a single ribosome, we take *γ*(*tRNA*) to be of the form

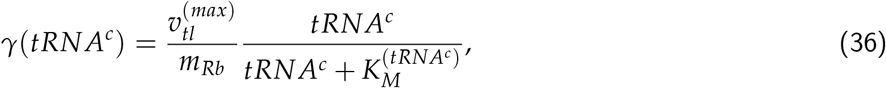

where 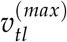 is the maximum translation speed and 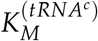 is the Michaelis-Menten constant. A similar argument can be made for the dependence of the metabolic rate *ν* on the uncharged-tRNA concentration,

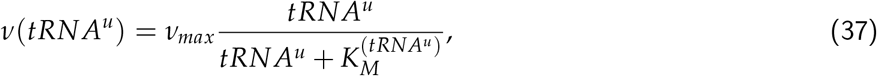

with 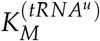 being another Michaelis-Menten constant. Together, Equations 30 through 37 mathematically describe a model for ppGpp-dependent regulation of translational and metabolic fluxes.

In principle, an analytical solution for this system of ODEs can be found, though it precludes evaluation by hand and is computationally intensive. While we do not solve this system of ODEs analytically here, we can numerically integrate them to sufficiently approximate the steady-state behavior. Depending on the choice of parameter values, such an approach can yield an allocation scenario nearly indistinguishable from that of the optimal allocation scenario (scenario III) of the simple model [Fig. 1(G and H)].

### Optimal allocation emerges from flux-parity regulation

While the previous section lays out the mathematics of the flux-parity model, we now discuss how this regulation scheme can lead to an optimal allocation. Towards this goal we first discuss in more detail what we mean when we say ‘flux-parity’. As described in the main text, we define flux-parity as a balance *and* mutual maximization of (i) the flux of uncharged-tRNAs through metabolism (termed the *metabolic flux J*_*Mb*_) and (ii) the flux of charged-tRNAs through protein synthesis (termed the *translational flux J*_*Tl*_). To demonstrate this point, assume that we can decouple the dependence of the allocation parameter *φ*_*Rb*_ from the ratio of charged-to uncharged-tRNAs. Mathematically speaking, we can define the metabolic flux as the collective action of metabolic proteins,

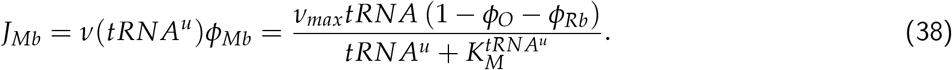

Similarly, we can state that the translational flux is the collective action of ribosomal proteins,

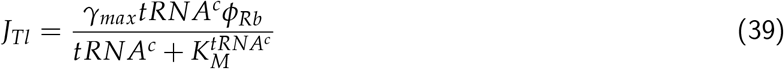

So long as these fluxes are equivalent, a steady-state is satisfied. However, this steady-state is not necessarily the optimal value. This is illustrated in Fig. S5. For example, if we consider that *φ*_*Rb*_ is too large for the given condition [Fig. S5(left)], a specific steady-state is realized (black point). If *φ*_*Rb*_ is further increased, the value of both the metabolic and translational fluxes (dashed lines) must decrease to reach a new steady-state and growth-rate thus declines. However, if *φ*_*Rb*_ is decreased, the value of both fluxes increase and growth-rate thus also increases as well. At optimum allocation [where growth is locally maximized, Fig. 5(middle)], any perturbation to *φ*_*Rb*_ will necessarily results in a decrease in the fluxes, indicating that at the optimal allocation the fluxes are *mutually maximized*.

As the concentrations of both tRNA species (Eqs. 30 and 31) are dependent on the allocation towards ribosomes *φ*_*Rb*_ in inverse ways, the ratio of their concentrations acts as an effective sensor of the magnitude of either flux. A large charged-to uncharged-tRNA ratio indicates that there is an abundance of charged-tRNAs, suggesting that the translational flux is too low. Conversely, a small charged-to uncharged-tRNA ratio indicates a translational flux that is too large, diminishing the metabolic flux. By tying the allocation towards ribosomes *φ*_*Rb*_ to this ratio, an allocation can emerge that optimizes the fluxes and thus growth.

### Assessing different assumptions of *φ*_*Rb*_ dependence on ppGpp

In Eq. 33, we made the assumption that the concentration of ppGpp was inversely proportional to the charging balance of the tRNA pools. We put this forward as an ansatz with the motivation that the degree of tRNA charging should be related to the amount of ppGpp synthesized. However, there are other ansatzes that could be made relating the amount of ppGpp to the individual concentrations of the tRNAs, or other ratiometric definitions.

To test how sensitive our findings are to the particular ansatz used, we considered other ways in which the ppGpp concentration could be related to the tRNA pools. There is strong biochemical evidence that a primary route of ppGpp synthsesis is via the enzyme RelA, which becomes active when associated to a “stalled” ribosome – one that is bound to an uncharged tRNA – though some details remain enigmatic. In manner similar to other works [42, 55, 65], we can assert that the amount of ppGpp is proportional to the abundance of stalled ribosomes. Mathematically, we can define the ppGpp concentration as being proportional to the probability of a ribosome binding an uncharged tRNA. Assuming that the tRNA concentration (of both charged and uncharged forms) is sufficiently high that all ribosomes are complexed with a tRNA, this equates to

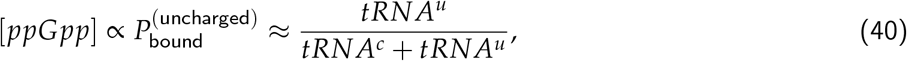

where *tRNA*^*c*^ and *tRNA*^*u*^ represent the absolute concentrations of charged and uncharged species, respectively. If the ppGpp concentration is inversely proportional to the allocation towards ribosomes, we can similarly make the argument that the ribosomal allocation *φ*_*Rb*_ will be proportional to the probability of a ribosome being bound to a *charged-tRNA*,

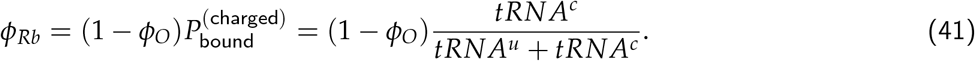

This equation mechanistically operates in a similar way as Eq. 34 – the allocation towards ribosomes depends on the relative amounts of charged- and uncharged-tRNAs. In the extreme limit where the total concentration of tRNA is fixed (for which there is conflicting evidence [55, 65, 77, 78, 93]), Eq. 41 and Eq. 34 are mathematically equivalent. However, the predicted scaling dependence of ppGpp takes a different form.

In the main text, we noted that the concentration of ppGpp relative to a reference growth rate 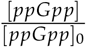 is equivalent to the inverse ratio of the charging balances. Under the ansatz that the ppGpp concentration is depending on the uncharged-tRNA binding probability, this relation takes the form

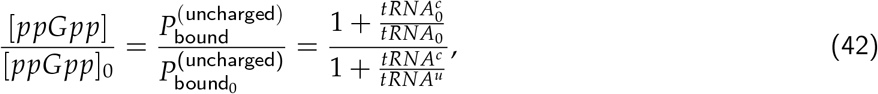

where the subscript 0 denotes the reference state value. This distinction, coupled with experimental measurements of the relative ppGpp concentrations, allows us to test the validity of the two assumed forms for *φ*_*Rb*_.

i Fig. S6 shows the predictive capacity of these two ansatzes with the simple binding (Eq. 41) and ratiometric (Eq. 33) predictions shown in solid-blue and dashed-red lines, respectively. While both of these assumptions are capable of predicting the scaling of the ribosome content and translation speed with quantitative equivalence, there is a distinct difference in the predicted behavior of the relative ppGpp concentrations. The simple binding ansatz predicts a significantly shallower dependence on the growth rate than is observed in the data and in the ratiometric prediction. Thus, it appears that relating [ppGpp] to the ratio of uncharged-to charged-tRNA concentrations accurately captures the behavior of *E. coli*, though there remain gaps in our understanding of this relationship at a biochemical level.

### Incorporating effects of ribosome-targeting antibiotics

To extend the flux-parity allocation model and incorporate the effects of antibiotic treatment, we must consider the mechanism of action of the antibiotic, specifically chloramphenicol. Chloramphenicol is a bacteriostatic antibiotic with tightly, but reversibly, binds to the ribosome. Once bound, the ribosome is unable to resume translation until chloramphenicol dissociates. Thus, we can model the effect of this drug by enumerating the probability that chloramphenicol is bound to a ribosome *P*_bound_ at a given concentration *c*_*cm*_. Mathematically, this can be stated as

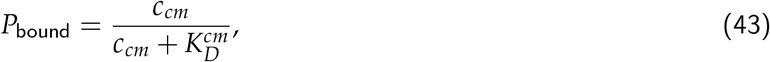

where 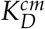 is an effective dissociation constant of chloramphenicol to a unit of ribosomal mass accounting for kinetics transport and ribosome binding. We can then say that the probability of a ribosome being active is equal to the probability of a ribosome being *unbound*,

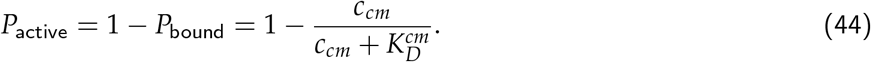

As only active ribosomes will contribute to the accumulation of biomass, we must rewrite the dynamics as

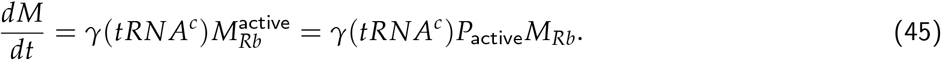

To make the predictions shown in Fig. 3(E and F) of the main text, we assumed that the chloramphenicol concentration in the growth medium is equal to the intracellular concentration and take 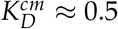.

### Incorporating effects of excess protein stress

We consider that the excess protein synthesis can be modeled as the introduction of a new protein class, which we consider to have a total mass of *M*_*X*_. Following the allocation parameters of the flux-parity model as defined in Eq. 32, we can introduce a new allocation parameter *φ*_*X*_ such that

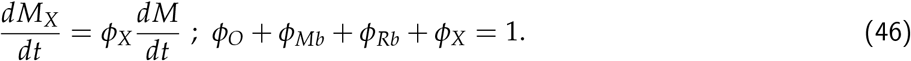

In the right-hand panel of Fig. 5B in the main text, we show that a collection of data can be collapsed onto a single line that relates the relative change in growth rate as a function of the excess protein that is synthesized. While we cannot fully solve the flux-parity model analytically, we can derive an analytical expression of this relation. Specifically, we note that the steady-state growth rate in the absence of excess expression *λ* follows the simple relation

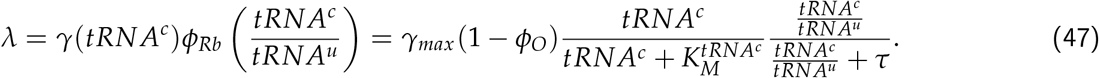

This can be easily extended to compute the growth rate under excess protein synthesis *λ*_*X*_ as

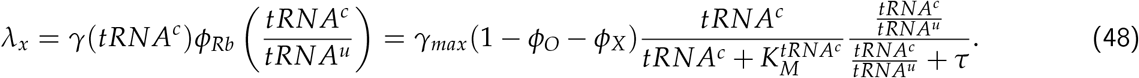

We can take the ratio of these growth rates to yield an expression for the collapse function

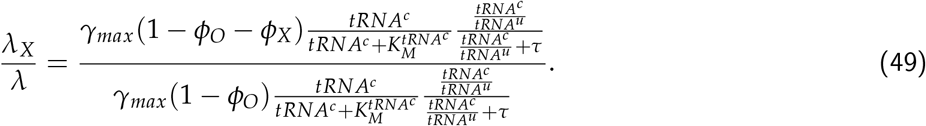

If we assume that the excess protein synthesis affects *only φ*_*X*_, leaving all other parameters untouched, Eq. 49 reduces to the concise form

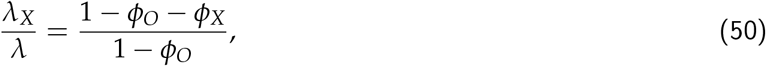

which is the linear relation plotted in the right-hand side of Fig. 5B of the main text.

Aside from the collapse, we also show how the flux-parity model quantitatively predicts the growth rate as a function of excess protein for three different media (Fig. 5C, center panel). In this case, we require some knowledge of what the metabolic rate *ν*_*max*_ is for those specific conditions. As the metabolic rate is an efficient rate incorporating the action of different metabolic reactions and serving as a proxy of the nutrient quality, it is not possible to make an *a priori* estimate of its value. To nevertheless estimate *ν*_*max*_ for each condition, we determined its value by using the simple allocation model as encoded in Sec., assuming the growth rate *λ* and the ribosomal content describes the allocation towards ribosomes *φ*_*Rb*_. Under the simple allocation model, we note that an expression for the metabolic rate can be solved from the steady-state precursor concentration. 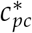 (Eq. 18) to yield

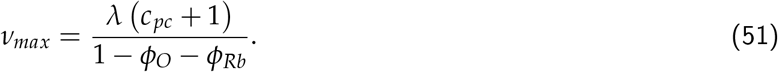

The steady-state precursor concentration 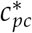 can be solved from the definition of the steady-state growth rate and has the form

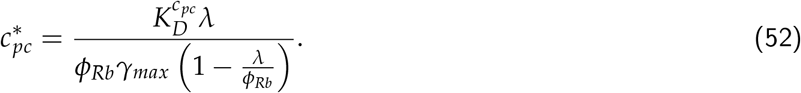

Combining Eqs. 51 and 52 yields an expression for the maximal metabolic rate *ν*_*max*_,

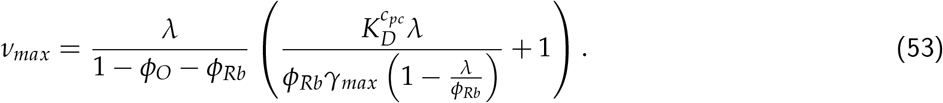

Thus, given knowledge of the steady-state growth rate *λ* and the allocation towards ribosomes *φRb* (which are both measured quantities), the value of *ν*_*max*_ can be derived.

### Incorporating effects of nutrient upshifts

To model the dynamics of growth in fluctuating conditions, we asserted that a nutritional upshift is equivalent to an instantaneous change in the metabolic rate such that 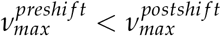. However, this is not completely sufficient to capture the phenomenology that is observed. It is becoming exceedingly clear that bacterial cells are non-optimal in *what* genes they express, with many proteins that are synthesized are ultimately useless in the specific condition [64]. This can have very important effects on the growth rate as any amount of conditionally useless protein that’s synthesized consumes resources that could otherwise be partitioned to the proteins that need to be synthesized. To incorporate this effect, we introduce another protein class with an allocation parameter *φ*_ø_. As the degree of conditionally useless expression is significantly more pronounced in slow rather than fast conditions [27, 33, 64], we further asserted that the magnitude of this sector also changed in response to the nutritional upshift such that 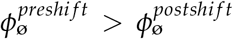. The precise value of this sector is less important than the difference in the pre- and post-shift condition and can be considered as an additional rescaling factor as described in Appendix 5. Thus, for all nutritional shifts in this work, we considered that 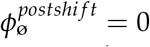 and the value of 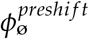 to be linearly proportional to the difference in the growth rates between the pre- and post-shift conditions.

### Incorporating effects of nutrient depletion

Up to this point, we have explored the flux-parity model under the assumption that the nutrients in the environment were saturating, such that *ν*(*c*_*nt*_) *≈ ν*_*max*_. However, a dependence on the environmental nutrient concentration *c*_*nt*_ can be easily included in the definition of the metabolic rate *ν* as

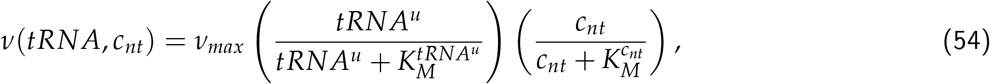

where 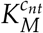 is the Michaelis-Menten constant. We can then model the dynamics of the nutrient concentration *c*_*nt*_ in a batch-culture system as

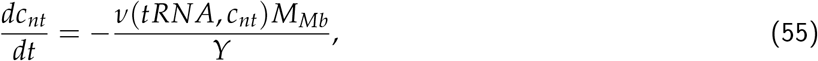

where *Y* is the yield coefficient.

### Data Sets

This work leverages a large collection of data, primarily from *E. coli*, to evaluate the accuracy of our model in describing biological phenomena. These data come from a range of studies spanning around 50 years of measurements from different groups and different geographical locations. Collecting and curating this large data set required the manual transcribing of data from papers as well as various standardization steps to ensure that measurements were truly comparable between studies, as is outlined in Table S2.

## Data and Code Availability

This work is accompanied by a website (cremerlab.github.io/flux parity) which houses a suite of interactive figures to help the reader gain an intuition for the way the theory operates. Data sets can be downloaded individually from this website. Alternatively, all data and Python code used in this work can be cloned as a GitHub repository (github.com/cremerlab/flux parity). This repository and its releases are also available via Zendodo with the DOI: 10.5281/zenodo.5893800.

## Acknowledgments

We thank Uri Alon, Markus Arnoldini, Rachel Banks, Suzy Beeler, Nathan Belliveau, Terence Hwa, Soichi Hirokawa, Christine Jacobs-Wagner, Sergey Kryazhimskiy, Armita Nourmohammad, Manuel Razo-Mejia, Tom Röschinger, Jan Skotheim, Cat Triandafillou, and members of the Tadashi Fukami, Dmitri Petrov, Alfred Spormann, and JC research groups for extensive discussions and the critical reading of the manuscript. GC acknowledges support by the NSF Postdoctoral Research Fellowships in Biology Program (Grant No. 2010807).

## Supplementary Figures

**Supplementary Figure 1:**
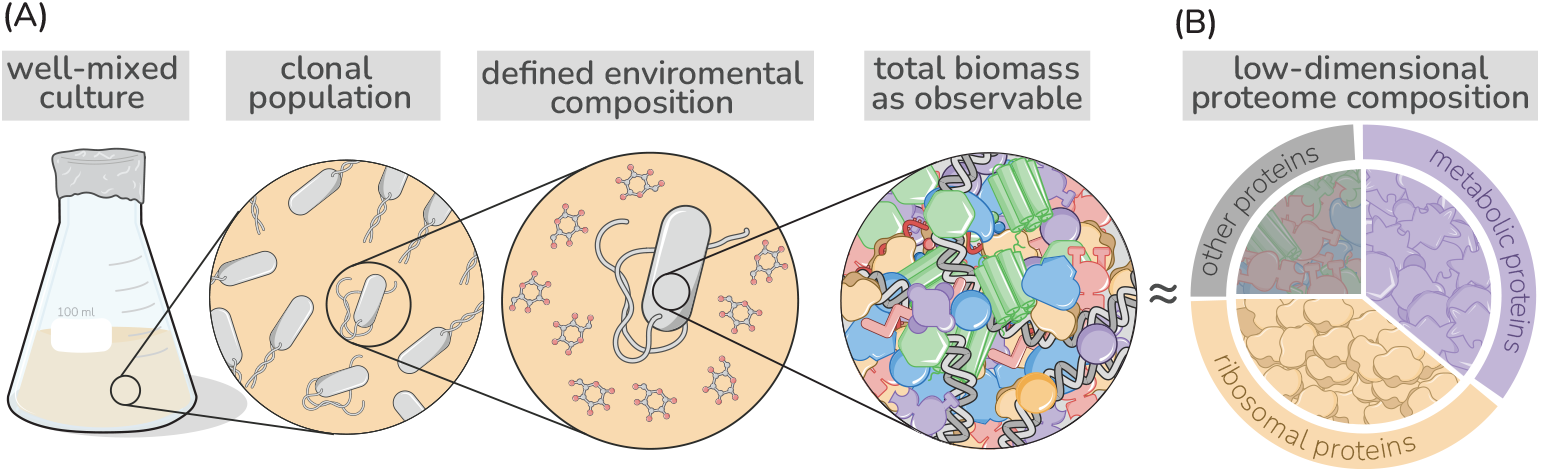
Coarse grained description of biomass and the proteome. Low-dimensional allocation models consider in their simplest form growth of a clonal population within a well-mixed environment. Biomass is described in a highly simplified manner focusing in the simplest case on protein synthesis alone [50, 52], with different protein species jointly considered by a few different protein classes. Here, metabolic proteins (purple), ribosomal proteins (gold), or “other” proteins (gray).

**Supplementary Figure 2:**
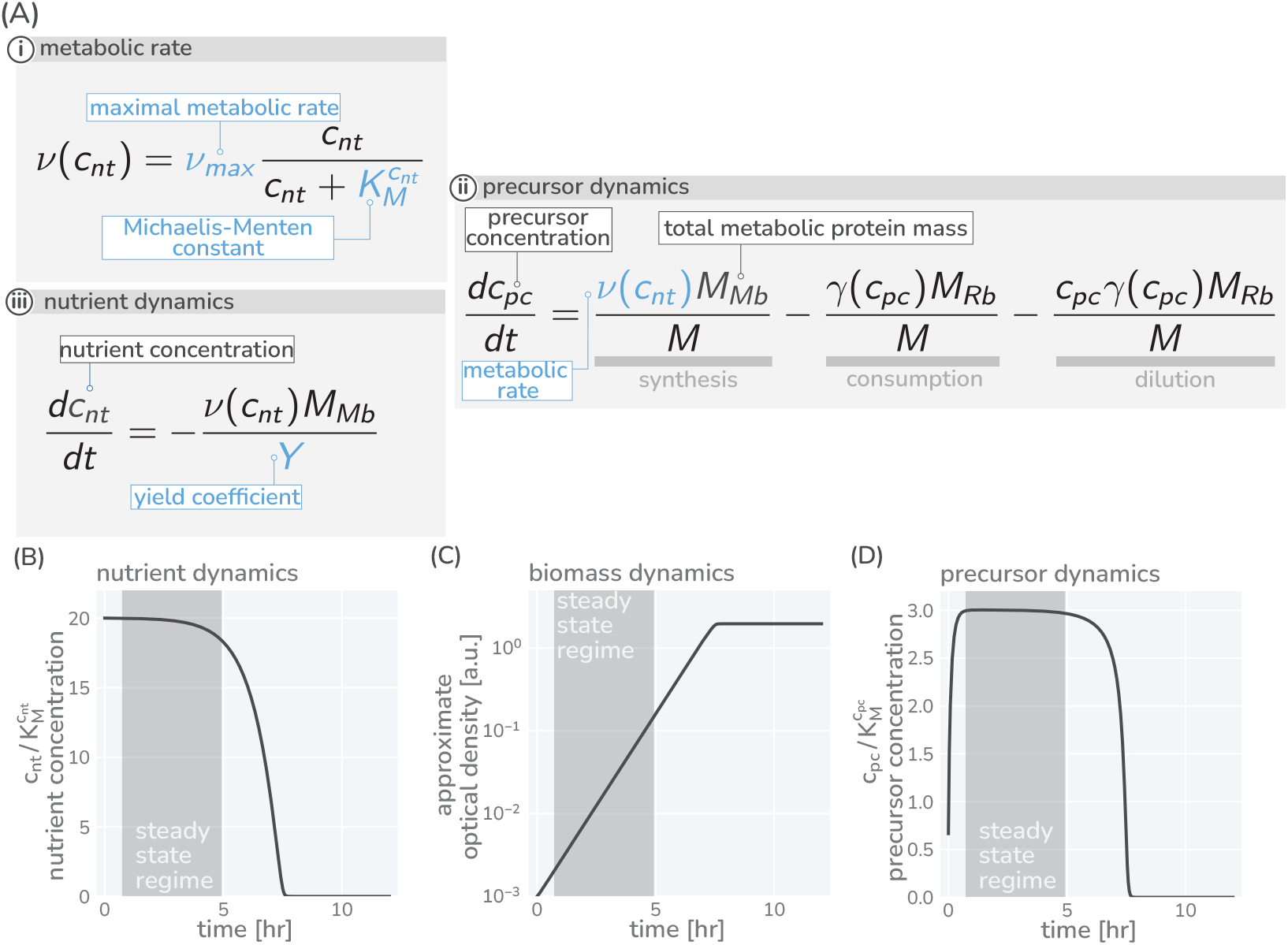
Precursor synthesis and growth when nutrients are not saturating. In general, the environmental conditions microbes encounter changes rapidly and nutrient availability commonly limits growth. Growing batch cultures, fox example run out of nutrients eventually and growth stops. The allocation modeling framework can account such a dynamics by including metabolic rates which depend on the nutrient concentrations in the environment. In the simplest case, one nutrient source is considered (concentration *n*) with the metabolic rate *ν*(*n*) depending on the concentration in a Michaelis-Menten manner with a maximal metabolic rate being reached only at high nutrient concentrations (A,i). The dynamics of precursors is given by a balance of synthesis, consumption, and dilution (A,ii), replacing the corresponding equation of the simple model in [Fig. 1(B,iv)]. The modeling of growth further requires the explicit modeling of nutrient concentrations. This dynamics depends on the specifics of the environment and, depending on the environment, can become very complex with multiple sources and sinks affecting the nutrient concentration. Here, we consider a typical batch culture scenario in which cells grow under well-mixed conditions. Nutrients are provided only initially and nutrient concentrations are falling because of consumption (A,iii). (B-D) Resulting temporal variation of nutrient concentrations (C), biomass accumulation (D), and precursor concentration (E) when integrating the model equations and using a parameter set descriptive of *E. coli* growing in a glucose-minimal medium with a growth rate ≈ 1 hr^−1^ and a starting glucose concentration of 10*mM*. As experimentally observed, initially abundant nutrients are consumed and biomass accumulates (exponential phase) until nutrients are exhausted and growth stops (saturation phase) (B and C). Importantly, precursor concentrations (D) quickly reach a constant plateau which lasts until nutrients become scare 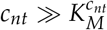 and *v*(*c*_*nt*_) ≈ *v*_*max*_. During this transient period (shaded regions) the synthesis of precursors matches the consumption by protein synthesis and dilution, meaning 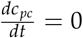. Given a constant precursor concentration 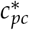, the translation rate 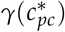 is also constant. As a consequence, the protein pool approaches a steady composition dictated by the allocation parameters 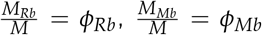 and 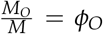. With precursor concentrations and protein composition remaining constant, the system is in a *steady-state* and biomass accumulates exponentially over time, 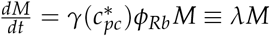. This is the steady state regime we focus on in the main text. Note that the steady state growth regime readily emerges when we consider dilution (see Appendix 3). Model parameters are provided in Table S1. Biomass units are converted to optical density assuming at *OD*_600*nm*_ = 1, there are 10^9^ cells per mL and 10^9^ amino acids per cell. An interactive version of these dynamics can be found on the paper website.

**Supplementary Figure 3:**
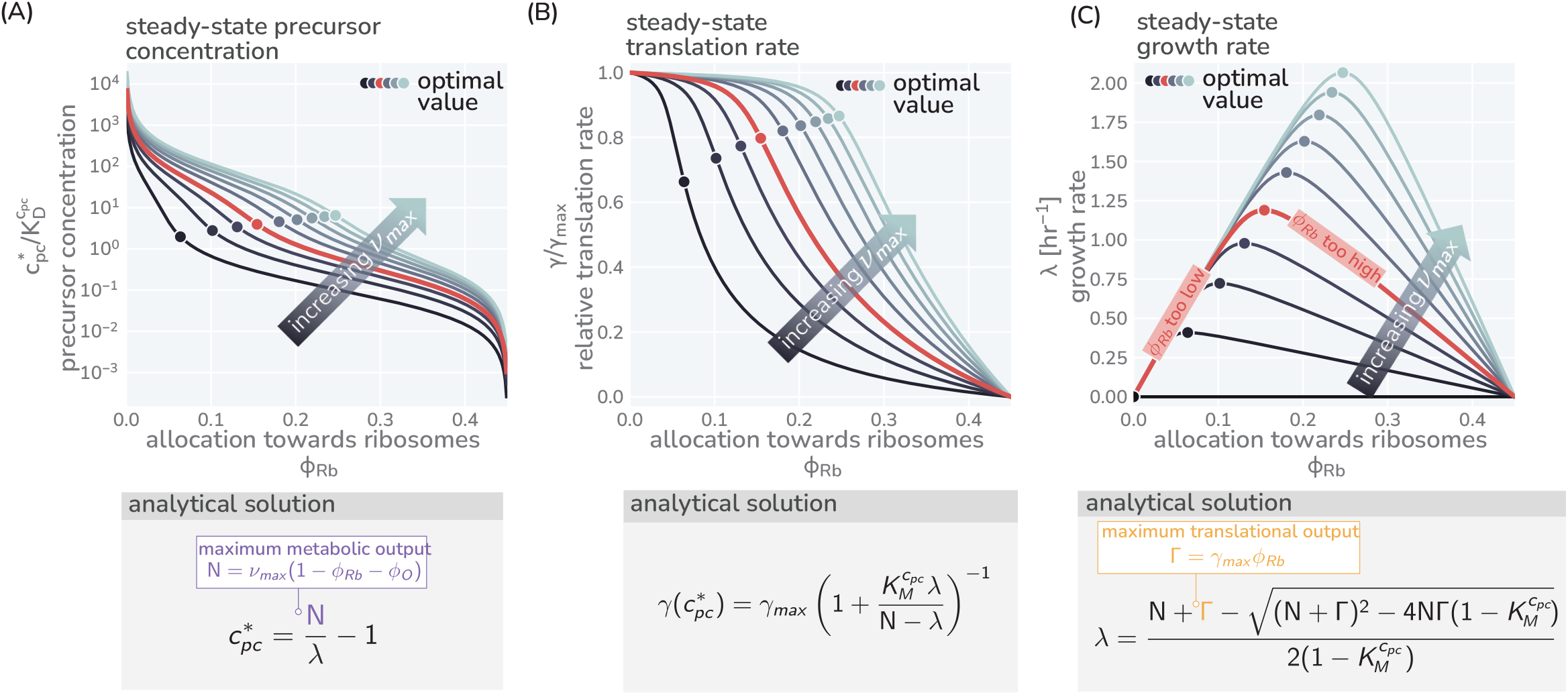
Modeling predictions of steady growth behavior. (A) Variation of the precursor concentration with varying allocation parameters (*φ*_*R*_) and maximal metabolic rate (*ν*_*max*_). (B C) Corresponding trends of translation and growth rate as also shown in Fig. 1 (C) and (D). Corresponding boxes show the analytical expression describing the steady-state precursor concentration, translation speed, and growth rate with details of the derivation provided in Appendix 7. Used model parameters provided in Table S1. Colors indicate different metabolic rates *ν*_*max*_ = 0.2 - 12.5 hr^−1^.

**Supplementary Figure 4:**
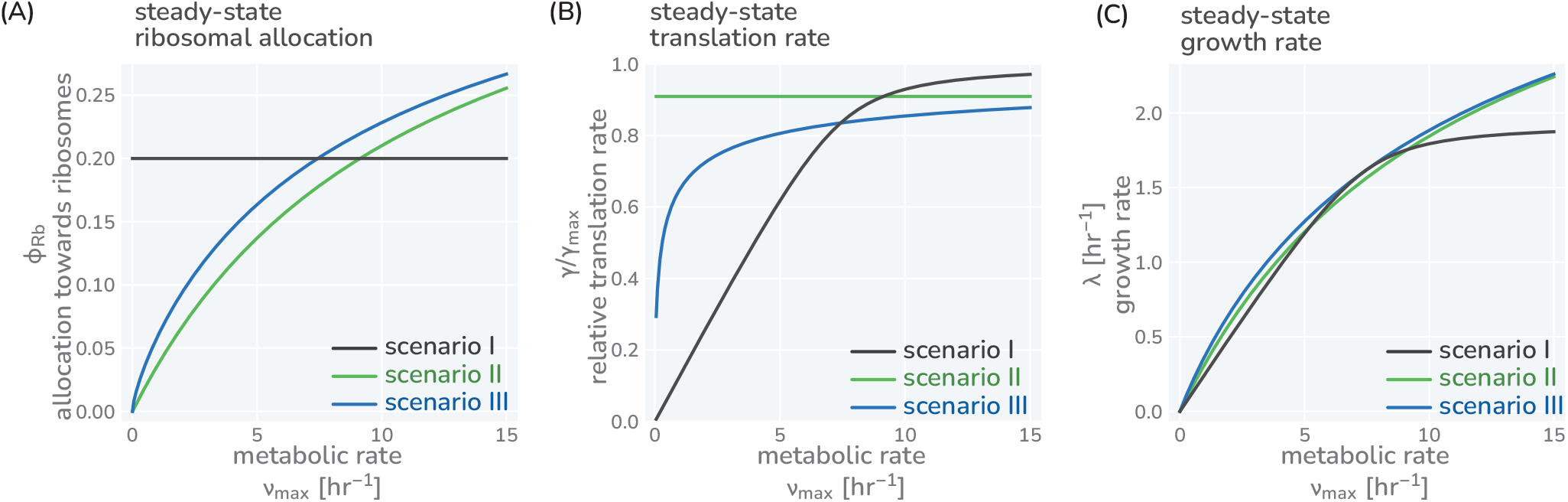
Three different allocation scenarios. The variation of precursor concentration (A), translation rate (B), and growth rate (C) with changing metabolic rate is shown for the three allocation scenarios introduced in the main text; fixed allocation (scenario I, black lines), prioritizing fast translation (scenario II, green lines), and growth-optimal allocation (scenario III, blue lines). Plotted are the analytical solutions provided in Fig. 1(E) and derived in the Methods. The resulting relations between growth rate and translation as well as growth rate and ribosome content are shown in Figure 1(G,H). We here discuss the consequence of these allocation scenarios in more detail. *Scenario I - fixed allocation*: In this scenario, allocation is fixed and does not vary with conditions. Locking in the ribosome allocation to *φ*_*Rb*_ = 0.25 (A, black line), for example, carries strong consequences for translation and growth rates (B and C, black lines). When conditions are poor (*ν*_*max*_ is small), the translation rate is significantly lower than the maximal rate as there are too many ribosomes competing for a small pool of precursors (B). The translation and growth rates increase with the metabolic rate *ν*_*max*_ until the influx of precursors is sufficiently high such that all ribosomes are translating close to their maximum and growth-rate is at its optimal value. Further increasing the metabolic rate does not increase the growth rate [plateau of black curve in (B)] as all ribosomes are already translating close to their maximum rate. *Scenario II - prioritizing fast translation*: In this scenario, allocation is adjusted such that translation rates are maintained at a high value. This is achieved by tuning the allocation between ribosomes and metabolic proteins such that a constant precursor concentration 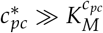 is maintained. For example, at higher metabolic rates, the metabolic proteins can sustain a higher influx of precursors allowing a larger allocation towards ribosomal proteins *φ*_*R*_ (green lines). *Scenario III - optimizing growth*: In this scenario, allocation is tuned to optimize growth rate across conditions, meaning that the fastest growth rate is achieved given a set metabolic rate and other model parameters. For example, the allocation towards ribosomes *φ*_*Rb*_ is adjusted with the metabolic rate such that the growth rate rests at the peak of the curves shown in Fig. 1(C) (blue lines). Accordingly, the growth rate continues to increase with higher metabolic rates always exceeding the growth rate of scenario I and II (C, green line)]. Model parameters follow the reference set for *E. coli* (Table S1). Black lines correspond to a constant allocation 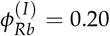 and green lines correspond to a constant precursor concentration 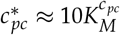, yielding a constant translation rate of 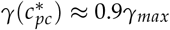. An interactive version of these figure panels is available on the paper website.

**Supplementary Figure 5:**
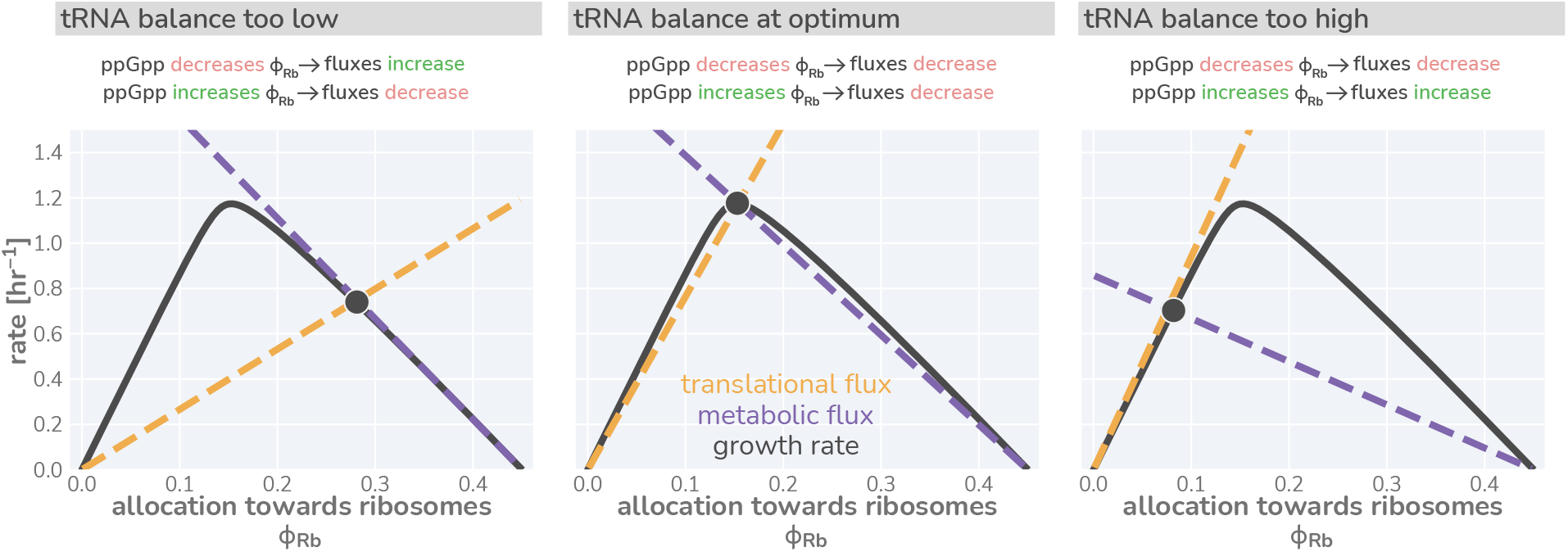
Flux-parity directs allocation parameters towards an optimum. Black lines represent the steady-state growth rate as a function of the allocation towards ribosomes *φ*_*Rb*_. Dashed gold and purple lines correspond to the translational and metabolic fluxes, with their intersection indicating the steady-state. The different panels consider from left to right three scenarios with a too low, optimal, and too high allocation towards ribosomes. An interactive version of this figure is available on the paper website

**Supplementary Figure 6:**
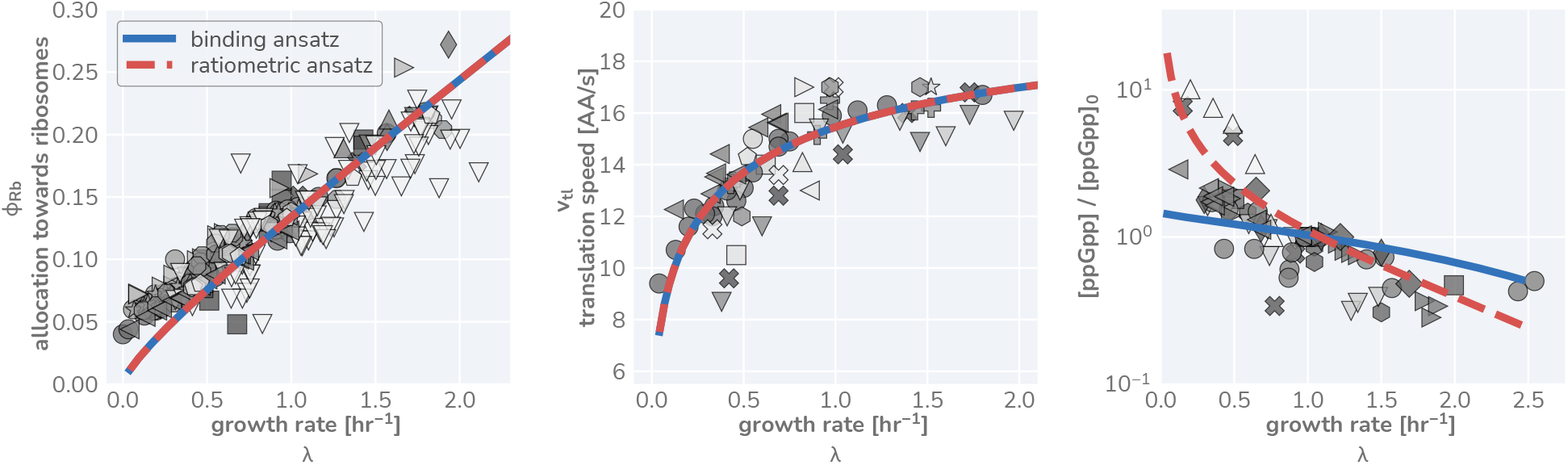
Comparison of predictive capacity of flux-parity allocation between ppGpp scaling ansatzes. Measurements are shown for ribosomal content, translation rate, and relative ppGpp concentration from left to right, respectively. Markers are the same as those in Fig. 3 of the main text. Solid blue line shows predicted steady-state behavior assuming a simple ansatz of ribosome-tRNA binding probabilities (Eq. 41). Dashed red line denotes predicted behavior using the ansatz that ppGpp concentration is dependent on the charging balance.

