## Appendix for "An Optimal Regulation of Fluxes Dictates Microbial Growth In and Out of Steady-State"

The past decade has seen a flurry of experimental studies further establishing the importance of protein synthesis in defining growth. Approaches include modern "-omics" techniques with molecular-level resolution [25–38], measurements of many core physiological processes and their coordination [38–46], and the perturbation of major cellular processes like translation [39, 47–49]. Together, these studies advanced a more thorough description of how cells allocate their ribosomes to the synthesis of different proteins depending on their metabolic state and the environmental conditions they encounter, called *ribosomal allocation*. Tied to the experimental studies, different theoretical *ribosomal allocation models* have further been formulated to dissect how ribosomal allocation influences growth [31, 32, 49–56, 56–64]. For example, high-dimensional models have been formulated which simulate hundreds to thousands of biological reactions [51,59] providing a detailed view of the emergence of distinct internal physiological states and the underlying processes which sustain them. Alternatively, other theoretical considerations follow coarse-grained approaches of moderate dimensionality which group different classes metabolic reactions together and mathematizicing their dynamics [60,62]. Distinct to those is further an array of extremely low-dimensional models, pioneered by Molenaar *et al.* 2009 [50], which have been developed to describe growth phenomena in varied conditions and physiological limits that rely on only a few parameters [49,50,52,55,57,58,64–66]. A more detailed overview of the different modeling approaches is provided in Appendix 1. Here, we focus particularly on the low-dimensional approaches which have been further linked with phenomenological descriptions [39,47,52,55,56,61,65] many of which took inspiration from a now semnial analysis by Scott *et al* [47] on growth and the adjustment of ribosome content in the presence of translation-inhibiting antibiotics. Several generalizations of the low-dimensional ribosomal allocation have further been proposed [55,61]. For example, Dourado *et al.* [61] were able to provide solutions of steady-state growth in a generalized manner when several protein classes are included. Though each model is distinct and often tailored to specific environments or growth conditions, their shared minimalism suggests that a single, self-consistent framework can by synthesized to rationalize the fundamental aspects of growth in both static and dynamic environments.

#### A simple allocation model describes translation-limited growth

We begin by formulating a simplified model of growth which follows the flow of mass from nutrients in the environment to biomass by building upon and extending the general logic of low-dimensional resource allocation models [39,47,50,52,55]. Specifically, we focus on the accumulation of *protein* biomass, as protein constitutes the majority of microbial dry mass [67,68] and peptide bond formation commonly accounts for $\approx 80\%$ of the cellular energy budget [33,69]. Furthermore, low-dimensional allocation models utilize a simplified representation of the proteome where proteins can be categorized into only a few functional classes [48,50,52,54,61]. In this work, we consider proteins to be either ribosomal (i.e. directly involved in peptide bond formation using charged-tRNAs), metabolic (i.e. enzymes catalyzing synthesis of charged-tRNA molecules from environmental nutrients), or being involved in all other biological processes (e.g. lipid synthesis, DNA replication, and chemotaxis) [47,48,50,52] (Supplementary Figure 1). Simple allocation models further do not distinguish between different cells but only consider the overall turnover of nutrients and biomass. To this end, we explicitly consider a well-mixed batch culture growth as reference scenario where the nutrients are considered to be in abundance. This low-dimensional view of living matter may at first seem like an unfair approximation, ignoring the decades of work interrogating the multitudinous biochemical and biophysical processes of cell-homeostasis and growth [48,51,59,70,71]. However, at least in nutrient replete conditions, many of these processes appear not to impose a fundamental limit on the rate of growth in the manner that protein synthesis does [33]. In Appendix 2, we discuss this along with other simplifications in more detail.

For constant allocation parameters $(\phi_R^*, \phi_M^*)$ a steady-state regime emerges from this system of differential equation. Particularly, the precursor concentration is stationary in time ($c_{pc} = c_{pc}^*$), meaning the rate of synthesis is exactly equal to the rate of consumption and dilution. Furthermore, the translation rate $\gamma(c_{pc}^*)$ is constant during steady-state growth and the mass-abundances of ribosomes and metabolic proteins are equivalent to the corresponding allocation parameters, e.g. $\frac{M_{Rb}}{M} \equiv \phi_{Rb}^*$. As a consequence, biomass is increasing exponentially $\frac{dM}{dt} = \lambda M$, with the growth rate $\lambda = \gamma(c_{pc}^*)\phi_{Rb}^*$. The emergence of a steady state and analytical solutions describing steady growth are further discussed in Supplementary Figures 2 and 3. Notably, dilution is important to obtain a steady state as has been highlighted previously by Giordano *et al.* [55] and Dourado *et al.* [61] but is often neglected (Appendix 3).

Fig. 1 (C) and (D) show how the steady-state growth rate $\lambda$ and translation rate $\gamma(c_{pc}^*)$ are dependent on the allocation towards ribosomes $\phi_{Rb}^*$. The figures also show the dependence on the metabolic rate $\nu_{max}$ which we here assert to be a proxy for the "quality" of the nutrients in the environment (with increasing $\nu_{max}$, less metabolic proteins are required to obtain the same synthesis of precursors). The non-monotonic dependence of the steady-state growth rate on the ribosome allocation and the metabolic rate poses a critical question: What biological mechanisms determine the allocation towards ribosomes in a particular environment and what criteria must be met for the allocation to ensure efficient growth?

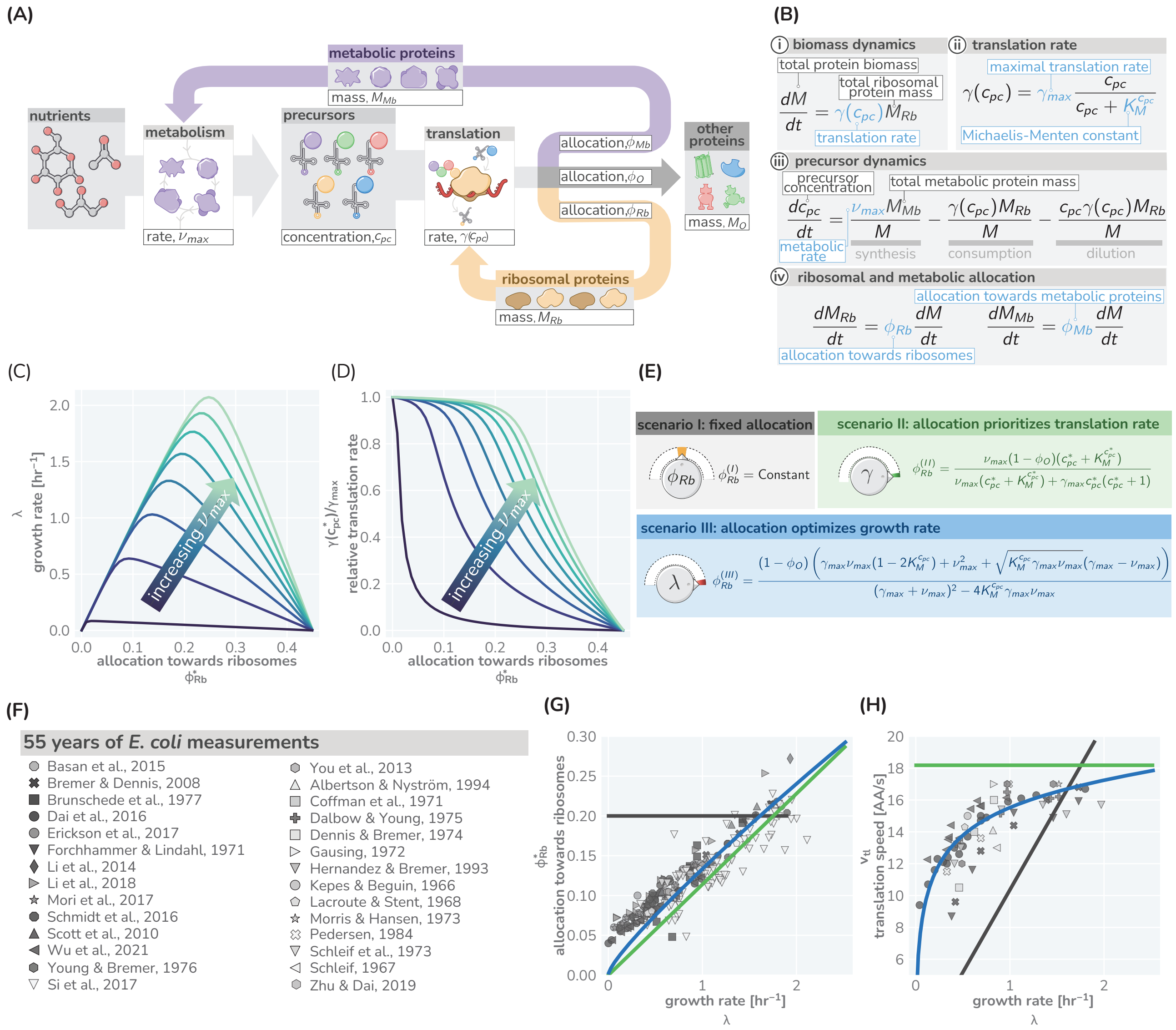

**Figure 1: A simple model of ribosomal allocation and hypothetical regulatory strategies.** (A) The flow of mass through the self-replicating system. Biomolecules and biosynthetic processes are shown as grey and white boxes, respectively. Nutrients in the environment passed through cellular metabolism to produce "precursor" molecules, here diagrammed as charged-tRNA molecules. These precursors are consumed through the process of translation to produce new protein biomass, either as metabolic proteins (purple arrow), ribosomal proteins (gold arrow), or "other" proteins (gray arrow). The mathematical symbols as used in the simplistic ribosomal allocation model are indicated. (B) Annotated equations of the model with key parameters highlighted in blue. An interactive figure where these equations can be numerically integrated is provided on paper website. The steady-state values of (C) the growth rate $\lambda$ and (D) the relative translation rate $\gamma(c_{pc}^*)/\gamma_{max}$, are plotted as functions of the allocation towards ribosomes for different metabolic rates (colored lines). (E) Analytical solutions for candidate scenarios for regulation of ribosomal allocation with fixed allocation, allocation to prioritize translation rate, and allocation to optimal growth rate highlighted in grey, green, and blue respectively. (F) A list of collated data sets of *E. coli* ribosomal allocation and translation speed measurements spanning 55 years of research. Details regarding these sources and method of data collation is provided in Table S1. A comparison of the observations with predicted growth-rate dependence of ribosomal allocation (G) and translation speeds (H) for the three allocation strategies. An interactive version of the panels allowing the free adjustment of parameters is available on the associated paper website (`cremerlab.github.io/flux_parity`).

#### Optimal Allocation Results From a Mutual Maximization of Translational and Metabolic Flux

To optimize the steady-state growth rate, cells must have some means of coordinating the flow of mass through metabolism and protein synthesis. In the ribosomal allocation model, this reduces to a regulatory mechanism in which the allocation parameters ($\phi_{Rb}$ and $\phi_{Mb}$) are dynamically adjusted such that the metabolic flux to provide new precursors ($\nu\phi_{Mb}$) and translational flux to make new proteins ($\gamma\phi_{Rb}$, equivalent to the steady-state growth rate $\lambda$) are not only equal, but are mutually maximized. Such regulation therefore requires a mechanism by which both the metabolic and translational flux can be simultaneously sensed.

Thus far, we have referred to the end-product of metabolism as ambiguous "precursors" which are used by ribosomes to create new proteins. In reality, these precursors are tRNAs charged with their cognate amino acids. One can think of metabolism as a two-step process where (i) an amino acid is synthesized from environmental nutrients and (ii) an amino acid is attached to the appropriate uncharged-tRNA. As we assume that nutrients are in excess in the environment, we make the approximation that the metabolic rate $\nu$ is dependent solely on the uncharged-tRNA concentration $\nu(\text{tRNA}^u)$. This enforces some level of regulation of metabolism; if the uncharged tRNA concentration is too low, the rate of metabolism slows and does not add to the already large pool of charged tRNA. But when charged-tRNA is available, translation occurs at a rate $\gamma(\text{tRNA}^c)$, forming new protein biomass and converting a charged-tRNA back to an uncharged state. This process is shown by grey arrows in Fig. 2(A).

To describe the state-dependent adjustment of the allocation parameters ($\phi_{Rb}$ and $\phi_{Mb}$), we further include in this feedback loop a regulatory system we term a "flux-parity regulator" [Fig. 2(A), red], which controls the allocation parameters in response to relative changes in the concentrations of the two tRNA species. Together, the arrows in Fig. 2 represents a more fine-grained view of a proteinaceous self replicating system, yet maintains much of the structural minimalism of the simple ribosomal allocation model without requiring explicit consideration of different types of amino acids [65], inclusion of their myriad synthesis pathways [60], or reliance on observed phenomenology [42].

The boxes and arrows of Fig. 2(A) can be mathematized to arrive at a handful of ordinary differential equations [Fig. 2(B)] structurally similar to those in Fig. 1(B). At the center of this model is the ansatz that the ribosomal

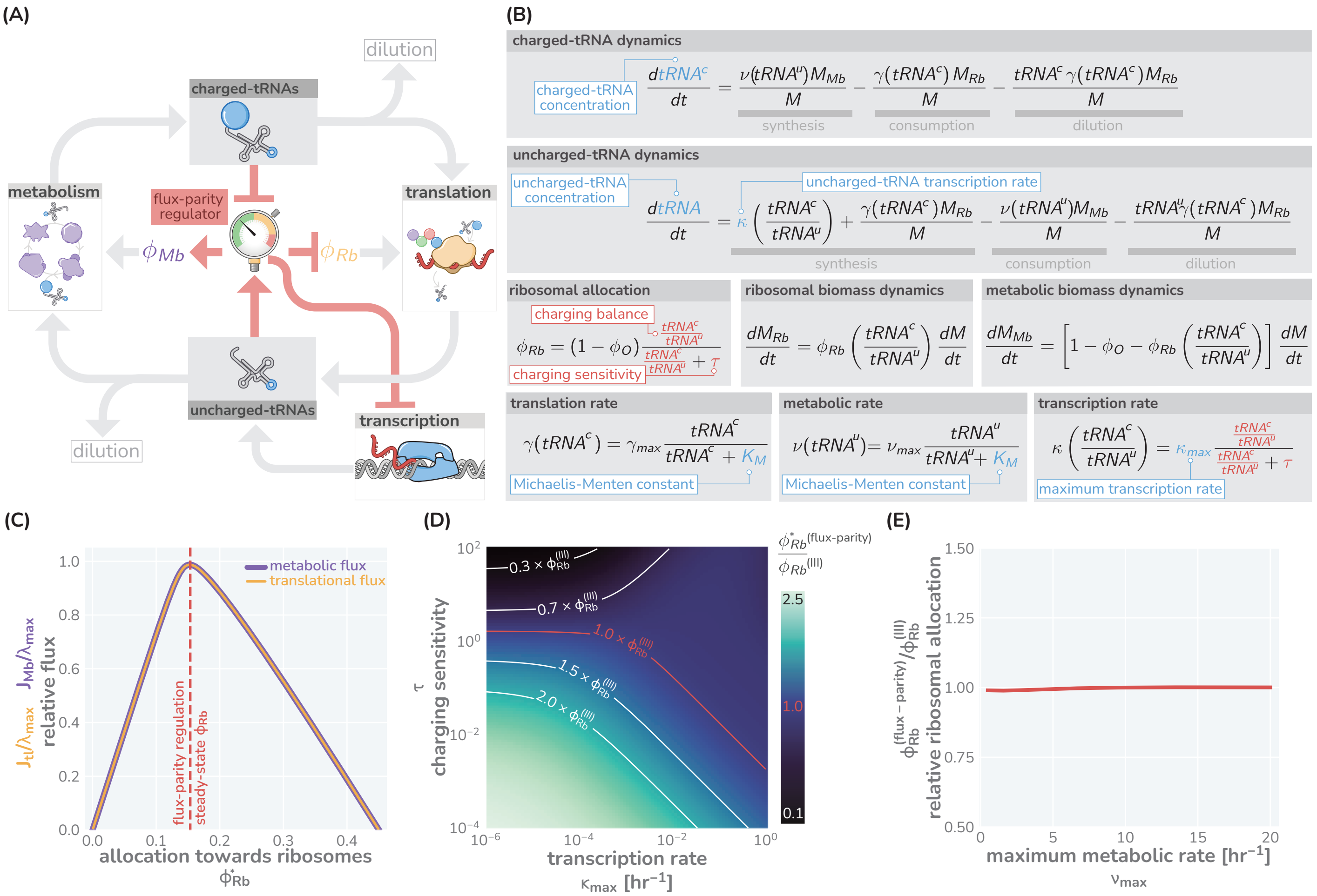


**Figure 2: The regulation of ribosome allocation via a flux-sensing mechanism.** (A) A circuit diagram of interactions between metabolic and translational fluxes with flux-parity regulatory connections highlighted in red. The fluxes are connected via a positive feedback loop through the generation of mutual starting materials (uncharged- or charged-tRNAs, respectively). The rates of each flux exhibit semi-autoregulatory behavior in that flux through each process reduces the standing pool of tRNAs. (B) The governing dynamics of the flux-parity regulatory circuit with key parameters highlighted in blue and flux-parity regulatory components highlighted in red. (C) The steady-state meabolic (purple) and translational (gold) fluxes plotted as a function of the ribosomal allocation under the simple allocation model. Vertical red line indicates the steady-state solution of the flux-parity model under physiological parameter regimes. (D) The steady-state allocation towards ribosomes emerging from flux-parity regulation ($\phi_{Rb}^{(flux-parity)}$) relative to the optimal allocation of the simplistic model ($\phi_{Rb}^{(III)}$) is shown across different parameter regimes for the charging sensitivity $\tau$ and the uncharged-tRNA transcription rate $\kappa$. Red contour demonstrate the plane of parameter space where the flux-parity regulatory circuit exactly matches the result of optimal allocation. (E) The relative allocation towards ribosomes ($\phi_{Rb}^{(flux-parity)}/\phi_{Rb}^{(III)}$) under physiological parameter regimes plotted as a function of the maximal metabolic rate, $\nu_{max}$.

allocation $\phi_{Rb}$ is dependent on the ratio of charged- and uncharged-tRNA pools and has the form

$$\phi_{Rb}\left(\frac{\text{tRNA}^c}{\text{tRNA}^u}\right) = (1-\phi_O)\frac{\frac{\text{tRNA}^c}{\text{tRNA}^u}}{\frac{\text{tRNA}^c}{\text{tRNA}^u}+\tau}, \tag{1}$$

where the ratio $\frac{\text{tRNA}^c}{\text{tRNA}^u}$ represents the "charging balance" of the tRNA and $\tau$ is a dimensionless "sensitivity parameter" which defines the charging balance at which the allocation towards ribosomes is half-maximal. Additionally, we make the assertion that the synthesis rate of new uncharged-tRNA via transcription $\kappa$ is coregulated with ribosomal proteins [77, 78] and has a similar form of

$$\kappa\left(\frac{\text{tRNA}^c}{\text{tRNA}^u}\right) = \kappa_{max}\frac{\frac{\text{tRNA}^c}{\text{tRNA}^u}}{\frac{\text{tRNA}^c}{\text{tRNA}^u}+\tau}, \tag{2}$$

where $\kappa_{max}$ represents the maximal rate of tRNA transcription relative to the total biomass.

$$\frac{[\text{ppGpp}]}{[\text{ppGpp}]_0} = \frac{\frac{\text{tRNA}^u}{\text{tRNA}^c}}{\frac{\text{tRNA}^u_0}{\text{tRNA}^c_0}}. \tag{4}$$

To test this, we compiled and rescaled ppGpp measurements of *E. coli* across a range of growth rates from various literature sources [Fig. 3(C)]. The quantitative agreement between the scaling predicted by Eq. 4 and the experimental measurements strongly suggests that ppGpp assumes the role of a flux-sensor and enforces optimal allocation through the discussed flux-parity mechanism.

#### The Flux-Parity Allocation Model Predicts E. coli Growth Behavior In and Out Of Steady-State

We find that the flux-parity allocation model is extremely versatile and allows us to quantitatively describe aspects of microbial growth in and out of steady-state and under various physiological stresses and external perturbations with the same core set of parameter. Here, we demonstrate this versatility by comparing predictions to data for four particular examples *using the same self-consistent set of parameters we have used thus far* (Table S1). First, we examine the influence of translation-targeting antibiotics like chloramphenicol [Fig. 3(D)] on steady-state growth in different growth media [39, 47]. By incorporating a mathematical description of ribosome inactivation via binding

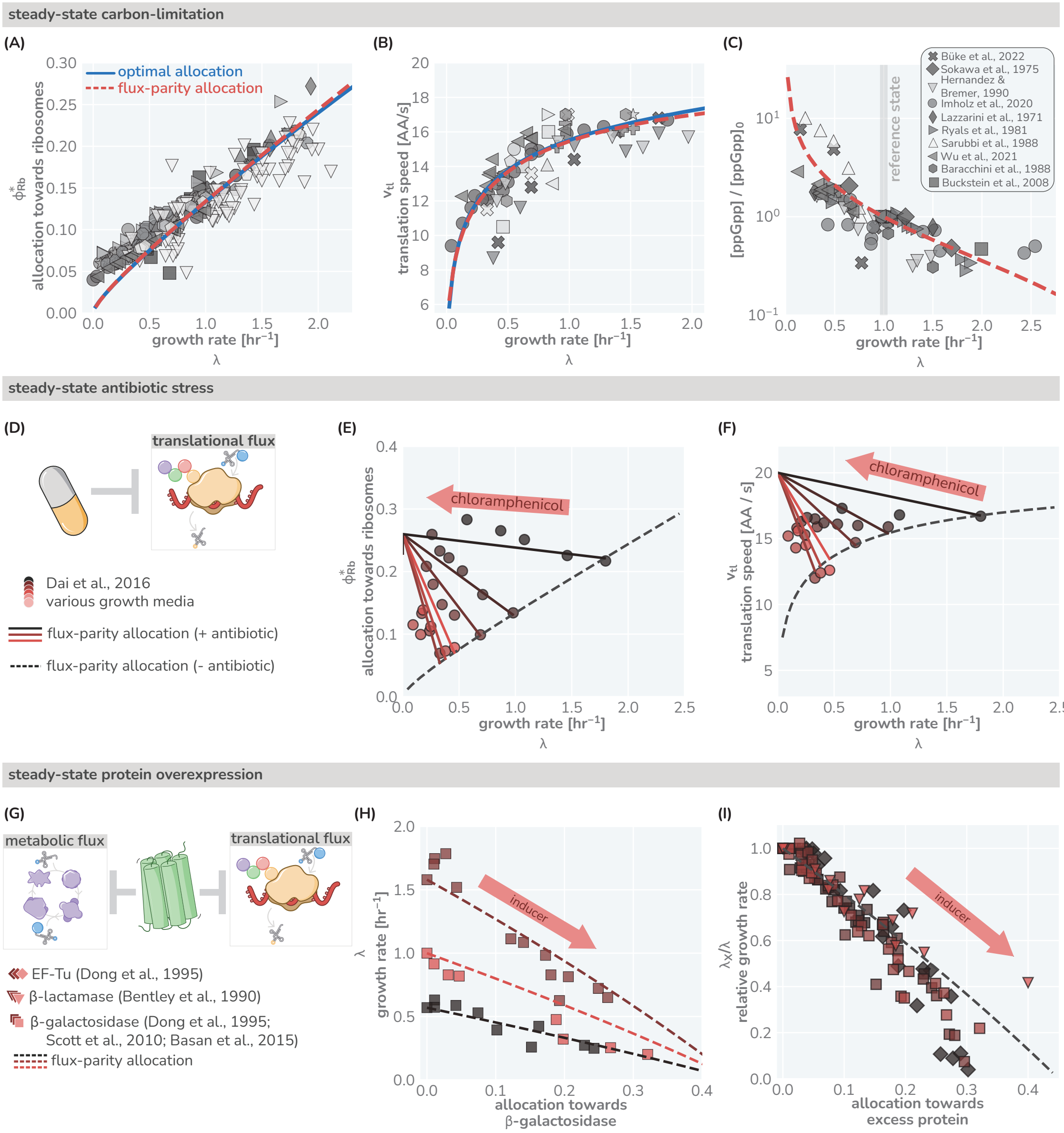


**Figure 3: The predictive power of flux-parity regulation in steady-state.** Measurements of the (A) ribosomal allocation and the (B) translation rate are plotted alongside the steady-state behavior of the flux-parity regulatory circuit (red dashed line) and the optimal behavior of scenario III (solid blue line). Points and markers are the same as those shown in Fig. 1(F). (C) Measurements of intracellular ppGpp concentrations relative to a reference condition ($\lambda_0 \approx 1\ \text{hr}^{-1}$) are plotted as a function of growth rate alongside the prediction emergent from the flux-parity regulatory circuit (red dashed line). (D-F) Inhibition of ribosome activity via antibiotic modeled repression of translational flux. Plots show comparison with data for different media (red shades) with the flux-parity model predictions (dashed lines). (G-I) Inhibition of metabolic and translational fluxes through excess gene expression. (H) shows data where $\beta$-galactosidase is expressed at different levels. Different shades of red correspond to different growth media. Right-hand panel shows collapse of the growth rates of overexpression of $\beta$-galactosidase (squares), $\beta$-lactamase (inverted triangles), and EF-Tu (diamonds) relative to the wild-type growth rate in different media conditions. The same set of model parameters listed in Table S2 has been used to generate the predictions.

Growth emerges as in previous allocation models [47, 50, 55] as a consequence of protein synthesis and the

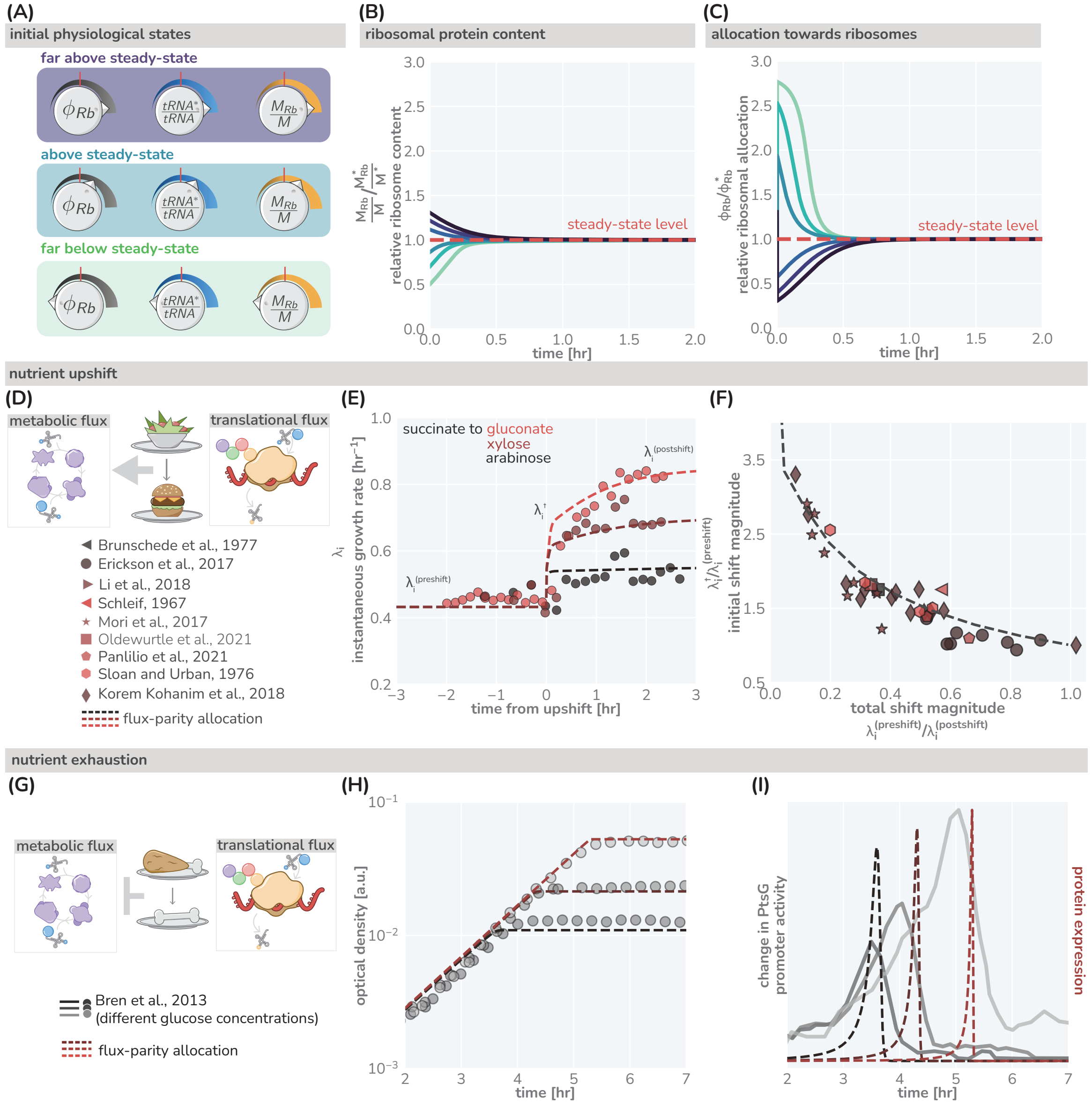


**Figure 4: The predictive power of flux-parity regulation out of steady-state.** (A) Hypothetical initial configurations of model parameters and variables before begining numerical integration. (B) The equilibration of the ribosomal protein content ($M_{Rb}/M$). (C) Dynamic adjustment of the ribosomal allocation parameter in response to the new environment. Green and purple colored lines correspond to the initial conditions of the culture from well above to well below the steady-state values, respectively. Dashed red line indicates the steady-state solution. (D-E) Nutrient upshifts with increased metabolic flux. (E) The instantaneous growth rate $\lambda_i$ for shifts from succinate to gluconate (bright red), xylose (dark red), or arabinose (black) [57]. (F) Collapse of instantaneous growth rate measurements immediately after the shift (relative to the preshift-growth rate) as a function of the total shift magnitude. (G-I) Exhaustion of nutrients in the environment yields a decrease in the metabolic flux, promoting expression of more metabolic proteins. (H) Growth curve measurements in media with different starting concentrations of glucose (0.22 mM, 0.44 mM, and 1.1 mM glucose from light to dark, respectively) overlaid with flux-parity predictions. (I) The change in total metabolic protein synthesis in the flux-parity model (dashed lines) overlaid with the change in expression of a fluorescent reporter from a PtsG promoter (solid lines).

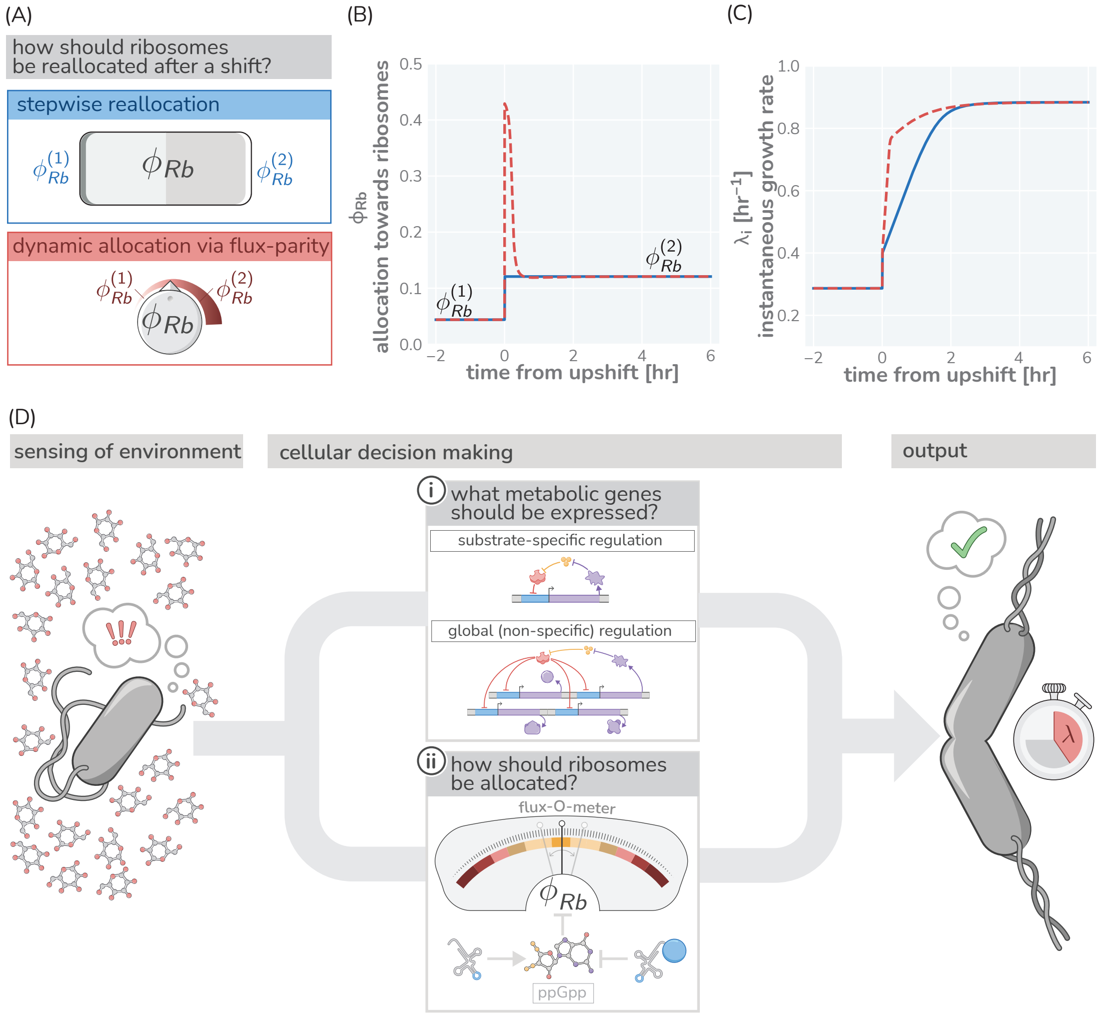


**Figure 5: Flux parity allocation as a strategy to adapt to fluctuating conditions.** (A) Ribosome reallocation strategies upon a nutrient upshift. After a nutrient upshift, cells either dynamically reallocate their ribosomes given flux-parity regulation (top, red) or they undergo stepwise reallocation from one steady-state value to the next (bottom, blue). (B) The allocation dynamics for both strategies in response to a nutrient upshift. (C) The instantaneous growth rate for both strategies over the course of the shift. Dashed red and solid blue lines correspond to model predictions for optimal allocation and flux-parity regulation, respectively. (D) Cellular decision making in fluctuating environments. Upon sensing features of the environment, cells undergo a two-component decision making protocol defining what metabolic genes should be expressed (top) and how the allocation towards ribosomes should be adjusted to maintain flux-parity. The combination of these processes yield an increase of biomass at a given characteristic growth rate.

$$v_{tl}(c_{pc}) = v_{tl}^{max}\left(\frac{c_{pc}}{c_{pc} + K_M^{c_{pc}}}\right), \tag{6}$$

where $K_M^{c_{pc}}$ is a Michaelis-Menten constant with the maximum speed $v_{tl}^{max}$ only observed for $c_{pc} \gg K_M^{c_{pc}}$. The number of ribosomes $N_{Rb}$ can be approximated given knowledge of the total mass of ribosomal proteins $M_{Rb}$ and the proteinaceous mass of a single ribosome $m_{Rb}$ via $N_{Rb} \approx M_{Rb}/m_{Rb}$(more details in Appendix 9). The increase in protein biomass (Eq. 5) is thus

$$\frac{dM}{dt} = v_{tl}(c_{pc})\frac{M_{Rb}}{m_{Rb}} \equiv \gamma(c_{pc})M_{Rb}. \tag{7}$$

The *translation rate* $\gamma(c_{pc}) \equiv v_{tl}(c_{pc})/m_{Rb}$ describes the rate at which ribosomes generate new protein.

The maximal translation rate $\gamma_{max} \equiv v_{tl}^{max}/m_{Rb}$ imposes a firm upper limit [33, 72, 73] of how rapidly biomass can accumulate, unrealistically assuming the system would consist of only ribosomes translating at maximum rate. Notably, however, this upper limit is not much faster than the fastest growth observed, highlighting the importance of protein synthesis in defining the timescale of growth. For example, the maximal translation rate for *E. coli* is $\approx 10$ hr$^{-1}$ and thus only $\approx 4$ times higher than the growth rates in rich LB media ($\lambda \approx 2.5$ hr$^{-1}$). Including the synthesis of rRNA, another major component of the cellular dry mass, lowers this theoretical limit only marginally, further supporting our sole consideration of protein synthesis in defining growth (Appendix 10). The difference between measured growth rates and the theoretical limits can be mostly attributed to the synthesis of metabolic proteins which generate the precursors required for protein synthesis, which we consider next.

**Synthesis of Precursors**

Microbial cells are generally capable of synthesizing precursors from nutrients available in the environment, such as sugars or organic acids. This synthesis is undertaken by a diverse array of metabolic proteins ranging from those which transport nutrients across the cell membrane, to the enzymes involved in energy generation (such as those of fermentation or respiration), and the enzymes providing the building blocks for protein synthesis (such as those involved in the synthesis of amino acids). While these enzymes vary in their abundance and kinetics, we group them all into single set of metabolic proteins with a mass $M_{Mb}$ which cooperate to synthesize the collective pool of precursors from nutrients required for protein synthesis. We make the approximation that these metabolic proteins generate precursors at an effective *metabolic rate* $\nu$. In general, this rate depends on the concentration of nutrients $c_{nt}$ in the environment. This relation is canonically described by a Monod (Michaelis-Menten) relation

$$\nu(c_{nt}) = \nu^{max}\left(\frac{c_{nt}}{c_{nt} + K_M^{c_{nt}}}\right), \tag{8}$$

where $\nu_{max}$ is the maximum metabolic rate describing how fast the metabolic proteins can synthesize precursors, and $K_M^{c_{nt}}$ is the Monod constant describing the concentration below which nutrient utilization slows [90]. Novel precursors are thus supplied with a total rate of $\nu(c_{nt})M_{Mb}$ and consumed via protein synthesis at a rate $\gamma(c_{pc})M_{Rb}$. Translation relies on precursors and, as introduced above, the translation rate $\gamma(c_{pc})M_{Rb}$ thus depends on the concentration of precursors in the cell, $c_{pc}$. As we do not explicitly model cell division, we here approximate this cellular concentration as the relative mass abundance of precursors to total protein biomass. This approximation is justified by the observation that cellular mass density and total protein content is approximately constant across a wide range of conditions [33, 91, 92]. The dynamics of precursor concentration follows from the balance of synthesis, consumption, and dilution as the total biomass grows:

$$\frac{dc_{pc}}{dt} = \overbrace{\frac{\nu(c_{nt})M_{Mb}}{M}}^{\text{production via metabolism}} - \underbrace{\frac{\gamma(c_{pc})M_{Rb}}{M}}_{\text{consumption via protein synthesis}} - \overbrace{\frac{c_{pc}\gamma(c_{pc})M_{Rb}}{M}}^{\text{dilution via growth}}. \tag{9}$$

While the dilution term is often assumed to be negligible, this term is critical to describe growth and derive analytical expressions (detailed discussion in Appendix 3). Furthermore, we note that the precursor concentration is defined such that the consumption of one precursor yields the addition of one amino acid to the biomass $M$. As we measure proteins in units of amino acids there is thus no conversion factor needed when describing the consumption of precursors by protein synthesis.

**Simplification of saturating nutrients**

The introduced dynamics simplifies when the nutrient concentration in the environment $c_{nt}$ well exceeds the Monod constant $K_M^{c_{nt}}$ as $\nu(c_{nt})$ simplifies to $\nu_{max}$. Steady growth for which biomass increases exponentially readily emerges. This is the scenario we focus on in in the first half of this work. It should be noted however, that biologically such a scenario can only be realized temporarily as the nutrient supply required by the exponentially growing biomass can only be sustained by the environment for a limited amount of time. In general, the nutrient levels vary.

##### Consumption of Nutrients in batch culture growth

The synthesis of novel precursors relies on the availability of nutrients which changes depending on the environment. In Supplementary Figure 2 we consider specifically a "batch culture" scenario in which nutrients are provided only at the beginning of growth and are never replenished. Therefore, growth of the culture continues until all of the nutrients have been consumed. The concentration of nutrients in the environment is thus given as

$$\frac{dc_{nt}}{dt} = -\frac{\nu(c_{nt})M_{Mb}}{Y}, \tag{10}$$

where $Y$ is the yield coefficient which describes how many nutrient molecules are needed to produce one unit of precursors.

##### Ribosomal Allocation of Protein Synthesis

As final step of the model definition, we must describe how cells direct their protein synthesis towards making ribosomes, metabolic proteins, or all other proteins that make up the cell [colored arrows in Fig. 1(A)]. We do so by introducing three *allocation parameters* $\phi_{Rb}$, $\phi_{Mb}$, and $\phi_O$ (such that $\phi_{Rb} + \phi_{Mb} + \phi_O = 1$) which define how novel protein synthesis is partitioned among these categories:

$$\frac{dM_{Rb}}{dt} = \phi_{Rb}\frac{dM}{dt}\ ;\ \frac{dM_{Mb}}{dt} = \phi_{Mb}\frac{dM}{dt}\ ;\ \frac{dM_O}{dt} = \phi_O\frac{dM}{dt}. \tag{11}$$

These equations are summarized in Fig.1(B) and Fig. S3 and define the accumulation of biomass, from nutrient uptake to protein synthesis.

#### Approximating concentration via relative abundance

In addition to maintaining the *total* macromolecular densities, cells also maintain an approximately constant protein density [93]. This observation allows for a major simplification when formulating the allocation model, namely the approximation of concentrations as relative mass abundances. The rate $\gamma$ at which ribosomes can synthesize protein is dependent on the abundance of precursors, $c_{pc}$, in the cell. To compute the concentration and/or density in typical units (e.g. $\mu$M, or mass / volume), we would require some measure of the total cellular volume, $V_{cell}$, such that the concentration follows

$$c_{pc} = \frac{M_{pc}}{V_{cell}}, \tag{12}$$

with $M_{pc}$ denoting the total mass of the precursor pool. By making the experimentally-supported assertion that the protein density $\rho$ is constant, we can say that

$$\rho = \frac{M}{V_{cell}} = \text{Constant}, \tag{13}$$

where $M$ is the total protein biomass. Thus, the total cellular volume $V_{cell}$ can be computed as

$$V_{cell} = \frac{M}{\rho}. \tag{14}$$

Plugging this result into Eq. 12, we arrive at the approximation

$$c_{pc} = \rho\frac{M_{pc}}{M} \approx \frac{M_{pc}}{M}. \tag{15}$$

In this work, we neglect $\rho$ as a multiplicative constant, and treat $c_{pc}$ as being dimensionless. We direct the reader to Refs. [47] and [94] for a further discussion of the conversion between concentration and relative abundance.

#### Derivation of Analytical Expressions

In the first section of this work, we present several analytical relations pertinent to steady-state growth. These relations follow from the simple allocation model and describe (i) how the growth rate depends on model parameters (Fig. 1C) and (ii) how ribosome content depends on other model parameters for the three different regulation scenarios we discuss (Fig. 1D and E). Here, we introduce a step-by-step derivation of these expressions.

##### Deriving the Steady-State Growth Rate

We begin with deriving an expression for the steady-state growth rate $\lambda$ which is similar to previous approaches taken by Giordano *et al* [55] and Dourado *et al* [61]. As discussed in Supplementary Figure 2, steady-state conditions are satisfied when two conditions are met. First, the dynamics of the precursor concentration is constant (i.e. $\frac{dc_{pc}}{dt} = 0$) and the composition of the proteome matches the allocation parameters (i.e. $\frac{M^*_{Rb}}{M^*} = \phi^*_{Rb}$ and $\frac{M^*_{Mb}}{M^*} = \phi^*_{Mb}$). Furthermore, we assume that in steady-state growth, the concentration of nutrients in the environment is saturating ($c_{nt} \gg K_M^{c_{nt}}$), meaning that $\nu(c_{nt}) \approx \nu_{max}$. With these conditions satisfied, we can rewrite Eq. 9 as

$$\frac{dc_{pc}}{dt} = \nu_{max}\phi_{Mb} - \gamma(c^*_{pc})\phi_{Rb} - c_{pc}\gamma(c^*_{pc})\phi_{Rb} = 0, \tag{16}$$

where $c^*_{pc}$ is the steady-state precursor concentration.

Noting that in steady-state conditions, the total biomass increases exponentially at a rate $\lambda \equiv \gamma(c_{pc})\phi^*_{Rb}$, Eq. 16 can be simplified to

$$\frac{dc_{pc}}{dt} = \nu_{max}\phi^*_{Mb} - \lambda(1 + c_{pc}) = 0. \tag{17}$$

We can therefore solve for the steady-state precursor concentration $c^*_{pc}$ to yield

$$c^*_{pc} = \frac{\nu_{max}\phi^*_{Mb}}{\lambda} - 1. \tag{18}$$

Assuming a Michaelis-Menten form for the translation rate $\gamma(c^*_{pc})$, we can now define it as a function of the growth rate $\lambda$ as

$$\gamma(c^*_{pc}) = \frac{\gamma_{max}}{1 + \frac{K_M^{c_{pc}}}{c_{pc}}} = \frac{\gamma_{max}}{1 + \frac{K_M^{c_{pc}}\lambda}{\nu_{max}\phi^*_{Mb} - \lambda}}. \tag{19}$$

Knowing that the growth rate $\lambda \equiv \gamma(c_{pc^*})\phi^*_{Rb}$, and $\phi^*_{Mb} = 1 - \phi^*_{Rb} - \phi^*_O$, we say that

$$\lambda = \frac{\gamma_{max}\phi^*_{Rb}}{1 + \frac{K_M^{c_{pc}}\lambda}{\nu_{max}(1 - \phi^*_{Rb} - \phi^*_O) - \lambda}}. \tag{20}$$

This can be algebraically manipulated to yield a quadratic equation of the form

$$\lambda^2\left(1 - K_M^{c_{pc}}\right) + \lambda\left(\nu_{max}(1 - \phi^*_{Rb} - \phi^*_O) + \gamma_{max}\phi^*_{Rb}\right) - \gamma_{max}\phi^*_{Rb}\nu_{max}(1 - \phi^*_{Rb} - \phi^*_{Mb}) = 0, \tag{21}$$

which has one positive root of

$$\lambda = \frac{\nu_{max}(1 - \phi^*_{Rb} - \phi^*_O) + \gamma_{max}\phi^*_{Rb} - \sqrt{(\nu_{max}(1 - \phi^*_{Rb} - \phi^*_O) + \gamma_{max}\phi^*_{Rb})^2 - 4(1 - K_M^{c_{pc}})\gamma_{max}\phi^*_{Rb}\nu_{max}(1 - \phi^*_{Rb} - \phi^*_O)}}{2\left(1 - K_M^{c_{pc}}\right)}. \tag{22}$$

For notational simplicity, we can define the maximum metabolic output and the maximum translational output as $\mathsf{N} = \nu_{max}(1 - \phi_{Rb} - \phi_O)$ and $\Gamma = \gamma_{max}\phi_{Rb}$, respectively, and substitute them into Eq. 22 to generate

$$\lambda = \frac{\mathsf{N} + \Gamma - \sqrt{(\mathsf{N} + \Gamma)^2 - 4\left(1 - K_M^{c_{pc}}\right)\mathsf{N}\Gamma}}{2\left(1 - K_M^{c_{pc}}\right)}, \tag{23}$$

.

##### Defining $\phi_{Rb}$ For Scenarios II and III

In Fig. 1(D), we provide a description of three plausible regulatory scenarios microbes may employ to regulate their ribosomal content. Scenario I assumes just a constant, arbitrary allocation parameter $\phi_{Rb} \in [0, 1 - \phi_O]$. Here, we provide a short derivation for the more complicated relations describing ribosomal content under scenarios II and III.

##### Scenario II: Constant Translation Rate

The second regulatory scenario assumes that the ribosomal content is adjusted to maintain a specific standing concentration of precursors, which we denote as $c_{pc}^*$. Noting that the growth rate $\lambda \equiv \gamma(c_{pc}^*)\phi_{Rb}^*$, we can restate Eq. 18 in the form

$$c_{pc}^* = \frac{\nu_{max}(1 - \phi_O^* - \phi_{Rb}^*)(c_{pc}^* + K_M^{c_{pc}})}{c_{pc}^* \gamma_{max} \phi_{Rb}^*}. \tag{24}$$

Some algebraic rearrangement allows us to solve for $\phi_{Rb}^*$, yielding

$$\phi_{Rb} = \frac{(1 - \phi_O^*)\nu_{max}\left(c_{pc}^* + K_M^{c_{pc}}\right)}{\nu_{max}\left(c_{pc}^* + K_M^{c_{pc}}\right) + \gamma_{max} c_{pc}^* \left(c_{pc}^* + 1\right)}. \tag{25}$$

This expression is equivalent to that shown for scenario II in Fig. 3 of the main text. In evaluating this scenario, we considered the regime in which precursors were in abundance, meaning $c_{pc}^* \gg K_M^{c_{pc}^*}$. Under this regime, Eq. 25 simplifies further to

$$\phi_{Rb}^* \approx \frac{(1 - \phi_O^*)\nu_{max}}{\gamma_{max}\left(c_{pc}^* + 1\right) + \nu_{max}}. \tag{26}$$

This represents a strategy where the cell adjusts $\phi_{Rb}^*$ to maintain a translation rate very close to $\gamma_{max}$.

##### Scenario III: Optimal Allocation

In this work, we define the optimal allocation of ribosomes $\phi_{Rb}^*$ to be that which maximizes the growth rate in a given environment and at a given metabolic state. To determine the optimal $\phi_{Rb}^*$, we can differentiate Eq. 22 with respect to $\phi_{Rb}^*$ to yield the cumbersome expression

$$\frac{\partial \lambda}{\partial \phi_{Rb}^*} = \frac{1}{2\left(1 + K_M^{c_{pc}}\right)} \times \tag{27}$$

$$\left[\gamma_{max} - \nu_{max} - \frac{2\gamma_{max}\nu_{max}\left(1 - K_M^{c_{pc}}\right)\left(2\phi_{Rb}^* + \phi_O^* - 1\right) + \left(\gamma_{max} - \nu_{max}\right)\left(\gamma_{max}\phi_{Rb}^* + \nu_{max}\left(1 - \phi_O^* - \phi_{Rb}^*\right)\right)}{\sqrt{\left(\gamma_{max}\phi_{Rb}^* + \nu_{max}\left(1 - \phi_O^* - \phi_{Rb}^*\right)\right)^2 - 4\left(1 - K_M^{c_{pc}}\right)\gamma_{max}\nu_{max}\phi_{Rb}^*\left(1 - \phi_O^* - \phi_{Rb}^*\right)}}\right]. \tag{28}$$

Setting this expression equal to zero and solving for $\phi_{Rb}$ results in

$$\phi_{Rb} = \frac{(1 - \phi_O^*)\left(\gamma_{max}\nu_{max}\left(1 - 2K_M^{c_{pc}}\right) + \nu_{max}^2 + \sqrt{K_M^{c_{pc}}\gamma_{max}\nu_{max}}(\gamma_{max} - \nu_{max})\right)}{(\gamma_{max} + \nu_{max})^2 - 4K_M^{c_{pc}}\gamma_{max}\nu_{max}} \tag{29}$$

$$\frac{dtRNA^c}{dt} = \overbrace{\frac{\nu(tRNA^u)M_{Mb}}{M}}^{\text{generation via metabolism}} - \underbrace{\frac{\gamma(tRNA^c)M_{Rb}}{M}}_{\text{consumption via protein synthesis}} - \overbrace{\frac{tRNA^c\gamma(tRNA^c)M_{Rb}}{M}}^{\text{reduction via dilution}}. \tag{30}$$

The dynamics for the pool of uncharged-tRNAs can be constructed in a similar manner, with the caveat that the generation of new uncharged-tRNAs occurs from both protein synthesis (converting one charged-tRNA into one uncharged-$tRNA^u$) and from transcription of the individual tRNA genes. We consider the latter to occur at a rate $\kappa$, which has dimensions of concentration per unit time. Using the same logic of mapping the productive and consumptive processes, we can enumerate the dynamics of the uncharged-tRNA pool as

$$\frac{dtRNA^u}{dt} = \overbrace{\kappa}^{\text{production via transcription}} + \underbrace{\frac{\gamma(tRNA^c)M_{Rb}}{M}}_{\text{occurance via protein synthesis}} - \overbrace{\frac{\nu(tRNA^u)M_{Mb}}{M}}^{\text{consumption via metabolism}} - \underbrace{\frac{tRNA\gamma(tRNA^c)M_{Rb}}{M}}_{\text{reduction via dilution}}. \tag{31}$$

$$\phi_{Rb}\left(\frac{tRNA^c}{tRNA^u}\right) = (1-\phi_O)\frac{\frac{tRNA^c}{tRNA^u}}{\frac{tRNA^c}{tRNA^u}+\tau}. \tag{34}$$

Here, the parameter $\tau$ represents the value of the charged- to uncharged-tRNA ratio where $\phi_{Rb}$ is at its half-maximal value. The maximal value itself depends on the magnitude of $\phi_O$, the allocation towards other proteins, which we are considering to be independent of ppGpp; $\phi_{Rb}^{(max)} = 1-\phi_O$.

The transcription of tRNA genes towards novel tRNA synthesis has also been shown to be regulated with ppGpp, appearing to closely match the regulatory behavior of ribosomal proteins [96]. We therefore model that the tRNA synthesis rate $\kappa$ [introduced in Eq. (31)] is similarly modulated by the charged- to uncharged-tRNA ratio,

$$\kappa\left(\frac{tRNA^c}{tRNA^u}\right) = \kappa_{\text{max}}\frac{\frac{tRNA^c}{tRNA^u}}{\frac{tRNA^c}{tRNA^u}+\tau}. \tag{35}$$

Here, $\kappa_{max}$ is the rate of tRNA transcription when all tRNA genes are fully saturated with RNA polymerase in rich growth conditions where gene dosage is high. Finally, we must establish functional forms for the tRNA dependencies on the metabolic and translation rate. Simple biochemical assumptions permit a formulation of a Michaelis-Menten function for each rate. Noting that the translation rate $\gamma$ is defined as $\gamma \equiv \frac{v_{tl}}{m_{Rb}}$, where $v_{tl}$ is the translation speed and $m_{Rb}$ is the proteinaceous mass of a single ribosome, we take $\gamma(tRNA^c)$ to be of the form

$$\gamma(tRNA^c) = \frac{v_{tl}^{(max)}}{m_{Rb}}\frac{tRNA^c}{tRNA^c + K_M^{(tRNA^c)}}, \tag{36}$$

where $v_{tl}^{(max)}$ is the maximum translation speed and $K_M^{(tRNA^c)}$ is the Michaelis-Menten constant. A similar argument can be made for the dependence of the metabolic rate $\nu$ on the uncharged-tRNA concentration,

$$\nu(tRNA^u) = \nu_{max}\frac{tRNA^u}{tRNA^u + K_M^{(tRNA^u)}}, \tag{37}$$

with $K_M^{(tRNA^u)}$ being another Michaelis-Menten constant. Together, Equations 30 through 37 mathematically describe a model for ppGpp-dependent regulation of translational and metabolic fluxes.

In principle, an analytical solution for this system of ODEs can be found, though it precludes evaluation by hand and is computationally intensive. While we do not solve this system of ODEs analytically here, we can numerically integrate them to sufficiently approximate the steady-state behavior. Depending on the choice of parameter values, such an approach can yield an allocation scenario nearly indistinguishable from that of the optimal allocation scenario (scenario III) of the simple model [Fig. 1(G and H)].

##### Optimal allocation emerges from flux-parity regulation

While the previous section lays out the mathematics of the flux-parity model, we now discuss how this regulation scheme can lead to an optimal allocation. Towards this goal we first discuss in more detail what we mean when we say 'flux-parity'. As described in the main text, we define flux-parity as a balance *and* mutual maximization of (i) the flux of uncharged-tRNAs through metabolism (termed the *metabolic flux* $J_{Mb}$) and (ii) the flux of charged-tRNAs through protein synthesis (termed the *translational flux* $J_{Tl}$). To demonstrate this point, assume that we can decouple the dependence of the allocation parameter $\phi_{Rb}$ from the ratio of charged- to uncharged-tRNAs. Mathematically speaking, we can define the metabolic flux as the collective action of metabolic proteins,

$$J_{Mb} = \nu(tRNA^u)\phi_{Mb} = \frac{\nu_{max} tRNA\,(1 - \phi_O - \phi_{Rb})}{tRNA^u + K_M^{tRNA^u}}. \tag{38}$$

Similarly, we can state that the translational flux is the collective action of ribosomal proteins,

$$J_{Tl} = \frac{\gamma_{max} tRNA^c \phi_{Rb}}{tRNA^c + K_M^{tRNA^c}} \tag{39}$$

$$[ppGpp] \propto P_{\text{bound}}^{(\text{uncharged})} \approx \frac{tRNA^u}{tRNA^c + tRNA^u}, \tag{40}$$

where $tRNA^c$ and $tRNA^u$ represent the absolute concentrations of charged and uncharged species, respectively. If the ppGpp concentration is inversely proportional to the allocation towards ribosomes, we can similarly make the argument that the ribosomal allocation $\phi_{Rb}$ will be proportional to the probability of a ribosome being bound to a *charged-tRNA*,

$$\phi_{Rb} = (1 - \phi_O) P_{\text{bound}}^{(\text{charged})} = (1 - \phi_O) \frac{tRNA^c}{tRNA^u + tRNA^c}. \tag{41}$$

This equation mechanistically operates in a similar way as Eq. 34 – the allocation towards ribosomes depends on the relative amounts of charged- and uncharged-tRNAs. In the extreme limit where the total concentration of tRNA is fixed (for which there is conflicting evidence [55, 65, 77, 78, 93]), Eq. 41 and Eq. 34 are mathematically equivalent. However, the predicted scaling dependence of ppGpp takes a different form.

In the main text, we noted that the concentration of ppGpp relative to a reference growth rate $\frac{[ppGpp]}{[ppGpp]_0}$ is equivalent to the inverse ratio of the charging balances. Under the ansatz that the ppGpp concentration is depending on the uncharged-tRNA binding probability, this relation takes the form

$$\frac{[ppGpp]}{[ppGpp]_0} = \frac{P_{\text{bound}}^{(\text{uncharged})}}{P_{\text{bound}_0}^{(\text{uncharged})}} = \frac{1 + \frac{tRNA_0^c}{tRNA_0}}{1 + \frac{tRNA^c}{tRNA^u}}, \tag{42}$$

where the subscript 0 denotes the reference state value. This distinction, coupled with experimental measurements of the relative ppGpp concentrations, allows us to test the validity of the two assumed forms for $\phi_{Rb}$.

i Fig. S6 shows the predictive capacity of these two ansatzes with the simple binding (Eq. 41) and ratiometric (Eq. 33) predictions shown in solid-blue and dashed-red lines, respectively. While both of these assumptions are capable of predicting the scaling of the ribosome content and translation speed with quantitative equivalence, there is a distinct difference in the predicted behavior of the relative ppGpp concentrations. The simple binding ansatz predicts a significantly shallower dependence on the growth rate than is observed in the data and in the ratiometric prediction. Thus, it appears that relating [ppGpp] to the ratio of uncharged- to charged-tRNA concentrations accurately captures the behavior of *E. coli*, though there remain gaps in our understanding of this relationship at a biochemical level.

#### Incorporating effects of ribosome-targeting antibiotics

To extend the flux-parity allocation model and incorporate the effects of antibiotic treatment, we must consider the mechanism of action of the antibiotic, specifically chloramphenicol. Chloramphenicol is a bacteriostatic antibiotic with tightly, but reversibly, binds to the ribosome. Once bound, the ribosome is unable to resume translation until chloramphenicol dissociates. Thus, we can model the effect of this drug by enumerating the probability that chloramphenicol is bound to a ribosome $P_{\text{bound}}$ at a given concentration $c_{cm}$. Mathematically, this can be stated as

$$P_{\text{bound}} = \frac{c_{cm}}{c_{cm} + K_D^{cm}}, \tag{43}$$

where $K_D^{cm}$ is an effective dissociation constant of chloramphenicol to a unit of ribosomal mass accounting for kinetics transport and ribosome binding. We can then say that the probability of a ribosome being active is equal to the probability of a ribosome being *unbound*,

$$P_{\text{active}} = 1 - P_{\text{bound}} = 1 - \frac{c_{cm}}{c_{cm} + K_D^{cm}}. \tag{44}$$

As only active ribosomes will contribute to the accumulation of biomass, we must rewrite the dynamics as

$$\frac{dM}{dt} = \gamma(tRNA^c) M_{Rb}^{\text{active}} = \gamma(tRNA^c) P_{\text{active}} M_{Rb}. \tag{45}$$

To make the predictions shown in Fig. 3(E and F) of the main text, we assumed that the chloramphenicol concentration in the growth medium is equal to the intracellular concentration and take $K_D^{cm} \approx 0.5$ nM.

#### Incorporating effects of excess protein stress

We consider that the excess protein synthesis can be modeled as the introduction of a new protein class, which we consider to have a total mass of $M_X$. Following the allocation parameters of the flux-parity model as defined in Eq. 32, we can introduce a new allocation parameter $\phi_X$ such that

$$\frac{dM_X}{dt} = \phi_X \frac{dM}{dt} \;;\; \phi_O + \phi_{Mb} + \phi_{Rb} + \phi_X = 1. \tag{46}$$

In the right-hand panel of Fig. 5B in the main text, we show that a collection of data can be collapsed onto a single line that relates the relative change in growth rate as a function of the excess protein that is synthesized. While we cannot fully solve the flux-parity model analytically, we can derive an analytical expression of this relation. Specifically, we note that the steady-state growth rate in the absence of excess expression $\lambda$ follows the simple relation

$$\lambda = \gamma(tRNA^c)\phi_{Rb}\left(\frac{tRNA^c}{tRNA^u}\right) = \gamma_{max}(1-\phi_O)\frac{tRNA^c}{tRNA^c + K_M^{tRNA^c}}\frac{\frac{tRNA^c}{tRNA^u}}{\frac{tRNA^c}{tRNA^u}+\tau}. \tag{47}$$

This can be easily extended to compute the growth rate under excess protein synthesis $\lambda_X$ as

$$\lambda_x = \gamma(tRNA^c)\phi_{Rb}\left(\frac{tRNA^c}{tRNA^u}\right) = \gamma_{max}(1-\phi_O-\phi_X)\frac{tRNA^c}{tRNA^c + K_M^{tRNA^c}}\frac{\frac{tRNA^c}{tRNA^u}}{\frac{tRNA^c}{tRNA^u}+\tau}. \tag{48}$$

We can take the ratio of these growth rates to yield an expression for the collapse function

$$\frac{\lambda_X}{\lambda} = \frac{\gamma_{max}(1-\phi_O-\phi_X)\frac{tRNA^c}{tRNA^c + K_M^{tRNA^c}}\frac{\frac{tRNA^c}{tRNA^u}}{\frac{tRNA^c}{tRNA^u}+\tau}}{\gamma_{max}(1-\phi_O)\frac{tRNA^c}{tRNA^c + K_M^{tRNA^c}}\frac{\frac{tRNA^c}{tRNA^u}}{\frac{tRNA^c}{tRNA^u}+\tau}}. \tag{49}$$

If we assume that the excess protein synthesis affects *only* $\phi_X$, leaving all other parameters untouched, Eq. 49 reduces to the concise form

$$\frac{\lambda_X}{\lambda} = \frac{1-\phi_O-\phi_X}{1-\phi_O}, \tag{50}$$

which is the linear relation plotted in the right-hand side of Fig. 5B of the main text.

Aside from the collapse, we also show how the flux-parity model quantitatively predicts the growth rate as a function of excess protein for three different media (Fig. 5C, center panel). In this case, we require some knowledge of what the metabolic rate $\nu_{max}$ is for those specific conditions. As the metabolic rate is an efficient rate incorporating the action of different metabolic reactions and serving as a proxy of the nutrient quality, it is not possible to make an *a priori* estimate of its value. To nevertheless estimate $\nu_{max}$ for each condition, we determined its value by using the simple allocation model as encoded in Sec. , assuming the growth rate $\lambda$ and the ribosomal content describes the allocation towards ribosomes $\phi_{Rb}$. Under the simple allocation model, we note that an expression for the metabolic rate can be solved from the steady-state precursor concentration $c_{pc}^*$ (Eq. 18) to yield

$$\nu_{max} = \frac{\lambda\left(c_{pc}+1\right)}{1-\phi_O-\phi_{Rb}}. \tag{51}$$

The steady-state precursor concentration $c_{pc}^*$ can be solved from the definition of the steady-state growth rate and has the form

$$c_{pc}^* = \frac{K_D^{c_{pc}}\lambda}{\phi_{Rb}\gamma_{max}\left(1-\frac{\lambda}{\phi_{Rb}}\right)}. \tag{52}$$

Combining Eqs. 51 and 52 yields an expression for the maximal metabolic rate $\nu_{max}$,

$$\nu_{max} = \frac{\lambda}{1-\phi_O-\phi_{Rb}}\left(\frac{K_D^{c_{pc}}\lambda}{\phi_{Rb}\gamma_{max}\left(1-\frac{\lambda}{\phi_{Rb}}\right)}+1\right). \tag{53}$$

Thus, given knowledge of the steady-state growth rate $\lambda$ and the allocation towards ribosomes $\phi Rb$ (which are both measured quantities), the value of $\nu_{max}$ can be derived.

##### Incorporating effects of nutrient upshifts

To model the dynamics of growth in fluctuating conditions, we asserted that a nutritional upshift is equivalent to an instantaneous change in the metabolic rate such that $\nu_{max}^{preshift} < \nu_{max}^{postshift}$. However, this is not completely sufficient to capture the phenomenology that is observed. It is becoming exceedingly clear that bacterial cells are non-optimal in *what* genes they express, with many proteins that are synthesized are ultimately useless in the specific condition [64]. This can have very important effects on the growth rate as any amount of conditionally useless protein that's synthesized consumes resources that could otherwise be partitioned to the proteins that need to be synthesized. To incorporate this effect, we introduce another protein class with an allocation parameter $\phi_{ø}$. As the degree of conditionally useless expression is significantly more pronounced in slow rather than fast conditions [27, 33, 64], we further asserted that the magnitude of this sector also changed in response to the nutritional upshift such that $\phi_{ø}^{preshift} > \phi_{ø}^{postshift}$. The precise value of this sector is less important than the difference in the pre- and post-shift condition and can be considered as an additional rescaling factor as described in Appendix 5. Thus, for all nutritional shifts in this work, we considered that $\phi_{ø}^{postshift} = 0$ and the value of $\phi_{ø}^{preshift}$ to be linearly proportional to the difference in the growth rates between the pre- and post-shift conditions.

##### Incorporating effects of nutrient depletion

Up to this point, we have explored the flux-parity model under the assumption that the nutrients in the environment were saturating, such that $\nu(c_{nt}) \approx \nu_{max}$. However, a dependence on the environmental nutrient concentration $c_{nt}$ can be easily included in the definition of the metabolic rate $\nu$ as

$$\nu(tRNA, c_{nt}) = \nu_{max}\left(\frac{tRNA^u}{tRNA^u + K_M^{tRNA^u}}\right)\left(\frac{c_{nt}}{c_{nt} + K_M^{c_{nt}}}\right), \tag{54}$$

where $K_M^{c_{nt}}$ is the Michaelis-Menten constant. We can then model the dynamics of the nutrient concentration $c_{nt}$ in a batch-culture system as

$$\frac{dc_{nt}}{dt} = -\frac{\nu(tRNA, c_{nt})M_{Mb}}{Y}, \tag{55}$$

where $Y$ is the yield coefficient.

##### Data Sets

This work leverages a large collection of data, primarily from *E. coli*, to evaluate the accuracy of our model in describing biological phenomena. These data come from a range of studies spanning around 50 years of measurements from different groups and different geographical locations. Collecting and curating this large data set required the manual transcribing of data from papers as well as various standardization steps to ensure that measurements were truly comparable between studies, as is outlined in Table S2.

#### Data and Code Availability

This work is accompanied by a website (`cremerlab.github.io/flux_parity`) which houses a suite of interactive figures to help the reader gain an intuition for the way the theory operates. Data sets can be downloaded individually from this website. Alternatively, all data and Python code used in this work can be cloned as a GitHub repository (`github.com/cremerlab/flux_parity`). This repository and its releases are also available via Zendodo with the DOI: 10.5281/zenodo.5893800.

#### Acknowledgments

We thank Uri Alon, Markus Arnoldini, Rachel Banks, Suzy Beeler, Nathan Belliveau, Terence Hwa, Soichi Hirokawa, Christine Jacobs-Wagner, Sergey Kryazhimskiy, Armita Nourmohammad, Manuel Razo-Mejia, Tom Röschinger,

Jan Skotheim, Cat Triandafillou, and members of the Tadashi Fukami, Dmitri Petrov, Alfred Spormann, and JC research groups for extensive discussions and the critical reading of the manuscript. GC acknowledges support by the NSF Postdoctoral Research Fellowships in Biology Program (Grant No. 2010807).

[31] Rohan Balakrishnan, Matteo Mori, Igor Segota, Zhongge Zhang, Ruedi Aebersold, Christina Ludwig, and Terence Hwa. Principles of gene regulation quantitatively connect DNA to RNA and proteins in bacteria. *bioRxiv*, page 2021.05.24.445329, May 2021.

[32] Matteo Mori, Zhongge Zhang, Amir Banaei-Esfahani, Jean-Benoît Lalanne, Hiroyuki Okano, Ben C Collins, Alexander Schmidt, Olga T Schubert, Deok-Sun Lee, Gene-Wei Li, Ruedi Aebersold, Terence Hwa, and Christina Ludwig. From coarse to fine: The absolute *Escherichia coli* proteome under diverse growth conditions. *Molecular Systems Biology*, 17(5), May 2021.

[33] Nathan M. Belliveau, Griffin Chure, Christina L. Hueschen, Hernan G. Garcia, Jane Kondev, Daniel S. Fisher, Julie A. Theriot, and Rob Phillips. Fundamental limits on the rate of bacterial growth and their influence on proteomic composition. *Cell Systems*, July 2021.

[34] Eyal Metzl-Raz, Moshe Kafri, Gilad Yaakov, Ilya Soifer, Yonat Gurvich, and Naama Barkai. Principles of cellular resource allocation revealed by condition-dependent proteome profiling. *eLife*, 6:e28034, August 2017.

[35] Joao A. Paulo, Jeremy D. O'Connell, Aleksandr Gaun, and Steven P. Gygi. Proteome-wide quantitative multiplexed profiling of protein expression: Carbon-source dependency in *Saccharomyces cerevisiae*. *Molecular Biology of the Cell*, 26(22):4063–4074, November 2015.

[36] Joao A. Paulo, Jeremy D. O'Connell, Robert A. Everley, Jonathon O'Brien, Micah A. Gygi, and Steven P. Gygi. Quantitative mass spectrometry-based multiplexing compares the abundance of 5000 S. *cerevisiae* proteins across 10 carbon sources. *Journal of Proteomics*, 148:85–93, October 2016.

[37] Jianye Xia, Benjamin Sánchez, Yu Chen, Kate Campbell, Sergo Kasvandik, and Jens Nielsen. Proteome allocations change linearly with specific growth rate of *Saccharomyces cerevisiae* under glucose-limitation. Preprint, In Review, May 2021.

[38] Michael Jahn, Vital Vialas, Jan Karlsen, Gianluca Maddalo, Fredrik Edfors, Björn Forsström, Mathias Uhlén, Lukas Käll, and Elton P. Hudson. Growth of Cyanobacteria Is Constrained by the Abundance of Light and Carbon Assimilation Proteins. *Cell Reports*, 25(2):478–486.e8, October 2018.

[39] Xiongfeng Dai, Manlu Zhu, Mya Warren, Rohan Balakrishnan, Vadim Patsalo, Hiroyuki Okano, James R. Williamson, Kurt Fredrick, Yi-Ping Wang, and Terence Hwa. Reduction of translating ribosomes enables *Escherichia* coli to maintain elongation rates during slow growth. *Nature Microbiology*, 2(2):1–9, December 2016.

[40] Markus Basan, Manlu Zhu, Xiongfeng Dai, Mya Warren, Daniel Sévin, Yi-Ping Wang, and Terence Hwa. Inflating bacterial cells by increased protein synthesis. *Molecular Systems Biology*, 11(10):836, October 2015.

[41] Conghui You, Hiroyuki Okano, Sheng Hui, Zhongge Zhang, Minsu Kim, Carl W. Gunderson, Yi-Ping Wang, Peter Lenz, Dalai Yan, and Terence Hwa. Coordination of bacterial proteome with metabolism by cyclic AMP signalling. *Nature*, 500(7462), August 2013.

[42] Chenhao Wu, Rohan Balakrishnan, Matteo Mori, Gabriel Manzanarez, Zhongge Zhang, and Terence Hwa. Cellular perception of growth rate and the mechanistic origin of bacterial growth laws. Preprint, Systems Biology, October 2021.

[62] Anjan Roy, Dotan Goberman, and Rami Pugatch. Transcription-translation machinery – an autocatalytic network coupling all cellular cycles and generating a plethora of growth laws. *arXiv:2103.15356 [physics, q-bio]*, March 2021.

[74] Ludovico Calabrese, Jacopo Grilli, Matteo Osella, Christopher P. Kempes, Marco Cosentino Lagomarsino, and Luca Ciandrini. Role of protein degradation in growth laws. Technical report, August 2021. Company: Cold Spring Harbor Laboratory Distributor: Cold Spring Harbor Laboratory Label: Cold Spring Harbor Laboratory Section: New Results Type: article.

[86] John S. Favate, Shun Liang, Srujana S. Yadavalli, and Premal Shah. The landscape of transcriptional and translational changes over 22 years of bacterial adaptation. *bioRxiv*, page 2021.01.12.426406, January 2021. Publisher: Cold Spring Harbor Laboratory Section: New Results.

### Supplementary Figures

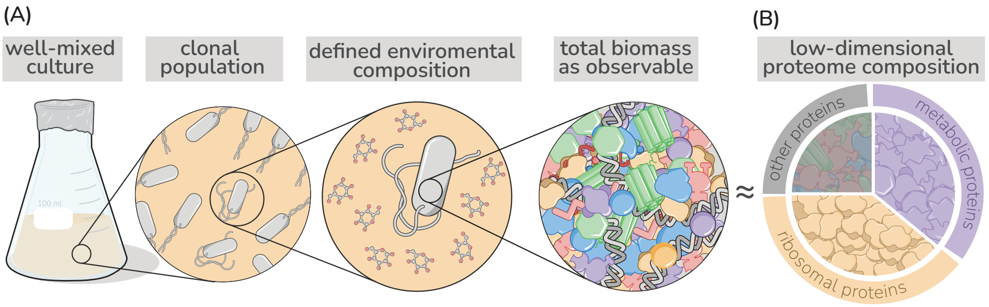


**Supplementary Figure 1: Coarse grained description of biomass and the proteome** Low-dimensional allocation models consider in their simplest form growth of a clonal population within a well-mixed environment. Biomass is described in a highly simplified manner focusing in the simplest case on protein synthesis alone [50, 52], with different protein species jointly considered by a few different protein classes. Here, metabolic proteins (purple), ribosomal proteins (gold), or "other" proteins (gray).

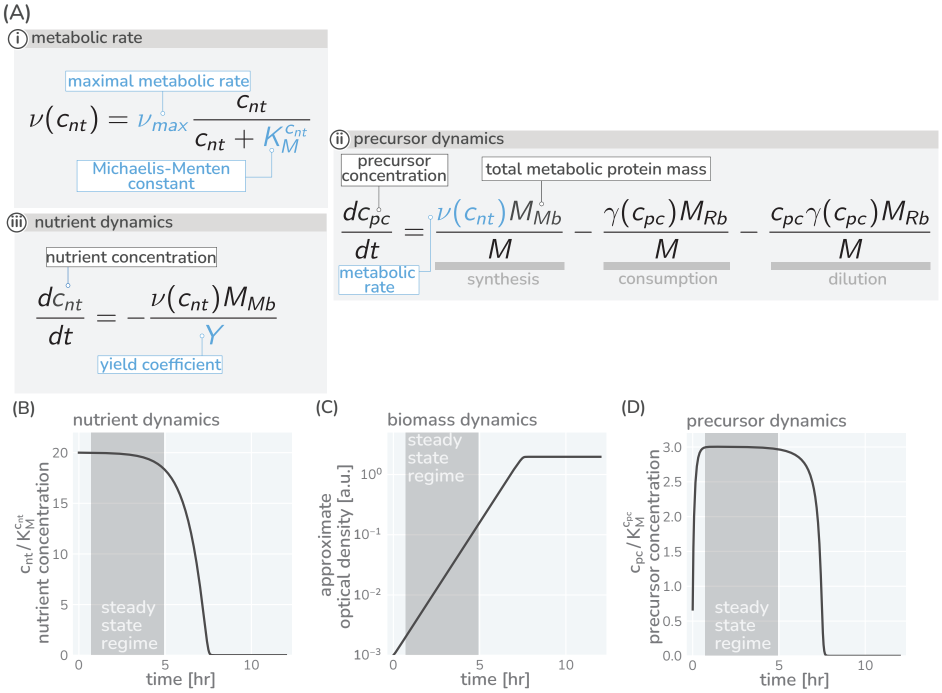


**Supplementary Figure 2: Precursor synthesis and growth when nutrients are not saturating.** In general, the environmental conditions microbes encounter changes rapidly and nutrient availability commonly limits growth. Growing batch cultures, fox example run out of nutrients eventually and growth stops. The allocation modeling framework can account such a dynamics by including metabolic rates which depend on the nutrient concentrations in the environment. In the simplest case, one nutrient source is considered (concentration $n$) with the metabolic rate $\nu(n)$ depending on the concentration in a Michaelis-Menten manner with a maximal metabolic rate being reached only at high nutrient concentrations (A,i). The dynamics of precursors is given by a balance of synthesis, consumption, and dilution (A,ii), replacing the corresponding equation of the simple model in [Fig. 1(B,iv)]. The modeling of growth further requires the explicit modeling of nutrient concentrations. This dynamics depends on the specifics of the environment and, depending on the environment, can become very complex with multiple sources and sinks affecting the nutrient concentration. Here, we consider a typical batch culture scenario in which cells grow under well-mixed conditions. Nutrients are provided only initially and nutrient concentrations are falling because of consumption (A,iii). (B-D) Resulting temporal variation of nutrient concentrations (C), biomass accumulation (D), and precursor concentration (E) when integrating the model equations and using a parameter set descriptive of *E. coli* growing in a glucose-minimal medium with a growth rate $\approx 1$ hr$^{-1}$ and a starting glucose concentration of $10mM$. As experimentally observed, initially abundant nutrients are consumed and biomass accumulates (exponential phase) until nutrients are exhausted and growth stops (saturation phase) (B and C). Importantly, precursor concentrations (D) quickly reach a constant plateau which lasts until nutrients become scare ($c_{nt} \gg K_M^{c_{nt}}$ and $\nu(c_{nt}) \approx \nu_{max}$). During this transient period (shaded regions) the synthesis of precursors matches the consumption by protein synthesis and dilution, meaning $\frac{dc_{pc}}{dt} = 0$. Given a constant precursor concentration $c_{pc}^*$, the translation rate $\gamma(c_{pc}^*)$ is also constant. As a consequence, the protein pool approaches a steady composition dictated by the allocation parameters ($\frac{M_{Rb}}{M} = \phi_{Rb}$, $\frac{M_{Mb}}{M} = \phi_{Mb}$ and $\frac{M_O}{M} = \phi_O$). With precursor concentrations and protein composition remaining constant, the system is in a *steady-state* and biomass accumulates exponentially over time, $\frac{dM}{dt} = \gamma(c_{pc}^*)\phi_{Rb}M \equiv \lambda M$. This is the steady state regime we focus on in the main text. Note that the steady state growth regime readily emerges when we consider dilution (see Appendix 3). Model parameters are provided in Table S1. Biomass units are converted to optical density assuming at $OD_{600nm} = 1$, there are $10^9$ cells per mL and $10^9$ amino acids per cell. An interactive version of these dynamics can be found on the paper website.

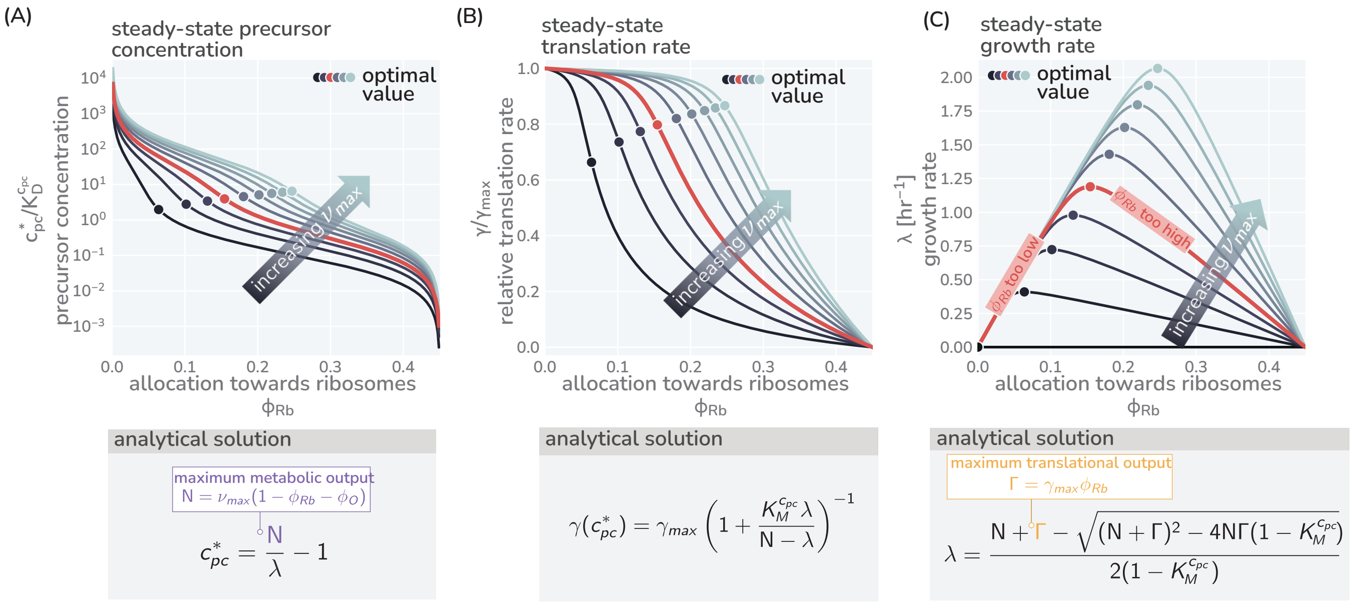


**Supplementary Figure 3: Modeling predictions of steady growth behavior**. (A) Variation of the precursor concentration with varying allocation parameters ($\phi_R$) and maximal metabolic rate ($\nu_{max}$). (B C) Corresponding trends of translation and growth rate as also shown in Fig. 1 (C) and (D). Corresponding boxes show the analytical expression describing the steady-state precursor concentration, translation speed, and growth rate with details of the derivation provided in Appendix 7. Used model parameters provided in Table S1. Colors indicate different metabolic rates $\nu_{max}$= 0.2 - 12.5 hr$^{-1}$.

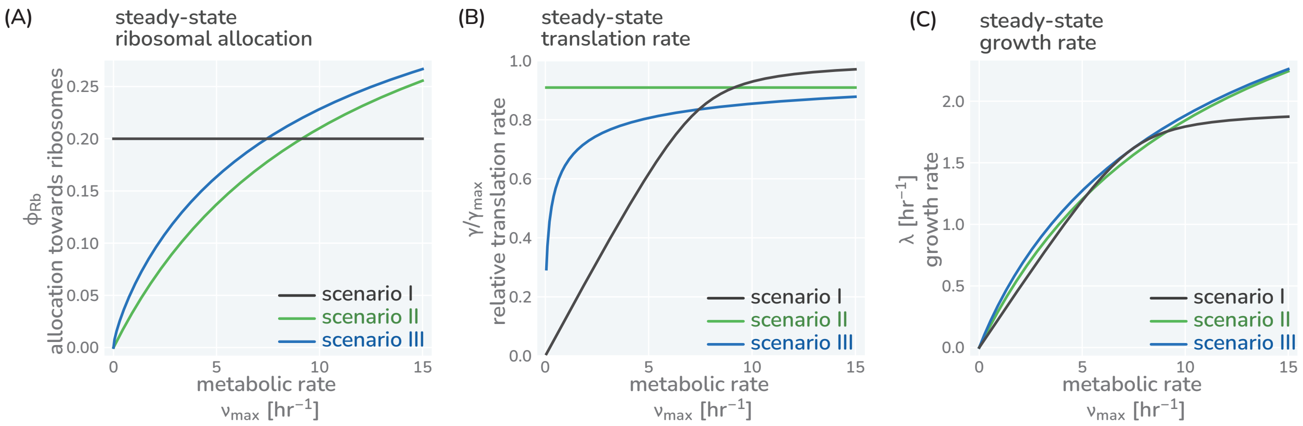


**Supplementary Figure 4: Three different allocation scenarios**. The variation of precursor concentration (A), translation rate (B), and growth rate (C) with changing metabolic rate is shown for the three allocation scenarios introduced in the main text; fixed allocation (scenario I, black lines), prioritizing fast translation (scenario II, green lines), and growth-optimal allocation (scenario III, blue lines). Plotted are the analytical solutions provided in Fig. 1(E) and derived in the Methods. The resulting relations between growth rate and translation as well as growth rate and ribosome content are shown in Figure 1(G,H). We here discuss the consequence of these allocation scenarios in more detail. *Scenario I - fixed allocation*: In this scenario, allocation is fixed and does not vary with conditions. Locking in the ribosome allocation to $\phi_{Rb} = 0.25$ (A, black line), for example, carries strong consequences for translation and growth rates (B and C, black lines). When conditions are poor ($\nu_{max}$ is small), the translation rate is significantly lower than the maximal rate as there are too many ribosomes competing for a small pool of precursors (B). The translation and growth rates increase with the metabolic rate $\nu_{max}$ until the influx of precursors is sufficiently high such that all ribosomes are translating close to their maximum and growth-rate is at its optimal value. Further increasing the metabolic rate does not increase the growth rate [plateau of black curve in (B)] as all ribosomes are already translating close to their maximum rate. *Scenario II - prioritizing fast translation*: In this scenario, allocation is adjusted such that translation rates are maintained at a high value. This is achieved by tuning the allocation between ribosomes and metabolic proteins such that a constant precursor concentration $c_{pc}^* \gg K_M^{c_{pc}}$ is maintained. For example, at higher metabolic rates, the metabolic proteins can sustain a higher influx of precursors allowing a larger allocation towards ribosomal proteins $\phi_R$ (green lines). *Scenario III - optimizing growth*: In this scenario, allocation is tuned to optimize growth rate across conditions, meaning that the fastest growth rate is achieved given a set metabolic rate and other model parameters. For example, the allocation towards ribosomes $\phi_{Rb}$ is adjusted with the metabolic rate such that the growth rate rests at the peak of the curves shown in Fig. 1(C) (blue lines). Accordingly, the growth rate continues to increase with higher metabolic rates always exceeding the growth rate of scenario I and II (C, green line)]. Model parameters follow the reference set for *E. coli* (Table S1). Black lines correspond to a constant allocation $\phi_{Rb}^{(I)} = 0.20$ and green lines correspond to a constant precursor concentration $c_{pc}^* \approx 10K_M^{c_{pc}}$, yielding a constant translation rate of $\gamma(c_{pc}^*) \approx 0.9\gamma_{max}$. An interactive version of these figure panels is available on the paper website.

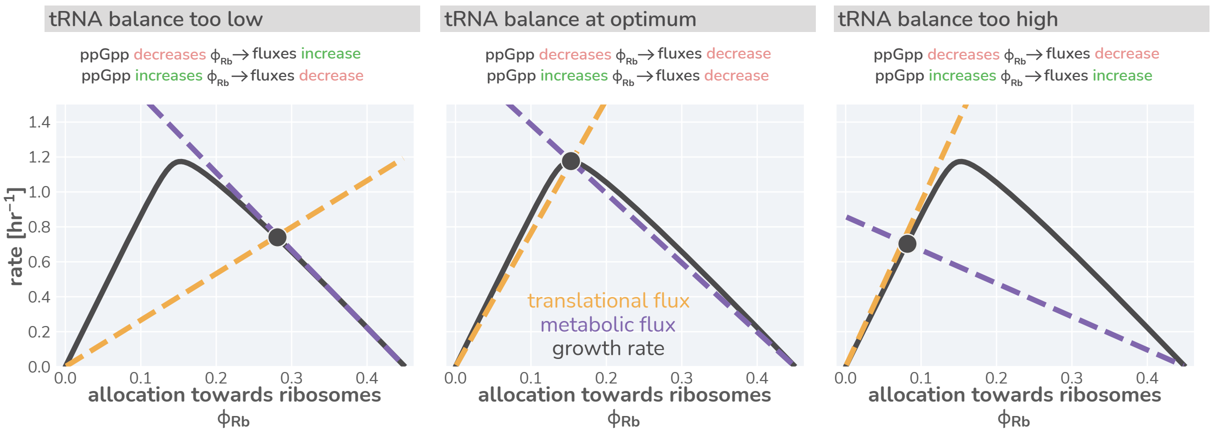


**Supplementary Figure 5: Flux-parity directs allocation parameters towards an optimum.** Black lines represent the steady-state growth rate as a function of the allocation towards ribosomes $\phi_{Rb}$. Dashed gold and purple lines correspond to the translational and metabolic fluxes, with their intersection indicating the steady-state. The different panels consider from left to right three scenarios with a too low, optimal, and too high allocation towards ribosomes. An interactive version of this figure is available on the paper website

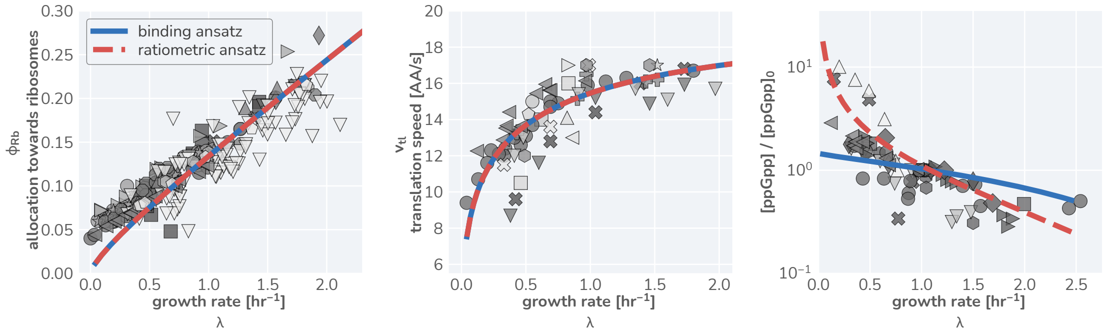


**Supplementary Figure 6: Comparison of predictive capacity of flux-parity allocation between ppGpp scaling ansatzes.** Measurements are shown for ribosomal content, translation rate, and relative ppGpp concentration from left to right, respectively. Markers are the same as those in Fig. 3 of the main text. Solid blue line shows predicted steady-state behavior assuming a simple ansatz of ribosome-tRNA binding probabilities (Eq. 41). Dashed red line denotes predicted behavior using the ansatz that ppGpp concentration is dependent on the charging balance.
